# Effects of HSP70 chaperones Ssa1 and Ssa2 on Ste5 scaffold and the mating mitogen-activated protein kinase (MAPK) Pathway in *Saccharomyces cerevisiae*

**DOI:** 10.1101/2022.08.19.503794

**Authors:** Francis W. Farley, Ryan R. McCully, Paul B. Maslo, Lu Yu, Mark A. Sheff, Homayoun Sadeghi, Elaine A. Elion

## Abstract

Ste5 is a prototype of scaffold proteins that regulate activation of mitogen-activated protein kinase (MAPK) cascades in all eukaryotes. Ste5 associates with many proteins including Gβγ (Ste4), Ste11 MAPKKK, Ste7 MAPKK, Fus3 and Kss1 MAPKs, Bem1, Cdc24. Here we show that Ste5 also associates with heat shock protein 70 chaperone (Hsp70) Ssa1 and that Ssa1 and its ortholog Ssa2 are together important for Ste5 function and efficient mating responses. The majority of purified overexpressed Ste5 associates with Ssa1. Loss of Ssa1 and Ssa2 has deleterious effects on Ste5 abundance, integrity, and localization particularly when Ste5 is expressed at native levels. The status of Ssa1 and Ssa2 influences Ste5 electrophoresis mobility and formation of high molecular weight species thought to be phosphorylated, ubiquitinylated and aggregated and lower molecular weight fragments. A Ste5 VWA domain mutant with greater propensity to form punctate foci has reduced predicted propensity to bind Ssa1 near the mutation sites and forms more punctate foci when Ssa1 Is overexpressed, supporting a dynamic protein quality control relationship between Ste5 and Ssa1. Loss of Ssa1 and Ssa2 reduces activation of Fus3 and Kss1 MAPKs and *FUS1* gene expression and impairs mating shmoo morphogenesis. Surprisingly, *ssa1*, *ssa2*, *ssa3* and *ssa4* single, double and triple mutants can still mate, suggesting compensatory mechanisms exist for folding. Additional analysis suggests Ssa1 is the major Hsp70 chaperone for the mating and invasive growth pathways and reveals several chaperone-network proteins required for mating morphogenesis.

## Introduction

The mating pathway in *S. cerevisiae* involves differentiative events that include cell cycle arrest in G1 phase, specialized polarized growth towards a pheromone gradient, cell-cell attachment and fusion to form a zygote that reenters the cell cycle (Fig1 A)(1–7). These events are mediated by a conserved receptor-G protein-coupled MAPK cascade that activates MAPKs Fus3 and Kss1, with Fus3 being more critical (5–7). Ste5, the first described MAPK scaffold/tether (8), has multiple functions in the mating pathway and is essential for activation of Fus3 and Kss1 (9–12). During mating in a cells, α factor pheromone binds to dimers of Ste2 (a type D GPCR) (13), which liberate bound Gβ-Ste4/Gγ-Ste18 dimers from Gα-Gpa1 subunits (7, 8). Ste4/Ste18 then binds to and recruits a Cdc42-activated form of PAK Ste20, a MAPKKKK. Ste5 also binds the liberated Gβ/Ste4-GγSte18 dimers and four protein kinases, MAPKKK Ste11, MAPKK Ste7 and MAPK Fus3 and MAPK Kss1, with MAPKK Ste7 bridging and enhancing MAPK association (Fig 1A, Fig S1A (1–13)). Ste5 orchestrates Ste20 activation of Ste11, Ste11 activation of Ste7, Ste7 activation of Fus3 and auto-activation of Fus3 (1,4,5–7,9,10,14) potentially through cis and trans kinase phosphorylation events on dimers or oligomers of Ste5 (12). These many functions are mediated through Ste4 interactions, conformational changes, oligomerization and interactions with Cln2/Cdc28, phosphatases, and the Bem1 complex that includes Cdc42 Rho-type GTPase and Cdc24 (Dbl guanine nucleotide exchange factor for Cdc42) (4-7, 15-19). Ste5 also regulates transcription in association with Fus3 (20, 21), and may have direct roles in cell polarization and cell fusion (1,3,6–7,22–25). Fus3 and Kss1 phosphorylate numerous targets required for mating that include transcription factors and repressors (e.g. Ste12, Dig1, Dig2 and others), the Far1 cyclin dependent kinase inhibitor, cytoskeletal regulators (e.g. Bni1) (1-7,20). Many of the mating pathway components are used in the invasive growth pathway which is activated by low pheromone concentrations, poor nutrients, and cell wall stress i (Fig. 1A)(26–29). Invasive growth (pseuodohyphal growth in diploids) is activated by plasma membrane mucins linked to Cdc42, Ste20, Ste11, Ste7 and Kss1. Kss1 regulates the same transcriptional repressors Dig1, Dig2 and transcription activator Ste12, in addition to Tec1 and other transcription factors which together promote invasive growth (Fig 1A). The mating and invasive growth/pseudohyphal pathways also regulate virulence of pathogenic yeast (e.g. *S. cerevisiae, C. glabrata*) that infect humans and crops (28, 29).

**FIG 1.**
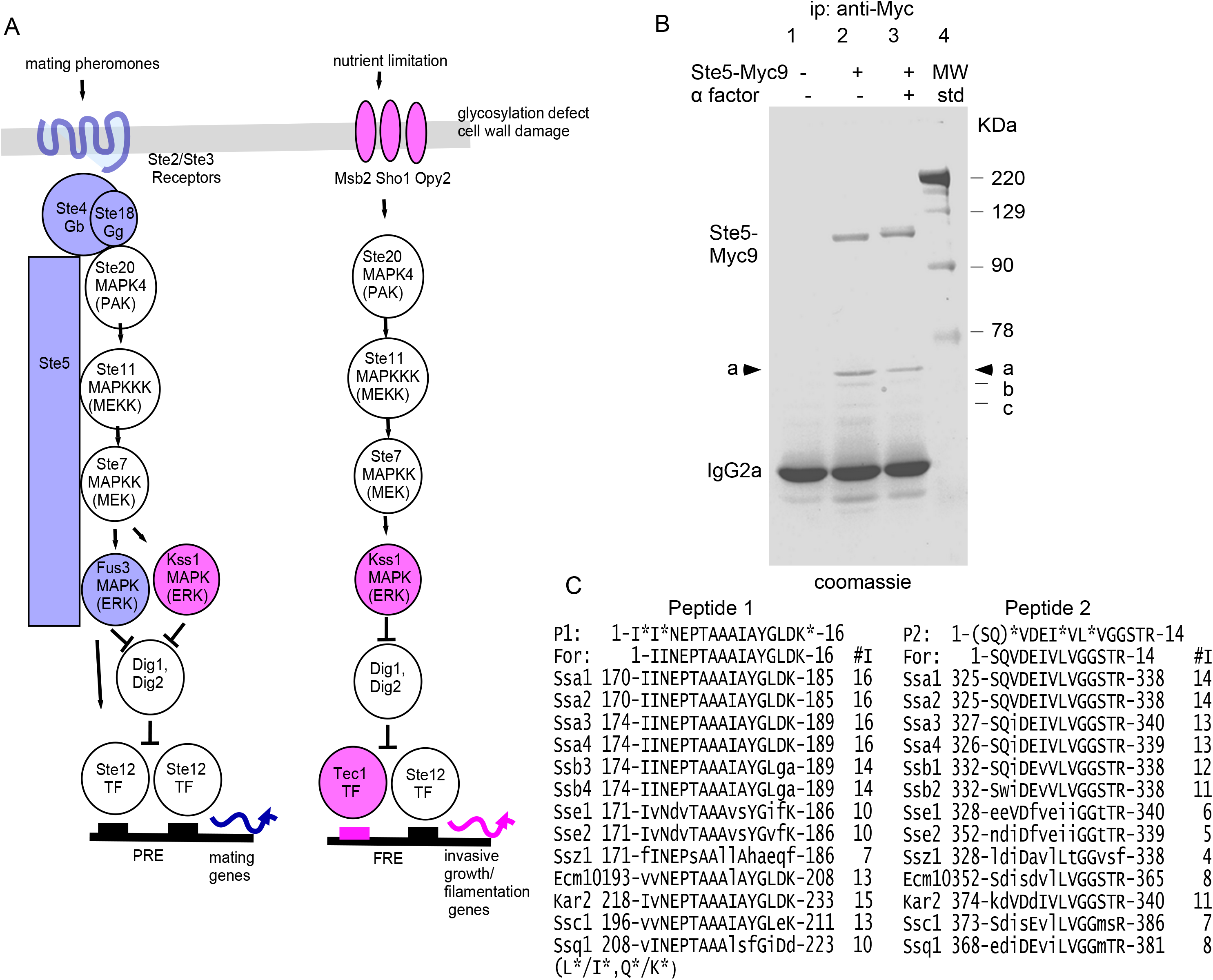
The majority of immunoprecipitated Ste5-Myc9 is bound to Hsp70 Ssa1 and possibly Ssa2. A. Schematics of mating pathway and invasive growth pathway MAPK cascades and Ste5 interactions. B. Ste5-Myc9 co-immunoprecipitates (co-IPs) a ∼70 KDa protein(s). Coomassie stain of an SDS-polyacrylamide gel of Ste5-Myc9 IP’d with 9E10 monoclonal antibody from yeast whole cell extracts prepared from mitotically dividing EY1775 (*MATa ste5 bar1*) cells expressing *STE5-MYC9-2μ* (pSKM19). Lane 1: EY957 (STE5) cells, lanes 2,3: EY1775 expressing STE5-MYC9 (pSKM19). “+” indicates treatment of cells with 50 nM ✓ factor for 2 hours. C. The 70kDa protein is most homologous to Ssa1 and Ssa2. Two peptide sequences from the 70 kDa band in A are aligned with *S. cerevisiae* HSP70 proteins. The “*” indicates residue uncertainty, *L/I and *Q/K.

Scaffold proteins play crucial roles in transmitting signals through MAPK cascades to downstream targets in eukaryotes (4,8,11–12,30–31). Ste5 is regulated at multiple levels to ensure differentiation occurs at the right time and place. Tight control of the conformation, localization and abundance of Ste5 regulates signaling specificity, proper timing and amplitude of MAPK activation and prevents deleterious misactivation of the mating pathway. These controls include assembly of Ste5 into an active dimer (and possibly higher order oligomers; 12,15,17), relief of autoinhibitory interactions between PH and VWA domains (7,12,16,17), membrane lipids (i.e. phosphatidylinositol-(4, 5)-bisphosphate and fatty acids) that bind Ste5 and enhance its interaction with Gβγ (7,32–33) and at least 16 phosphorylation events including overlapping phosphorylation by Fus3 and Cln/Cdc28 that influence Ste5 residency at the plasma membrane and abundance (34, https://phosphogrid.org/sites/32160). Ste5 is spatially controlled through nucleocytoplasmic shuttling which imports a nuclear pool for transcription, limits the amount of Ste5 available to bind to Gβγ and also exports Ste5 for binding to Gβγ during pheromone signaling (1, 35–37). During vegetative growth, high phosphorylation of Ste5 by Cln/CDK adds negative charges that disrupt membrane localization of Ste5 and reduces its stable binding to Gβγ (24, 34). The phosphorylation of Ste5 also triggers its ubiquitinylation and degradation through SCF^Cdc4^ and the proteasome in the nucleus (37). During pheromone signaling, Ste5 abundance increases through Fus3 and Kss1 (25), in part through inhibition of Cln/Cdk by Far1, a CDK inhibitor. Loss of Fus3 causes enhanced, depolarized accumulation of Ste5 at the plasma membrane (24) whereas loss of both Cdc28 and Fus3 restores polarized localization of Ste5 at the plasma membrane but also causes Ste5 to accumulate in vesicles that that are likely vacuolar (24), possibly because of loss of solubility derived from phosphorylation (e.g.). Here, we present evidence of novel requirements for Hsp70 chaperones Ssa1 and Ssa2 for Ste5 stability and integrity.

The heat shock 70 (Hsp70) proteins regulate evolutionarily conserved responses of all cells to proteotoxic stress (39–44). Their misregulation is associated with numerous diseases including cancer (45–47), Alzheimer’s Disease (48), Parkinson’s Disease (48), and fungal infections (29, 49–51). *S. cerevisiae* has 14 Hsp70 proteins of which 9 are cytosolic and nuclear. The “classic” cytosolic Hsp70s are Ssa1-Ssa4 in *S. cerevisiae* and Hsc70s in metazoans and mammals (51). Ssa1-Ssa4 function with J-domain proteins/Hsp40s (i.e. Ydj1, Sis1) and nucleotide exchange factors (e.g. Fes1/HBp1) (39-44,52) and bind substrates on surface exposed hydrophobic stretches with certain characteristics (39, 53). Ssa1 and Ssa2 are expressed constitutively through heat shock transcription factor 1 (HSF1, 54) and function at cool and ambient temperatures. Heat stress increases the level of Ssa2 but not that of Ssa1. Much of the heat shock response also relies on Msn2/Msn4 transcription factors which modulate membrane composition, carbohydrate flux, and cell wall integrity independently of Hsf1, Ssa1 and Ssa2 (60, 61).

The classic human and *S. cerevisiae* Hsp70 proteins (Ssa1, Ssa2, Ssa3, Ssa4) are together essential for viability and have major functions in folding nascent translated polypeptides, refolding of misfolded proteins and disaggregation of protein aggregates in cooperation with specific co-chaperones (39-44,57-58). They deliver proteins to translocation machinery at organellar membranes, and manage irretrievably damaged and misfolded proteins (including aggregated proteins, prions and RNA stress granules) by facilitating their degradation by the proteasome or by the lysosome or autophagy (39-41,59-60). Ssa1 and Ssa2 have many functional similarities and subtle functional differences (61**add). Major functions of Ssa1 and Ssa2 include assisting the ribosome-associated Hsp70s Ssb1 and Ssb2 to fold emerging polypeptides during translation (57–58) and transfer polypeptides to other chaperones such as Hsp90s for protein maturation (62). Ssa1 and Ssa2 help translocate proteins through endoplasmic reticulum and mitochondrial membranes (61, 62). They promote degradation of misfolded and damaged proteins and impact various types of inclusion bodies (63–67), and promote ER-associated degradation of ER proteins in cytosolic proteasomes (ERAD; (68)), proteasome degradation of ubiquitinylated and nonubiquinylated proteins (65–67), degradation in vacuoles and autophagosomes (39, 68–69). Ssa1 and Ssa2 also facilitate nuclear import (70, 71), assembly and disassembly of multi-protein complexes (e.g. prions (72), proteasomes (73), clathrin coats (74), and can be essential subunits of enzyme complexes (e.g. Ssa1/Ssa2/Rad9/Rad53 checkpoint complex (75)).

Despite a multitude of functions, obvious *ssa1* mutant phenotypes are rare. For example, whole cell extracts from a *ssa1* null mutant in one study had no obvious differences in individual proteins by 2d gel electrophoresis, (76). More obvious phenotypes have been seen when both *SSA1* and *SSA2* are deleted, although pulse labeling experiments indicate small (∼3%) global differences in protein degradation rates (77). The expression of *SSA3* and *SSA4* is elevated in an *ssa1D ssa2D* mutant, however, *SSA3* and *SSA4* can not substitute for *SSA1* and *SSA2* and neither can Hsf1 (40, 78–79).

Little is known about the role of Ssa proteins during mating. To date, only one mating pathway protein, Dig1, has been categorized as a candidate substrate for Ssa1 in cells treated with α factor mating pheromone, compared to 317 from vegetatively growing cells in the same study (80). Here we report the purification of Ssa1 with Ste5 during vegetative growth and α factor stimulation. Ssa1 is predominantly cytoplasmic but also accumulates in nuclei. Ssa1 and Ssa2 together enhance Ste5 abundance, integrity and localization in nucleus and at plasma membrane. Ssa1 and Ssa2 prevent degradation of Ste5 at specific proteolytic sites, especially as temperatures rise, and they influence antibody recognition of N- and C-terminal tags on Ste5. By contrast, Ssa1 and Ssa2 may downregulate Fus3 and have no obvious effect on the level of Tcm1, Ste7 or Kss1 by immunoblot analysis. Overexpression of Ssa1 and loss of Ssa1 and Ssa2 influences the localization of wild type and mutant forms of Ste5. Ssa1 and Ssa2 are required for efficient activation of the mating MAPK cascade, *FUS1* expression, and shmoo morphogenesis. Despite these many effects, loss of Ssa1 and Ssa2 does not block mating, suggesting strong compensatory mechanisms exist to tolerate reduced protein quality control. Analysis of other Hsp70 network mutants supports the existence of compensatory mechanisms.

## Materials and Methods

### Yeast strains and media

See Table S1 for a list of the yeast strains and plasmids used in this study made, propogated and stored following standard procedures (81, 82). Null deletion alleles of *ssa1, ssa2, ssa3 and ssa4* were used for all analyses. If no allele is specified, then it is a null deletion allele (indicated as *ste5D*). *S. cerevisiae* and *E. coli* strains were grown in standard selective media (termed SC) and yeast extract peptone (YEP) media following standard protocols (81, 82). Cells with *GAL1prom*-driven genes were pre-grown in media containing 2% raffinose before induction in medium containing 2% galactose. Previously constructed *ste5* mutant proteins (16,23,35–36) screened for aberrant localization or aggregates with or without excess Ssa1 included Ste5K49,50A-Myc9 (EY3398), Ste5K64-66A-Myc9 (EY3399), Ste5K49,50,64-66A-Myc9 (EY3400), Ste5(1-242)-GFP2 (EY3401), Ste5(1-242D49-66)-GFP2 (EY3402), ste5L482/485A-Myc9 (pYMW37/aka ste5NES1mII/EYL2778/YMY533), ste5D522-527-Myc9 (pYMW5/ aka steNES2D/ EYL2779/YMY534), ste5L633/636A-Myc9 (pYMW15/ aka ste5NES3mII/ EYL2780/YMY535), ste5L610A/614A/634A/637A-Myc9 (pYMW109/ aka ste5NES4mIINES3mII/ EYL2781/YMY536). Yeast and bacterial transformations were performed as described (83–84) except for temperature sensitive strains which had the following protocol modifications: strains were not heat shocked, more DNA was used for transformation, and strains were recovered for 20 minutes in rich medium before pelleting, and resuspending in selective medium for plating. Some *ssa* mutant strains were generated by sporulation of a *MATa/MATa ssa1::HIS3/SSA1 ssa2D::LEU2/SSA2 SSA3/ssa3::URA3 SSA4/ssa4::TRP1 ura3-52/ura3-52 his3/his3 leu2-3,112/leu2-3,112, trp1-1/trp1-1 his3/his3* diploid, (kind gift of E. Craig, University of Wisconsin) followed by phenotypic analysis of ascospore segregants. Backcrossing was done to make strains more congenic. Resistance to 5-fluoro-orotic acid and subsequent complementation tests and confirmation of mitochondrial function was done to isolate *ura3* mutant derivatives that revert at very low frequency for several *ssa* mutant combinations to permit introduction of plasmids harboring a *URA3* gene for selection. Mating type was determined by a qualitative patch mating assay (85). Strains were grown at room temperature and 30°C unless noted otherwise. Colony size was measured by enlarging the image of the petri plate to a diameter of 240 mm, measuring 50 colonies per strain and determining mean value with standard deviation.

### Antibodies

Antibodies were from the following sources: Mouse monoclonal 9E10 and mouse monoclonal 12CA5 (Harvard University antibody facility), anti-active phosphorylated p42p44 MAPK rabbit polyclonal antibody (Cell Signaling Technology Phospho-p44/42 MAPK(Erk1/2) (Thr202/Tyr204) #9101), anti-Fus3 peptide rabbit polyclonals (14, 20), anti-Kss1 peptide rabbit polyclonal (6775-kss1-yc-19, Santa Cruz Biotechnology, Inc.), monoclonal Tcm1 mouse monoclonal antibody (gift of J. Warner, Albert Einstein College of Medicine). The anti-phospho-p42p44 antibody recognizes the TpEYpWYRAPE motif and was made against a peptide from ERK2 that overlaps human ERK1. ERK1 residues 237-264 and ERK2 residues 220-247 are 92.9% identical (26/28 residues identical) and have homology to Fus3 residues 167-192 (15/28 identical) and Kss1 residues 170-195 (16/28 identical).

### Whole cell extracts, co-immunoprecipitation and immunoblot analysis

Cells were grown to an A_600_ of 0.8-1 and induced with α factor for 15 minutes where indicated, then pelleted by centrifugation, washed once with ice-cold water, then frozen in dry ice/ethanol. Pellets were thawed on ice and resuspended in 1.0 ml of modified H-buffer with sodium chloride and a variety of inhibitors of phosphatases and proteases (i.e. 10% glycerol, 25 mM Tris-HCl pH 7.4, 150 mM NaCl (in some experiments 200-250 mM NaCl), 15 μM MgCl_2_, 2mM EDTA, 15 mM EGTA, 1 mM DTT, 50 mM NaF, 0.1% Triton-X-100, 1 mM sodium azide, 0.025 each of mM meta- and ortho-sodium vanadate (pH7 stock solution), ∼0.5 mg/ml phenylmethylsufonyl fluoride (PMSF, Sigma), 5 mg/ml pepstatin-A (Sigma), chymostatin (5mg/ml, Sigma), 1 mM benzamidine (Sigma). In Figure 3B and Supplemental Figure 3 the extraction buffer also contained phosphatase inhibitor cocktail 1 (Sigma P2850 microcystin LR, cantharidin and (-)-p-bromotetramisole). In all Ste5-Myc9 experiments except in Figures 1 and 2, we also added 50 μM proteasome inhibitor [Z-Ile-Glu(OBut)-Ala-Leu-H (aldehyde) (Peptide International, Catalogue Number IAT-3169-v). Cell extracts were prepared by glass-bead breakage as described (16,20, 86). Co-IPs were done in the same buffer adjusted to 200 mM NaCl, and immunoblot analyses were carried out as described (16,20,23,86). The protein concentration of whole cell extracts was determined with the BioRad assay. For detection of Ste7-MYC, 1 mg of *CYC1prom-STE7-MYC* whole cell extracts was concentrated by 40% ammonium precipitation then dissolved in loading buffer prior to immunoblot analysis as described (85). Co-IPs used 250 μg to 2 mg of whole cell protein extracts, 15 μg of 12CA5, 1-2 μg of 9E10, and 30 μl of protein A agarose beads (Sigma). Samples were subjected to SDS-PAGE and immunoblot analysis as described (16,20,23,82,86–87). For analysis of Ste5-Myc9 phosphorylation, samples were run on native polyacrylamide gels that were 8% acrylamide, 0.19% bis-acrylamide, 0.75mM gel thickness and run at 40 mAmps. Benchmark prestained protein ladder (Life Technologies Cat. No.10748-010) was used for many of the gels as indicated in lab notebooks and in the figures. We note that several of the original films were accidentally labeled incorrectly with wrong protein ladder sizes taken from the catalog rather than the Benchmark package insert. The correct package insert molecular weights are in the figures. Immunoreactivity in immunoblots was detected with horseradish peroxidase-conjugated secondary antibody with ECL (Amersham, Arlington Heights, IL). Additional details of extraction methods in Fig 2 are in Supporting Information. The apparent molecular weight of protein bands in gels was determined by measuring the distance of migration of molecular weight markers (Rf) in the same gel, creating a graph of the log10 of the molecular weight of marker proteins versus their Rfs, then extrapolating from the graph a log10 for an Rf of a specific band and then taking the antilog of this value for the band’s apparent molecular weight.

**Figure 2.**
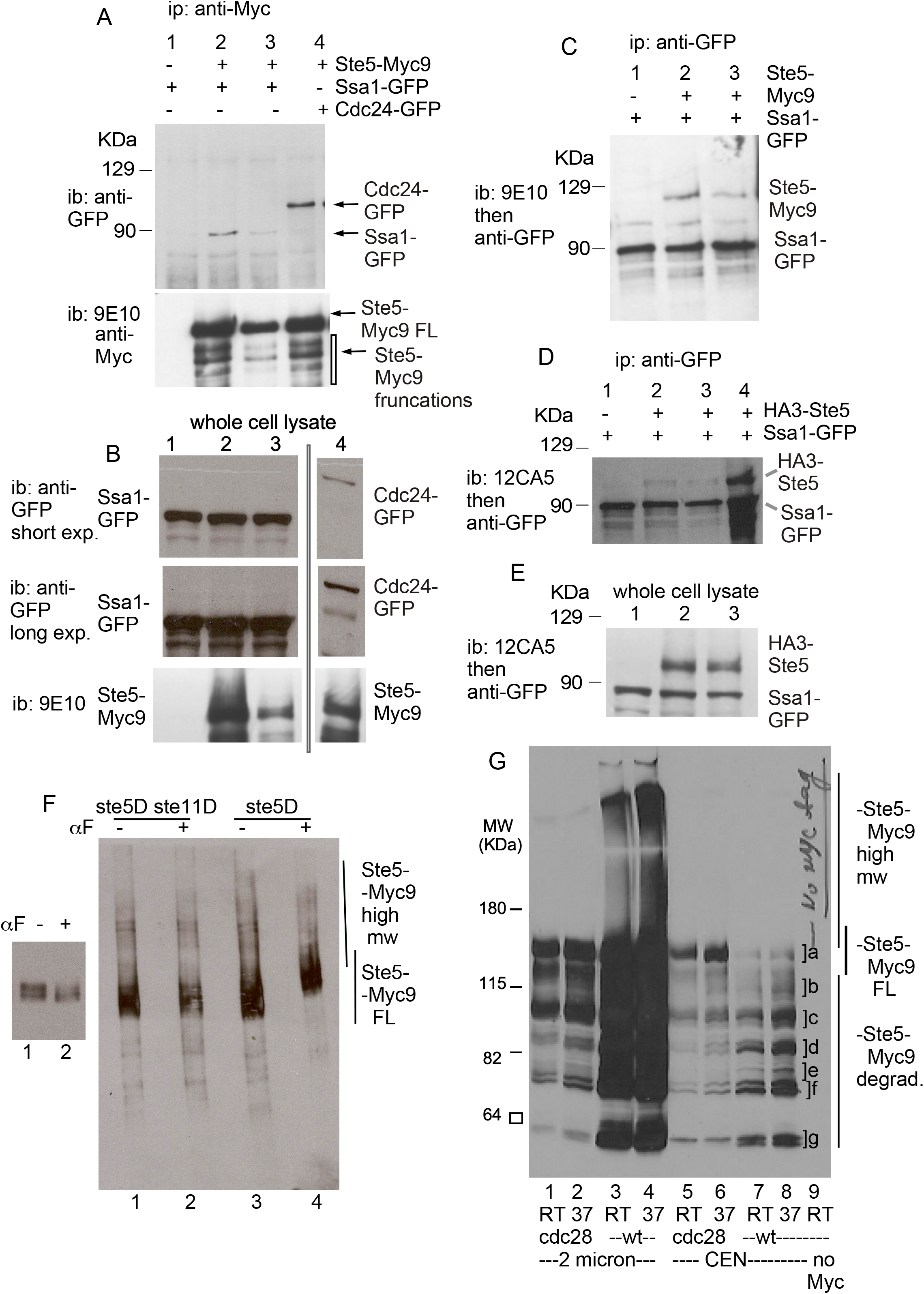
Ste5 co-immunoprecipitates with Ssa1-GFP and vice versa. **A.** Ste5-Myc9 co-IPs(co-IPs) Ssa1-GFP and *vice versa*. Co-IPs were done on whole cell lysates (extracts) prepared from EY1775 cells expressing STE5-MYC9 (pSKM90) with or without SSA1-GFP (pYMW122). Ste5-Myc9 was IP’d with 9E10 monoclonal antibody and Ssa1-GFP was IP’d with anti-GFP polyclonal antibodies. A control IP was done on extracts with Ste5-Myc9 co-expressed with GFP-Cdc24. lanes 2 and 3 are different plasmid transformants expressing different levels of Ste5-Myc9.B Immunoblots of whole cell extracts of strains in A. C. IPs of Ssa1-GFP with Ste5-Myc9. Anti-GFP polyclonal antibodies were used to IP Ssa1-GFP and blots were probed with either anti-GFP polyclonal antibodies or 9E10 to detect Ste5-Myc9. D. Co-IPs of Ssa1-GFP with HA3-Ste5. HA3-Ste5 was detected with 12CA5 monoclonal antibody. E. Immunoblot of whole cell lysates from strains in D. F. Ste5-Myc9 in native polyacrylamide gels from whole cell extracts prepared with extra phosphatase inhibitors and proteasome inhibitor. Ste5-Myc9 was expressed from 2 micron plasmid in EY1775 (W303a *MATa bar1D ste5D*) and EY1881 (W303a *MATa bar1D ste5D ste11D*). G. Comaprison of full-length, high molecular weight and truncated forms of Ste5-Myc in *CDC28* and *cdc28-4* strains. Ste5-Myc9 was expressed from 2μ (pSKM19) and CEN (pSKM12) plasmids in W303a *CDC28 (*EY957) and *cdc28-4* (PY1236) strains (44) grown at room temperature and after 3 hours shifted to 37°C and whole cell extracts prepared. Lanes 1,2: *cdc28-4 STE5-M9-2μ* RT, 37°C. Lanes 3, 4: WT *STE5-MYC9-2μ* RT, 37°C. Lanes 5,6: cdc*28-4 STE5-MYC9-CEN* RT, 37°C. Lanes 7,8: WT *STE5-MYC9-CEN* RT, 37°C. Ste5-Myc9 is detected with 9E10 anti-Myc monoclonal antibody. .]a high mw forms, | high mw forms, a] full-length, ]b-g truncated forms.

**FIG 3.**
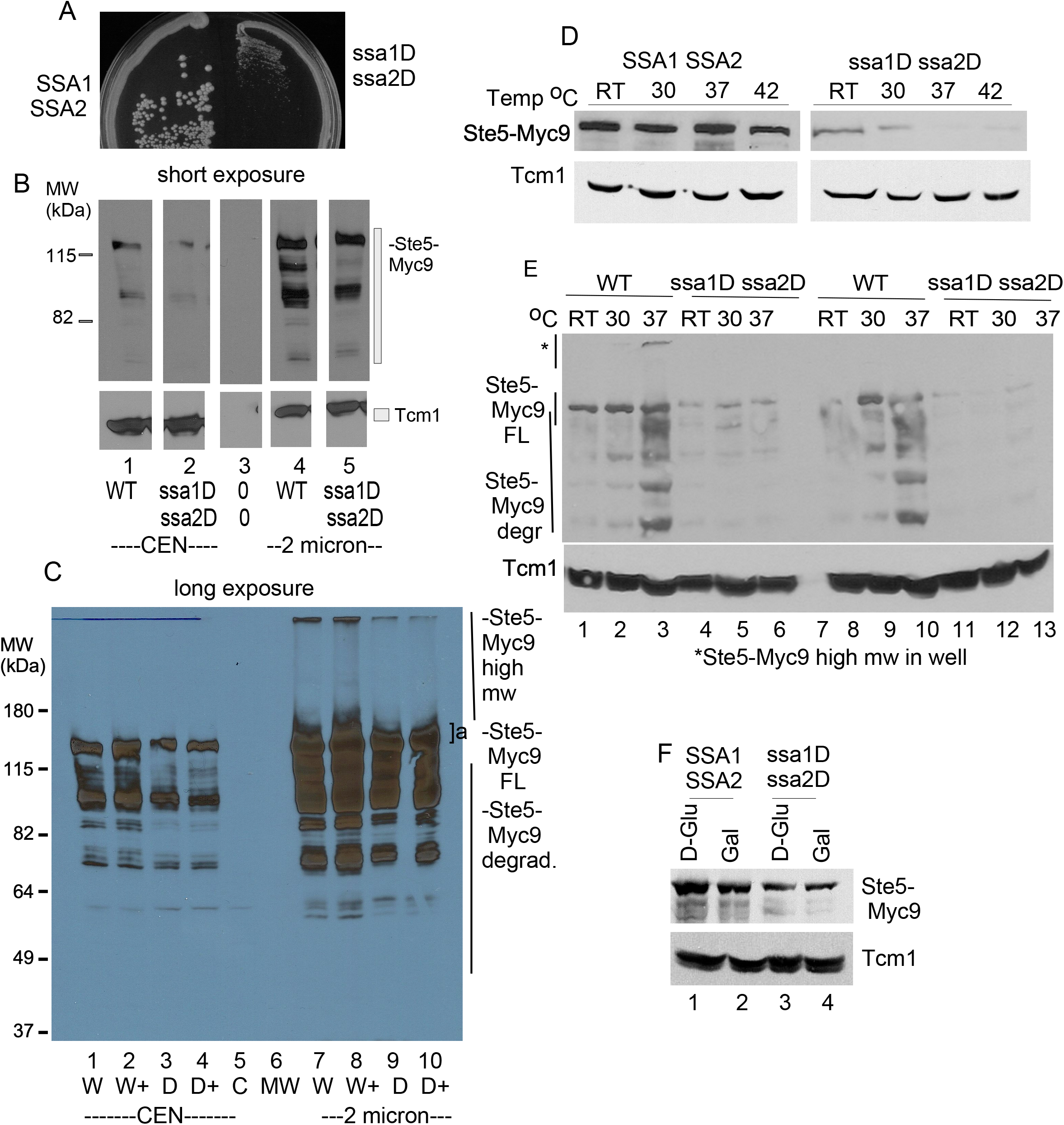
The abundance of Ste5 is lower in a *ssa1D ssa2D* double mutant. A. *ssa1D ssa2D* strains grow poorly. *SSA1D SSA2D* (EY3141) and *ssa1D ssa2D* (EY3136) strains were streaked on a YPD plate and grown at 30°CThe plate was photographed after 3 days. B-C. Ste5-Myc9 abundance and high molecular weight forms are reduced in a *ssa1D ssa2D* strain. Whole cell extracts were prepared from EY3139 *MATa ssa1::HIS3 ssa2::LEU2*, EY3141 *MATa SSA1D SSA2D* strains with *STE5-MYC9-CEN-TRP1* (pLSSte5Myc9TRP)(lanes 1-4), *CEN-TRP1* (EB406) (lane 5) or *STE5-MYC9-LEU2-2*μ (pLS40, lanes 7-10) grown in either SC-tryptophan or SC-leucine medium containing 2% dextrose and incubated with α factor where indicated. W=WT, D=*ssa1D ssa2D*. C is a short exposure of lanes 1,3,4,7,9 of the long exposure blot in D. D-E. Ste5-Myc9 *CEN* abundance is greatly reduced in an *ssa1D ssa2D* strain especially as temperature increases. Strains harboring either *STE5-MYC9-CEN* (pSKM12) or *STE5-MYC9-2*μ (pSKM19) were grown in SC-uracil medium containing 2% dextrose at different temperatures. Cells were pre-grown at room temperature (∼25°C) then shifted to pre-warmed medium at either 30°C, 37°C or 42°C shaken for hours then pelleted and extracts prepared. The immunoblots were first probed for Ste5-Myc9 then stripped and probed for Tcm1. F. Comparison of Ste5-Myc9 levels in *SSA1D SSA2D* and *ssa1D ssa2D* strains grown with either dextrose or galactose. Samples were grown as in Ai in SC-uracil medium with either 2% dextrose (Dex) or 2% galactose (Gal).

### Identification of 70 kd protein

Ste5-Myc9 was expressed from its own promoter on a multicopy *2μ* plasmid pSKM19 in EY1775 *MATa bar1D ste5D* cells in SC-uracil selective medium with 2% dextrose. Cells were grown and whole cell extracts prepared as described (86). In Fig. 1B, Ste5-Myc9 was immunoprecipitated (IP’d) with mouse 9E10 monoclonal antibody from 10 mg of yeast whole cell extracts prepared using liquid nitrogen and grinding with a mortar and pestle to break the cells as in (86). The buffer had 20 mM Tris-HCl pH 7.2 (Fisher), 125 mM potassium acetate (Fisher), 0.5 mM EDTA (Sigma), 0.5 mM EGTA (Sigma), 0.1% Tween 20 (Sigma), 1 mM DTT (BioRad), 20 mM 12.5% glycerol (Fisher), 10 μg/ml pepstatin (Sigma), 1mM benzenesulfonylfluoride (Sigma), 1 mM sodium ortho-vanadate (Sigma), 25 mM β-glycerophosphate (Sigma). The 70kd Ssa1,2 proteins were isolated by electrophoresis on an SDS polyacrylamide gel followed by staining with colloidal coomassie blue (Invitrogen). The 70kd band was excised from the gel and subjected to reduction, carboxyamidomethylation and in-gel digestion with trypsin. Recovered peptides were separated by microcapillary reversed-phase high-pressure liquid chromatography (HPLC) and analyzed using a Finnegan LCQ qudadrupole ion trap mass Spectrometer by 3D-ion trap tandem mass spectrometry (LC-MS/MS)(by Eric Spooner and Daniel P. Kirby of Harvard Microchemistry). Two peptide sequences were identified: 1-I*I*NEPTAAAI*AYGL*DK*-16 (peptide 1 theoretical monoisotropic mass 1458.77, experimental monoisotropic mass 1458.49) and 1-(SQ)*VDEI*VL*VGGSTR-14 (peptide 2 theoretical monoisotropic mass 1658, experimental monoisotropic mass 1658.69) (Datafile 0719yfse5-gt.dat) with qualifications indicated as an asterisk. Qualification in the sequence determination from MSMS spectra are necessary due to indistinguishable molecular mass of pairs of amino acids (the isobaric pairs are Leu/Ile MW 113, or Gln/ Lys MW 128 and Phe/oxyMet MW 148) which is indicated by an asterisk, and from areas of the MSMS spectra that could be solved only by pairs of amino acids, which is indicated by both a parenthesis and an asterisk.

### Densitometry

The band intensities of immunoblots were measured using either ImageJ software (ImageJ 1.50i, Wayne Rasband, National Institutes of Health, U.S.A. http://imagej.nih.gov/ij) or Adobe Photoshop CS5 (Version 12.0 x64). TIFFs of gels were prepared, all 300 resolution, 8 bit, all made without image contrast adjustment or tiff compression, and analyzed using ImageJ software from the N.I.H. and associated information in addition to European Union methods from Certus Technology. For each band, all lanes within boxes for area plots were cropped with the line tool and area was measured with the wand tool. Average background (calculated from several readings) was subtracted. Two to three readings were made for each measurement. In some instances, when ImageJ software was no longer compatible with updated operating systems, the histogram function of Canvas Draw was used for densitometry on non-adjusted images.

### β-galactosidase assays

The FUS1::ubiYlacZ construct in pDL1460 (88, 89) has *S. cerevisiae* ubiquitin with tyrosine fused to the N-terminus of beta-galactosidase which reduces its half-life from >20 hours to 10 minutes and avoids confounding effects of accumulation. Cells were grown overnight at room temperature to an A_600_ of ∼0.3, pelleted, and resuspended at an A_600_ of 0.3 in 10 ml aliquots of fresh media. α-factor was added from a concentrated stock (6 mM in methanol) and samples were shaken for 2 hours, pelleted, washed once with ice-cold water and frozen at −80°C. Pellets were thawed and extracts prepared and quantified exactly as described (85). Units of beta-galactosidase were calculated by the following formula: Units = [A_420_ x (1.7/0.0045)]/ [time (min) x extract volume (ml) x protein concentration (mg/ml)].

### Extract centrifugation

150 mls of cells were grown to A_600_ of 0.3 at room temperature then pelleted in either a Sorvall RC5B or Sorvall RT 6000B centrifuge at 5,000 rpm for 5 minutes at 4°C. Pellets were washed once with ice cold sterile doubly distilled water that had been chilled to 4°C, then they were resuspended in fresh ddH_2_0, transferred and pelleted again. Pellets were drained and frozen in a dry ice/ethanol bath and stored overnight at −80°C. After being thawed on ice, 1 ml lysis buffer and glass beads were added to the pellets and the samples were vortexed 5 times for 30 seconds each, chilling on ice between each vortexing. Then 0.4 ml of lysis buffer were added to the samples on ice and they underwent a sixth vortex for 30 seconds. Samples were then spun in the swinging bucket Sorvall RT 6000B centrifuge at 5,000 rpm (3,000 x g) for 10 min at 4°C. Supernatants were transferred equally to two chilled eppendorf tubes, saving one as a control for total input. The other tube of sample was centrifuged in a refrigerated Eppendorf 5415C centrifuge at 14,000 rpm (16,000 x g) for 15 minutes and then the supernatant was transferred to a new tube. An equal volume of lysis buffer was added to each pellet. Protein concentration was determined and 5x loading buffer added to samples for running on an SDS polyacrylamide gel. The buffer used for extract preparation had 25 mM Tris-Cl pH7.4 (Fisher), 200 mM NaCl (Sigma or Fisher), 15 mM MgCl_2_ (Sigma or Fisher), 1 mM DTT (BioRad), 15 mM EGTA (Fisher), 0.1% triton-X 100 (Fisher), 10% glycerol (Fisher), 1 mg/ml phenylmethylsulfonyl (PMSF), 4 mM 1,10-phenanthroline (Sigma), 2 mM benzamidine (Sigma), 5 μg/ml leupeptin (Sigma), 5 μg/ml pepstatin A (Sigma), 5 μg/ml chymostatin (Sigma), 5 μg/ml aprotonin (Sigma), 0.25 mM sodium ortho-vanadate (Sigma), 0.25 mM sodium meta-vanadate (Sigma), 50 mM sodium fluoride (Sigma), 1 mM sodium azide (Sigma). In some experiments we also added 50 μM proteasome inhibitor Z-Ile-Glu(OBut)-Ala-Leu-H (aldehyde) (Peptide International) and phosphatase inhibitor cocktail 1 (microcystin LR, cantharidine and (-)-p-brotetramisol, Sigma).

### Fus3 kinase assay

The EY3136 and EY3141 strains were made *bar1D* by crossing them to S288c BY4742 (RG1104) *MATa bar1D0::KAN^R^* strain. No *MATa bar1D0::KAN^R^ ssa1::HIS3 ssa2::LEU2* ascospores were recovered out of ∼40 tetrads dissected, suggesting that the *MATa bar1D0::KANR ssa1::HIS3 ssa2::LEU2* ascospores germinate poorly based on presence of dead ascospores among tetrads dissected. Two *MATα bar1D::KANR ssa1::HIS3 ssa2::LEU2* ascospores were recovered and switched to *MATa* with a *pGAL-HO* plasmid, then ura3-clones that had lost the plasmid were used for analysis (EYL1798). Strains were transformed with pYEE121 (FUS3-HA5-CEN-URA3) and pYEE128.30-1 (fus3K42R-HA5-CEN-URA3) for kinase assays. Approximately 150 mls of cells were grown to A600 of 0.7-0.8 and processed exactly as described using an extraction buffer that contained 250 mM NaCl. Transformations and growth were done at room temperature.

### Immunofluorescence microscopy

Indirect immunofluorescence and live cell imaging were performed exactly as described (35). Quantification of nuclear and cortical localization were performed as described (16, 35), counting at least 500 cells for each sample. Cells were examined under an Axioscope 2 microscope (Carl Zeiss, Thornwood, N.Y.) linked to a digital camera (C4742-95, Hamamatsu, Bridgewater, N.J.) with filters (set 41018 for GFP and set 51006 Texas Red filter for fluorescein isothiocyanate) from Chroma Technology (Brattleboro, Vermont, USA). Strains harboring pCUP1-GFP-STE5 (pSKM21; 41) were grown overnight at room temperature in selective SC medium to early logarithmic phase (A_600_∼0.15), then shifted to selective SC medium containing 0.5 mM CuSO_4_ for 2 hours before treating cells with α factor as previously described (16, 35). Cells were examined on a Nikon TE2000E epifluorescence microscope using 360/40, 457/17 (DAPI) filter set (Chroma Corp.). Digital images were collected with a Cooke 12-bit SensiCam CCD driven by IPLab 3.9 software. Whole-image adjustments were performed in Adobe Photoshop. Some cells were visualized with a Hamamatsu ORCA ER digital camera and MetaMorph 7 software at the Department of Cell Biology Microscopy Facility of the Harvard Medical School.

## Results

### Ste5 co-purifies with Ssa1

We purified proteins that co-immunoprecipitate (co-IP) with over-expressed Ste5-Myc9 from logarithmically dividing cells grown at 30°C before and after α factor treatment. We used low NaCl extraction conditions and the 9E10 monoclonal antibody to the Myc epitope. A number of proteins co-IP’d specifically with Ste5-Myc9, but not with untagged Ste5. Figure 1B labelling indicates six bands that are most prominent: Ste5-Myc9 (∼115 kDa), band “a” also labeled as ∼70 kDa, bands “b”, “c” (which are ∼68-58 kDa) and IgG2a (∼50 kDa)(Fig 1B). The signal transduction partners of Ste5 that are closest in size to bands “a”, “b” and “c” are Ste7 (57,723.7 Da) and Ste11 (80,718.8 Da) whereas Fus3 (40,770.5 Da) and Kss1 (42,680.6 Da) may be obscured by the wide IgG2a band. Ste5-Myc9 is largely full-length and the +α factor sample migrates more slowly (Fig 1B, lane 3), presumably from hyper-phosphorylation (38). Thus, the preparation of Ste5-Myc9 was intact and the α factor treatment induced a robust pheromone response. Band “a” is present in an amount similar to that of Ste5-Myc9, both in absence and presence of α factor treatment (Fig 1B, band “a” lane 2 –αF, lane 3 +αF). The band “a” protein is specifically associated with Ste5-Myc9 and not present in IPs of extracts expressing untagged Ste5 (Fig 1B, lane 1). Similar results were found in repeated immunoprecipitations and a scale-up for mass spectrometry. ImageJ densitometry of the coomassie stained gel in Fig. 1B indicates a ratio of 1.2 of Ste5-Myc9:70 kDa Hsp70 –α factor and a ratio of 0.6 Ste5-Myc9: 70 kDa Hsp70 +α factor suggesting less Hsp70 associated with Ste5-Myc9 in the +α factor extract (TableS2). The two other bands, “b,” “c,” are present in sub-stoichiometric amounts compared to Ste5-Myc9. Focus was placed on band “a” due to its high abundance. A large-scale IP of Ste5-Myc9 was performed, the ∼70 kDa band “a” was excised from the ge, proteolyzed with trypsin, and microcapillary HPLC-ion trap tandem mass spectrometry was done on two trypsin-generated peptides. The tandem mass spectra of two major HPLC peaks identified residues 170-185 (IINEPTAAAIAYGLDK) of Hsp70 isoforms Ssa1, Ssa2, Ssa3 and Ssa4 (Fig 1C). The second peptide identified residues 325-338 of Ssa1 and Ssa2 (SQVDEIVLVGGSTR). Thus, the majority of Ste5-Myc9 was complexed to Ssa1 and/or Ssa2 both in absence and presence of α factor.

To confirm the presence of Ssa1, we determined whether Ste5-Myc9 and GFP-Ssa1 co-IP using higher salt (200 mM NaCl), nondenaturing immunoprecipitation conditions we have used to reveal Ste5 binding to Ste4, Fus3, Ste7, Ste11, Cdc24, and Msn5/Ste21 (9,10,16,23,86). Ste5-Myc9 associates with Ssa1-GFP when Ste5-Myc9 is IP’d with 9E10 anti-Myc monoclonal antibody (Fig 2A-B) and Ste5-Myc9 associates with Ssa1-GFP when Ssa1-GFP is IP’d with anti-GFP polyclonal antiserum (Fig2C). The interaction between Ste5-Myc9 and Ssa1-GFP is specific based on the absence of a signal when extracts lack Ste5-Myc9 (Fig 2A, lane 1), more Ssa1-GFP in the co-IP when more Ste5-Myc9 is in the whole cell lysate (Fig 2A,B compare lanes 2,3), and the correct migration position of Ssa1-GFP for its predicted size which is distinct from GFP-Cdc24, a positive control (Fig 2B, lane 4). HA3-Ste5 also co-immunoprecipitated with Ssa1-GFP (Fig 2D, lanes 2-4) with a better signal when more whole cell lysate was used for the IP (Fig 2D Lane 4. Fig 2E is the whole cell lysates). These results confirm that Ste5 is complexed to Ssa1 under native conditions.

### Ssa1 and Ssa2 positively regulate Ste5 abundance

Ste5 has a complex banding pattern in native- and SDS- and polyacrylamide gels that is affected by expression level, genetic background and posttranslational modification by Cln/Cdk Cdc28, Fus3 (and other protein kinases), ubiquitinylation by SCF^Cdc4^ and the presence of phosphatase and numerous protease inhibitors including an inhibitor of the proteasome (i.e.. Z-Ile-Glu(But)-Ala-Leu-H (aldehyde)) during whole cell lysate preparation. In a native gel, a broad diffuse banding pattern is detected for full-length Ste5-Myc9 that shifts to higher apparent molecular weight with α factor treatment in a manner that requires signaling through the mating MAPK cascade is blocked by a *ste11D* mutation (Fig 2F, 38). Ste5-Myc9 also accumulates high molecular weight species that are likely ubiquitinylated, because they enrich in presence of proteasome inhibitor (Fig 2F) and disappear when Cdc28 CDK is mutated (Fig 2G). Strong overexpression of Ste5-Myc9 with a leu2-d 2 micron plasmid, leads to increased amounts of these high molecular weight species which smear and aggregate in the gel well during SDS-PAGE (Fig 2G, compare lanes 3,4 with 7,8 centromeric plasmid). The high molecular weight smear is more pronounced in extracts from wild type (WT) cells grown at 37°C compared to room temperature (Fig 2G, lanes 3,4). The *cdc28-4* mutation in CDK Cdc28 increases the relative amount of full-length Ste5-Myc9 at 37°C and room temperature (Fig 2G, lanes 5,6, ]a) and reduces by 0.2 mm the spread of the full-length band, presumably because of loss of phosphates from Cln/Cdc28 (Fig 2G, ]a, lanes 2, 7, 9). The *cdc28-4* mutation also abolishes aggregated forms of Ste5-Myc9 in the gel well and high molecular weight forms in the smear, suggesting these species include phosphorylated and ubiquitinylated versions of Ste5-Myc9 (Fig 2G, compare lanes 1,2 with 3,4). Overexpression of Ste5-Myc9 also leads to accumulation of proteolyzed fragments whose abundance declines in a *cdc28-4 ts* mutant, presumably from loss of the ubiquitinylation-mediated degradation (Fig 2G, lanes 6-9).

We determined the effect of a *ssa1D ssa2D* double mutation on the abundance and gel migration pattern of Ste5-Myc9 expressed from *CEN* (∼1 copy/cell) and *2μ* (multiple copies/cell) plasmids. A M*ATα ssa1D ssa2D* double mutant generously provided by the Craig lab grew poorly compared to the isogenic *MATa* wild type *SSA1D SSA2D* strain at all temperatures examined and generated colonies that were only 19.4% the size of wild type (Fig 3A shows day 3 of strains streaked on a YPD plate kept at 30°C, mean colony size in arbitrary units for wild type is 4.06 SD 1.05 and for ssa1D ssa2D is 0.788 SD 0.18).

Ste5-Myc9 protein levels were reduced in the *ssa1D ssa2D* double mutant at room temperature during vegetative growth when it was expressed from its own promoter on a *CEN* plasmid (Fig 3B, lanes 1-2). Densitometry showed that the level of Ste5-Myc9 was ∼20% that of wild type after normalization with ribosomal protein Tcm1 (Table S2). This observation was striking given that Ste5 has a long half-life of ∼90 minutes by immunoblot analysis (37–38) and 3 hours by heavy lysine isotope labeling (90) and the abundance of Ste5 protein, but not its mRNA, increases somewhat during α factor stimulation, peaking after 1 hour of stimulation (38). When Ste5-Myc9 was overexpressed with the 2-micron multicopy plasmid, the abundance of Ste5-Myc9 was ∼57% that of the wild type control, indicating that overexpression bypasses some of the reduction caused by loss of *ssa1D ssa2D* (Ste5-Myc9 2μ Fig 3B, lanes 4,5; Table S2). Longer exposure of the immunoblot revealed that Ste5-Myc expressed in the wild type Craig strain formed aggregated species in the well of the gel and a somewhat higher molecular weight smear (Fig 3C, lanes 7,8). In addition, degradation fragments of Ste5-Myc9 were present with both 2 micron and *CEN* expression levels (Fig 3C, lanes 1,2,7,8). By contrast, Ste5-Myc9 from the *ssa1D ssa2D* strain had no obvious aggregated forms, less obvious higher molecular weight smear, a narrower full-length Ste5-Myc9 band (Ste5-Myc9FL) and an altered pattern of Ste5-Myc9 fragments (Fig 3C, lanes 2,3,9,10). Thus, the *ssa1D ssa2D* mutations appear to affect the abundance and integrity of Ste5.

To test the generality of the requirement of Ssa1 and Ssa2 and the effect of temperature, we created additional *MATa ssa1D ssa2D* and *MATa SSA1D SSA2D* strains by sporulating and dissecting ascospores from a *MATa/MATα ssa1/+ ssa2/+ ssa3/+ ssa4/+* heterozygous diploid from the Craig lab (Table S1). Ste5-Myc9 was expressed from a *CEN* plasmid in selective medium containing 2% dextrose in one of the *MATa ssa1D ssa2D* ascospore strains (the whole cell lysates were prepared with Z-Ile-Glu(But)-Ala-Leu-H (aldehyde) as in Fig 2F). In the wild type control, some of the Ste5-Myc9 was in the gel well at 37°C and fragmented (Fig 3D, lanes 1-3,8-10), suggesting elevated temperature leads to more Ste5 aggregation and degradation. Remarkably, when expressed at native levels, Ste5-Myc9 was nearly undetectable at 30°C, 37°C, and 42°C in *ssa1D ssa2D* strains compared to in wild type (Fig. 3D-E). Densitometry revealed that Ste5-Myc9 abundance was ∼20% that of wild type at room temperature and further reduced at higher temperatures to ∼3.1% at 30°C, ∼0.03% at 37°C and ∼0.02 % at 42°C. By contrast, the control Tcm1 protein had similar abundance in WT and *ssa1D ssa2D* strains at all temperatures and was not fragmented (Fig. 3D-E, Table S2). Similar results were obtained with two different transformed ascospores (Fig. 3E, Table S2). Thus, temperature increase appears to increase degradation of Ste5-Myc9 both in wild type and the *ssa1D ssa2D* double mutant. The effects are most likely posttranscriptional since the Ste5 promoter is not regulated by Ssa1, Ssa2 or Hsf1 or Msn2/Msn4. Furthermore, the truncated species are unlikely to be premature translational stop peptides since the Myc tags are at the carboxyl-terminus.

Reduced levels of Ste5-Myc9 were also apparent when *ssa1D ssa2D* cells were grown in 2% D-glucose and 2% D-galactose (Fig 3H; note that Ste5-Myc9 abundance is slightly lower in 2% galactose compared to 2% D-glucose) further substantiating a general requirement for Ssa1 and Ssa2 for Ste5 abundance under a different carbon source. Thus, Ssa1 and Ssa2 are important for Ste5 protein abundance and integrity at room temperature and crucial at elevated temperatures.

### Loss of Ssa1 and Ssa2 alters the integrity of Ste5

We have previously shown the Ste5 exists in a high molecular weight complex that sediments with kinases in a glycerol gradient and is likely also associated with cytoskeletal proteins. We examined the distribution of Ste5 in supernatant and pellet fractions of our whole cell lysate preparations from WT *SSA1 SSA2* (EY3407) and *ssa1D ssa2D* (EY3409) strains, using an initial brief 3,000 x g centrifugation followed by a 16,000 x g centrifugation that is known to pellet nuclei, cell debris, contractile/cytoskeletal apparatus, cell wall and some aggregated proteins. The lanes in the immunoblot in Fig 4A have 100 μg protein for total and supernatant samples and 20 μg protein for pellet samples.. Strikingly, densitometry revealed ∼4-fold more full length (FL) Ste5-Myc9 in *WT* compared to in *ssa1D ssa2D* and different pellet to supernatant distributions: ∼10% of Ste5-Myc9 FL was in the *ssa1D ssa2D* pellet compared to ∼76% in pellet of WT extracts. Furthermore, more Ste5-Myc9 was fragmented in the supernatant of the *ssa1D ssa2D* lysate compared to the WT lysate (Fig 4A, note black marks at very top of blot are from labeling the film). These results further substantiate that the *ssa1D ssa2D* mutations alter abundance and integrity of Ste5 and suggest the mutations may change subcellular distribution.

**FIG 4.**
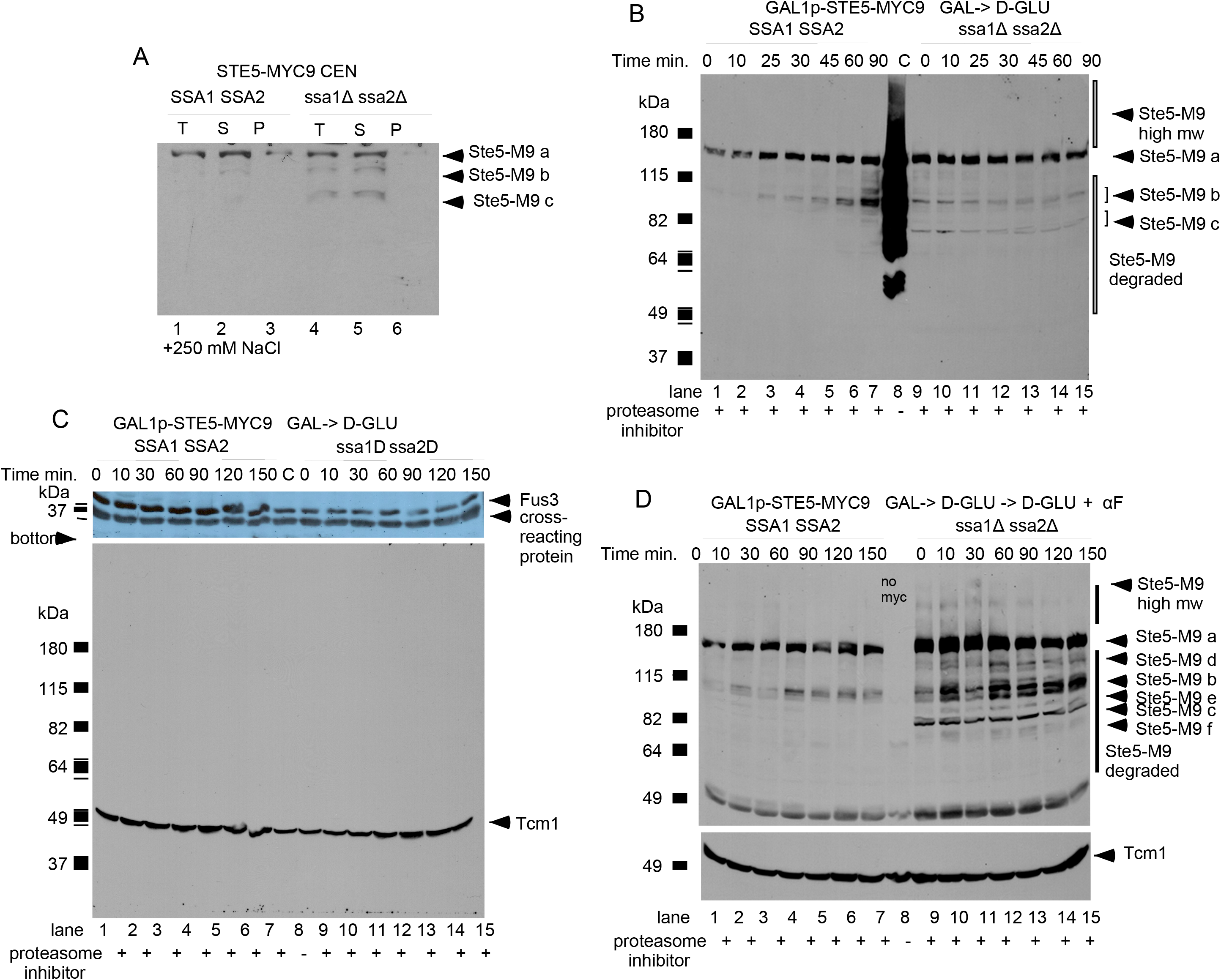
N-terminal truncations of Ste5-Myc9 accumulate in the *ssa1D ssa2D* double mutant. A. Comparison of Ste5-Myc9 in pellet and supernatant of wild-type and *ssa1D ssa2D* extracts subjected to 16,000 x g centrifugation. EY3407 *SSA1D SSA2D* and EY3409 *ssa1D ssa2D* cells expressing pSKM12 (*STE5-MYC9 CEN URA3*) were grown to A_600_ of 0.3 and extracts prepared, then an aliquot was centrifuged at 16,000 x g into supernatant (S) and pellet (P) fractions. The immunoblot has 100 μg of total (T) WCE and supernatant and 20 μg of pellet. “a” is full length Ste5-Myc9 and “b” and “c” are truncation products. B-D. Ste5-Myc9 N-terminal truncation products accumulate in a *ssa1D ssa2D* double mutant. B. *GAL1* promoter shut-off of Ste5-Myc9 during vegetative growth. Extracts prepared from EY3136 and EY3141 cells harboring pSKM30 (*GAL1p-STE5-MYC9 URA3 CEN*). Cells were pregrown in SC-uracil-2% raffinose medium, then grown in SC-uracil-2% galactose medium for 4 hours then pelleted, washed and resuspended in 2% dextrose medium. “C” is for control and is the MW standard co-run with overloaded EY1775 + *STE5-MYC9 CEN* (W303a background) grown in 2% dextrose medium. C. Abundance of Fus3 and Tcm1 during *GAL1prom-STE5-Myc9* promoter shut-off. Cells were grown as in B. except they were 5 hours in SC-uracil-2%galactose medium. Blot is probed with anti-Fus3 antibodies and anti-Tcm1 antibody. D. Ste5-Myc9 and Tcm1 during *GAL1-STE5-MYC9* shut off in WT and *ssa1D ssa2D* cells treated with α factor. Cells and extracts prepared as in C except that samples were treated with 5 μM α factor. Blot is probed with 9E10 and anti-Tcm1 antibodies, D is probed with 9E10 and Tcm1 antibodies. The “a-f” indicate full-length and truncated forms of Ste5-Myc9.

To examine a previously synthesized, post-transcriptional pool of Ste5-Myc9, we induced its overexpression with the *GAL1* promoter for 5 hours in galactose medium, then stopped expression with dextrose medium and followed abundance over time in the absence and presence of α factor as done previously (38). In a W303a *ste5* null strain, the abundance of Ste5-Myc9 declines to 24% its initial level by 150 minutes (38) and it accumulates as high mw species when overexpressed sufficiently in W303a *ste5D* with a 2-micron plasmid in selective media containing 2% dextrose (Fig 4B, lane 8 labeled C). The abundance of *GAL1*promoter-induced full-length Ste5-Myc9 was significantly lower in the wild type *SSA1D SSA2D* Craig lab background compared to W303a and did not decline noticeably over 90 minutes, but more fragments of Ste5-Myc9 accumulated by 90 minutes (”Ste5-Myc9 a”, Fig 4B, lanes 1-7 compared to lane 8, note Ste5-Myc9 fragment labeled b). Some reduction in abundance of full-length Ste5-Myc9 (labeled as “Ste5-Myc9 a”) could be discerned in the *ssa1D ssa2D* strain (Fig 4B, lanes 9-10). We did not detect high molecular weight species of Ste5-Myc9. A proteolytic fragment indicated as “Ste5-Myc9 b” was present in both the Craig background wild-type and *ssa1D ssa2D* whole cell extracts, but the pattern of accumulation was distinct. The wild type “Ste5-Myc9 b” fragments are visible at 25 min to 90 min and increase in abundance. By contrast, in the *ssa1D ssa2D* extracts, the “Ste5-Myc9 b” fragments are present at all time points with no increase in abundance. Notably, a second set of proteolytic fragments, “Ste5-Myc9 c” were readily visible in the *ssa1D ssa2D* extract but not the wild type extract. Thus, Ste5-Myc9 fragments with distinct proteolytic sites accumulate in the *ssa1D ssa2D* mutant compared to wild type.

Further changes in the Ste5-Myc9 fragment profile occurred with α-factor (Fig 4D). The “Ste5-Myc9 a” full-length band became 1.7+/-0.22-fold broader in the *ssa1D ssa2D* strain compared to in wild type together with a low amount of high molecular weight Ste5-Myc9 species at the 0 to 90 minute time points. In addition, the *ssa1D ssa2D* extracts had a broader “Ste5-Myc9 b” set of fragments, the “Ste5-Myc9-b” and “Ste5-Myc9 c” fragments were more abundant, and there were additional sets of fragments (labeled “d,e,f,g” in Fig 4D). By comparison, in the wild type extracts with α factor, the Ste5-Myc9 fragments at position “b” were only weakly detected at the 10 minute time point and there were no obvious “Ste5-Myc9 c, d,e,f,g” fragments (Fig 4B,D). Thus, Ste5-Myc9 is more vulnerable to proteolysis without Ssa1 and Ssa2 especially during α-factor signaling. The strong protective function of Ssa1 and Ssa2 for Ste5-Myc9 was quite specific, because the *ssa1D ssa2D* mutations did not obviously alter the integrity of the protein band profiles of Fus3, Tcm1 (Fig 4C), or Fus3-HA, Ste7-Myc or Kss1 (for example, Kss1 accumulates a single proteolytic fragment in wild type that is of same abundance in *ssa1D ssa2D*).

Antibody accessibility is a well-established method to monitor changes in peptide flexibility, conformation, and binding to partners (e.g. 91,92). The N-and C-termini of Ste5 become more available to antibody in some *ste5* mutants (16). The N-terminal ∼1-161 and C-terminal ∼760-877 regions of Ste5 are predicted to have high intrinsic disorder according to seven intrinsic disorder algorithms (Fig S1B-E) suggesting that the ends of Ste5 may be more sensitive to conformational changes. We examined whether Ssa1 and Ssa2 influence the accessibility of antibodies to C-terminal Myc9 and N-terminal HA3 tags on Ste5. Strikingly, although wild type whole cell extracts had >3-fold more Ste5-Myc9 than *ssa1D ssa2D* whole cell extracts, 3-fold more Ste5-Myc9 and 7-fold more HA3-Ste5 were IP’d from the *ssa1D ssa2D* whole cell extracts than from the wild type whole cell extracts (Fig. 5A-C). ImageJ densitometry (Table S2) revealed 1.9 +/-0.26 s.d.-fold more Ste5-Myc9 was IP’d from *ssa1D ssa2D* extracts expressing the *STE5-MYC9 2μ* plasmid, although the abundance of Ste5-Myc9 was 3.4 +/-0.51 s.d. higher in the isogenic wild type strain (Fig 5A) and 1.7-fold more Ste5-Myc9 was IP’d from the *ssa1D ssa2D* extracts expressing the *STE5-MYC9 CEN* plasmid compared to wild type extracts, although the abundance of Ste5-Myc9 was 2.6-fold higher in the wild type extracts (Fig 5B). Moreover, 3.0-fold more HA3-Ste5 was IP’d from the *ssa1D ssa2D* strain expressing *HA3-STE5 CEN* plasmid than from the isogenic wild type strain, although the wild type strain had ∼7.3-fold more HA3-Ste5 than the *ssa1D ssa2D* strain. Therefore, both N-and C-terminal tags on Ste5 are more available to antibodies in the *ssa1D ssa2D* extracts.

**Fig 5.**
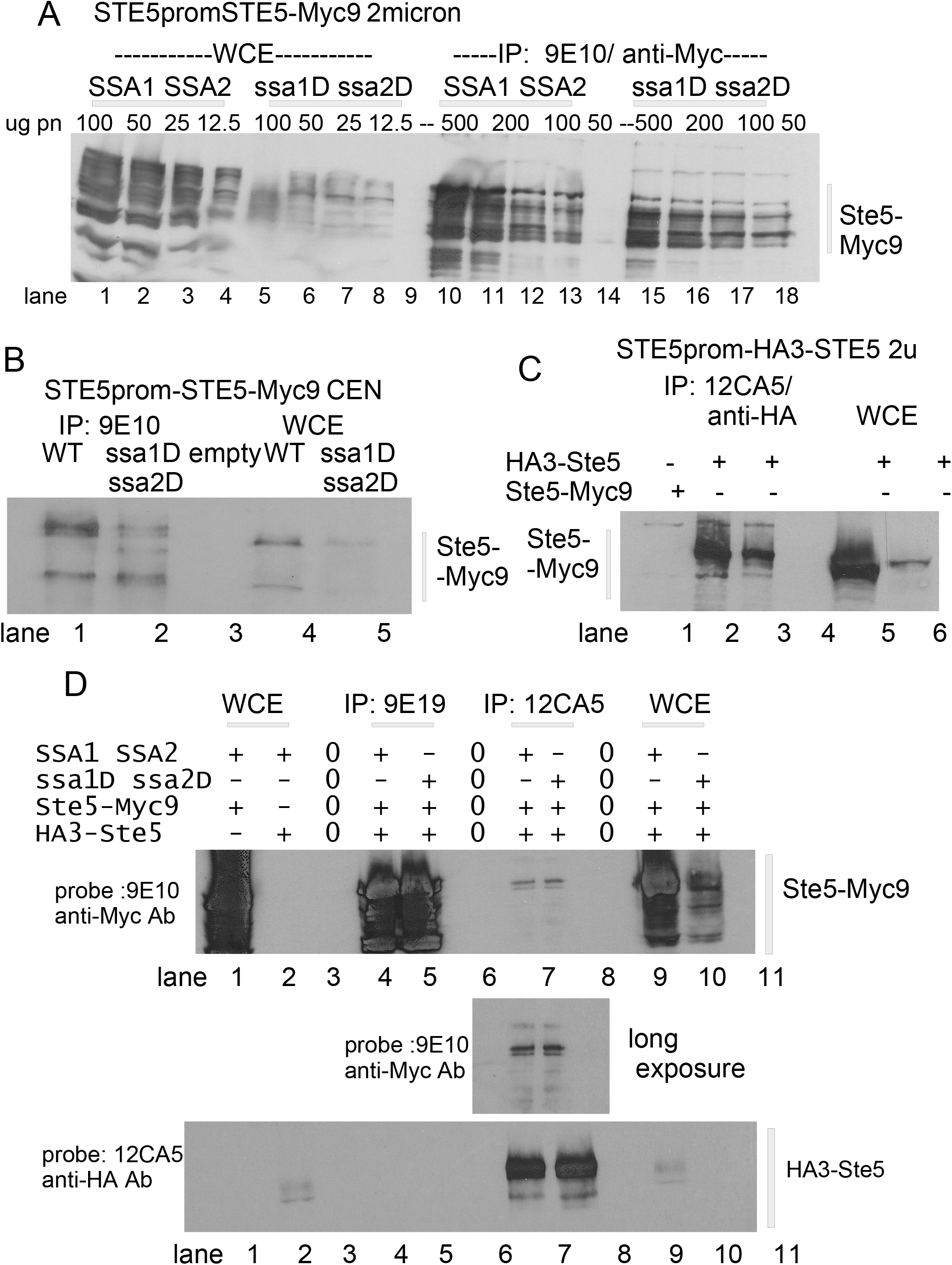
N- and C-terminal epitope-tagged forms of Ste5 from *ssa1D ssa2D* strains have enhanced antibody accessibility and oligomerization. A. *STE5-Myc9-2μ* WCE ips from WT and *ssa1D ssa2D* extracts. Decreasing μg amounts of whole cell extract were used. Lanes 1-4: *SSA1D SSA2D* WCE 100, 50, 25, 12.5; lanes 5-8: *ssa1D ssa2D* WCE 100, 50, 25, 12.5. Lane 9 empty. Lanes 10-14: *SSA1D SSA2D* WCE IPs 500, 200 and 100, 50. Lane 15 empty. Lanes 16-19: *ssa1D ssa2D* IP WCE 500, 200, 100, 50. Blot probed with 9E10 antibody. B. *STE5-MYC9-CEN* WCE ips from WT and *ssa1D ssa2D* extracts. 0.2 mg WCE was IP’d with 9E10 and compared to 50 μg WCE. Blot was probed with 9E10. Lane 1: *SSA1D SSA2D STE5-MYC9* WCE. Lane 2: *ssa1D ssa2D STE5-MYC9* WCE. Lane 3: *SSA1D SSA2D* Ste5-Myc9 IP. Lane 5-Myc9 IP. Lane 4: *ssa1D ssa2D* Ste5-Myc9 IP. C. *HA3-STE5-2μ* WCE ips from WT and *ssa1D ssa2D* extracts. 12CA5 IPs were done with 0.5 mg of WCE and run next to 0.1 mg WCE. Blot was probed with 12CA5. Lane 1: *SSA1D SSA2D* Ste5-Myc9 WCE IP with 12CA5. Lane 2: *SSA1 SSA3* HA3-Ste5 WCE IP with 12CA5. Lane 3: *ssa1 ssa1* HA3-Ste5 WCE IP with 12CA5. Lane 4 empty. Lane 5: *SSA1D SSA2D* HA3-Ste5 WCE. Lane 6: *ssa1D ssa2D* HA3-Ste5 WCE. D. Oligomerization. Ste5-Myc9 (pRM01) and HA3-Ste5 (pSKM26) co-expressed in *SSA1D SSA2D* and *ssa1D ssa2D* strains. For A-C 0.5mg whole cell extract was IP’d with either 12CA5 or 9E10 and then probed with 9E10 in the immunoblot. Lane 1: WCE *SSA1D SSA2D* Ste5-Myc9. Lane 2: WCE *SSA1D SSA2D* HA3-Ste5. Lane 3: Empty. Lane 4: IP 9E10, *SSA1D SSA2D* Ste5-Myc9 + HA3-Ste5. Lane 5: IP 9E10, *ssa1D ssa2D* Ste5-Myc9 HA3-Ste5. Lane 6: Empty. Lane 7: IP 12CA5 *SSA1D SSA2D* Ste5-Myc9 + HA3-Ste5. Lane 8: IP 12CA5 *ssa1D ssa2D* Ste5-Myc9 + HA3-Ste5. Lane 9: Empty. Lane 10: WCE *SSA1D SSA2D* Ste5-Myc9. Lane 11: WCE *ssa1D ssa2D* Ste5-Myc9. For A-D, extracts were prepared from wild-type and *ssa1D ssa2D* strains (EY3136, EY3141) expressing *STE5-MYC9 2μ* (pSKM19), *HA3-STE5* (pSKM87) or *STE5-MYC9 CEN* (pSKM12) grown at 30°C in SC-uracil medium containing 2% dextrose.

The active form of Ste5 may be a dimer that forms through internal RING-H2::RING-H2 and other intermolecular interactions (Fig.S1A, FigS2A,D; 38). Less than 1% of the total pool of Ste5 is detected as oligomers during vegetative growth (38) presumably because of auto-inhibition within the monomer (38, 93). We examined oligomerization between HA3-Ste5 and Ste5-Myc9 overexpressed with 2 micron plasmids in wild type and *ssa1D ssa2D* whole cell lysates (Fig 5D). The abundance of Ste5-Myc9 was several-fold less in the *ssa1D ssa2D* extracts compared to wild type (Fig 5D lanes 9,10) yet equivalent amounts of Ste5-Myc9 associated with HA3-Ste5 (Fig. 5D lanes 7,8). Therefore, Ste5-Myc9 dimerizes equivalently or slightly more in the *ssa1D ssa2D* extracts than the wild type extracts (Fig.5D).

### Defective cortical localization of Ste5 in the *ssa1D ssa2D* mutant

Localization of Ste5 to the cell periphery is associated with activation of signaling and shmoo formation and is thought to involve an induced conformational change co-incident with binding to Gβγ that correlates with RING-H2 domain oligomerization at the plasma membrane. GFP-Ste5 localized throughout the cytoplasm in the *ssa1D ssa2D* double mutant before and after α factor treatment, but did not efficiently accumulate at the growth site of the emerging projection tip (6.7% rim staining for *ssa1D ssa2D* compared to 70% for wild type after 90 minutes in 50 nM α factor, Table 1; Fig 6A shows representative experiment). The *ssa1D ssa2D* cells were more enlarged and misshapen compared to wild type which is a sign of reduced α factor signaling. The fluorescence signal of GFP-Ste5 was lower in the *ssa1D ssa2D* strain than the wild-type strain leading to fewer and less intensely fluorescing GFP positive cells, consistent with lower abundance. The cells that exhibited enrichment at the growth site were the most shmoo-like in morphology. These findings were corroborated by indirect immunofluorescence with two additional Ste5 constructs, Ste5-Myc9 and TAgNLS^K128T^-Ste5-Myc9 which has enhanced ability to be stably recruited to the cell cortex (35)(Table S3). Thus, Ssa1 and Ssa2 are required for efficient recruitment of Ste5 to the cell periphery even when the level of Ste5 is increased or when the Ste5-Myc9 derivative has enhanced ability to be recruited. Collectively, these findings suggest the integrity of Ste5 is not optimal for being recruited to the plasma membrane.

**FIG 6.**
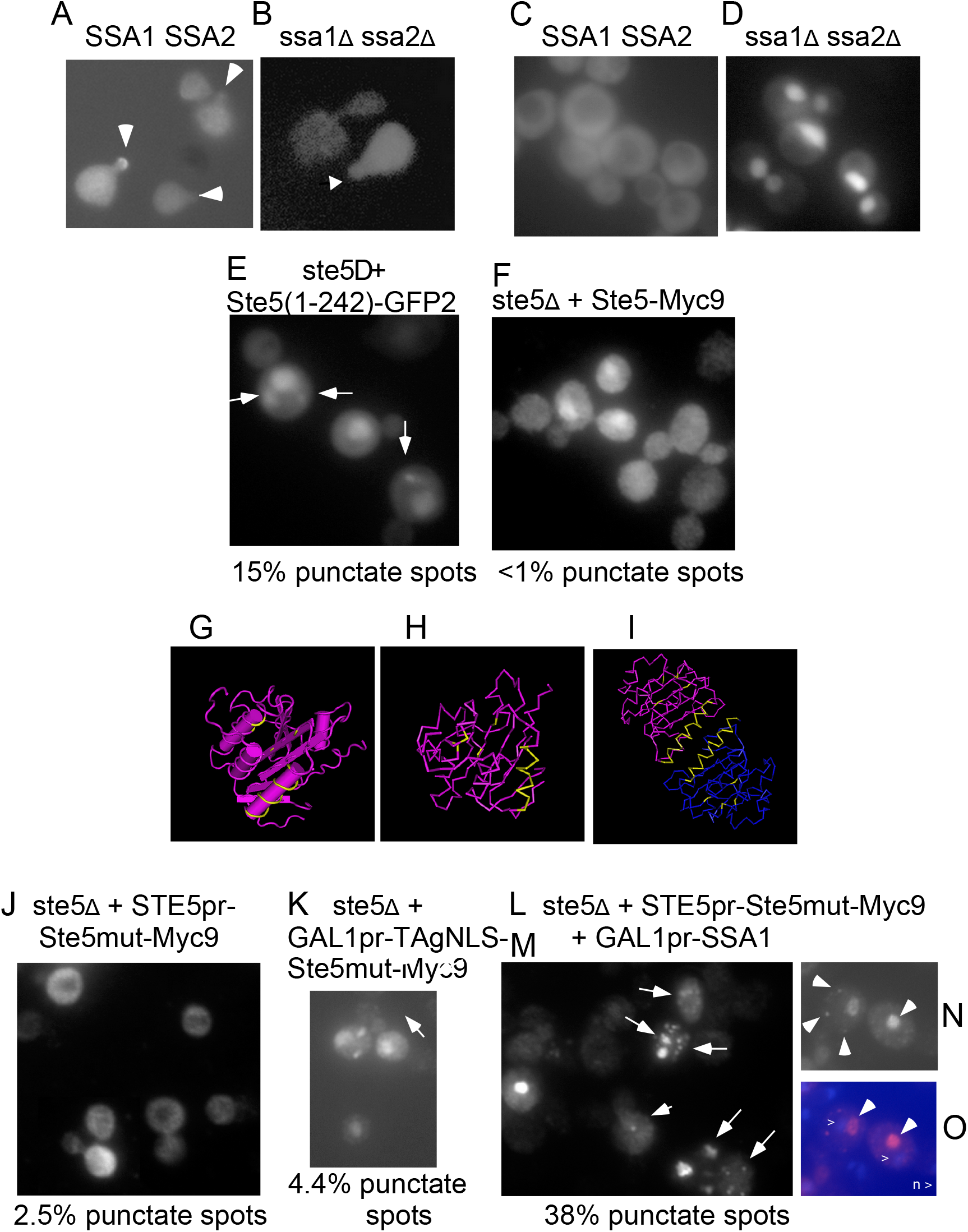
Ssa1 and Ssa2 alter the localization of wild-type and mutant Ste5. A-B. Live cell image of GFP-Ste5 in α factor-induced *SSA1D SSA2D* and *ssa1D ssa2D* cells. GFP-Ste5-(pSKM21) expressed from a *CEN* plasmid in *ssa1D ssa2D* cells has fewer and weaker GFP positive cells. Arrows indicate GFP-Ste5 accumulation at shmoo tips. C-D. Live cell images of Ssa1-GFP in *SSA1D SSA2D* and *ssa1D ssa2D* strains. E. *ste5D*+Ste5(1-242)-GFP2 room temperature. Arrows indicate punctate foci. F. *ste5Δ* + Ste5-Myc9. G-H Ste5-Ms 3FZE monomer crystal structure. I. Ste5 4F2H dimer crystal structure. The positions of the Ste5L610A/L614A/ L634A/L637A mutations within the VWA-like domain and the helix that dimerizes are yellow against the pink ribbon backbone (17, 27). See FigS1A for larger images. J. *ste5D* + ste5L610/614/634/637A-Myc9 (indicated as Ste5mut-Myc9), K. *ste5D*+ GAL1pr-TAgNLS-ste5L610/614/634/637A-Myc9 L-O. Overexpression of Ssa1 induces punctate foci of ste5L610/614/634/637A-Myc9. M. GAL1pr-SSA1 + Ste5mut-Myc9 clone 1, 9E10 indirect immunofluorescence. N. Same as M, Clone 2, 9E10. O. Same as N, DAPI stain. For L-O Arrows indicate punctate foci. *ste5D* strains with *ste5L610/614/634/637A-Myc9* and *GAL1promSSA1* were grown at 30°C in SC-2% dextrose selective medium then shifted to SC-2% galactose selective medium to induce Ssa1.

**Table 1.**
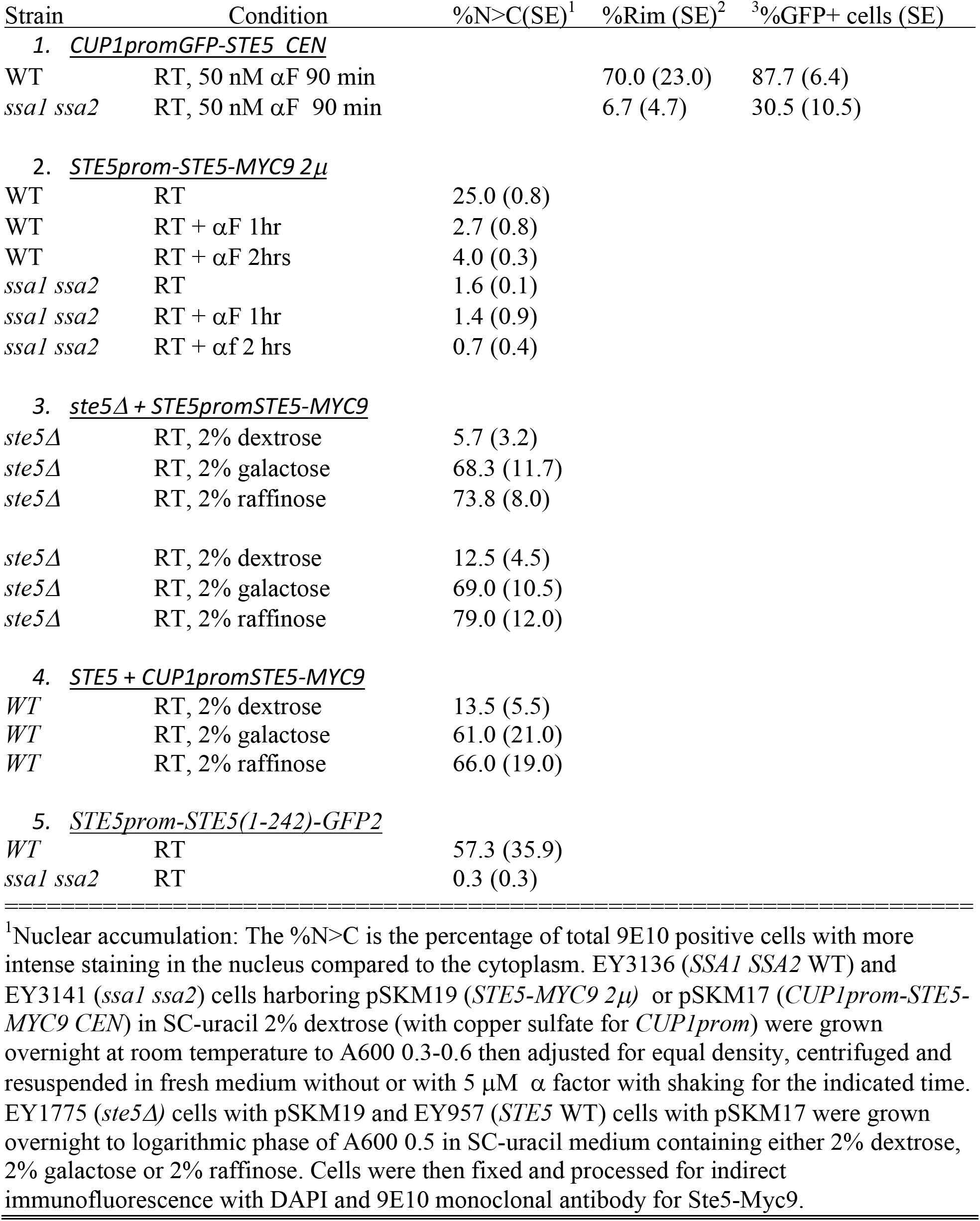
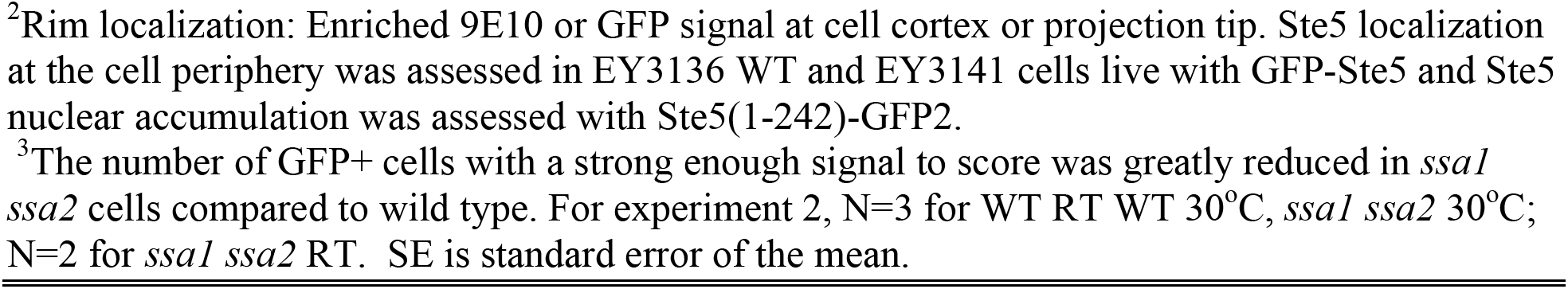
Localization of Ste5 to cell periphery and nucleus in wild type and *ssa1D ssa2D* cells.

| Strain | Condition | %N>C(SE)<sup>1</sup> | %Rim (SE)<sup>2</sup> | <sup>3</sup>%GFP+ cells (SE) |
| --- | --- | --- | --- | --- |
| **1. *CUP1promGFP-STE5 CEN*** |  |  |  |  |
| WT | RT, 50 nM αF 90 min | | 70.0 (23.0) | 87.7 (6.4) |
| *ssa1 ssa2* | RT, 50 nM αF 90 min | | 6.7 (4.7) | 30.5 (10.5) |
| **2. *STE5prom-STE5-MYC9 2μ*** |  |  |  |  |
| WT | RT | 25.0 (0.8) |  |  |
| WT | RT + αF 1hr | 2.7 (0.8) | | |
| WT | RT + αF 2hrs | 4.0 (0.3) | | |
| *ssa1 ssa2* | RT | 1.6 (0.1) |  |  |
| *ssa1 ssa2* | RT + αF 1hr | 1.4 (0.9) | | |
| *ssa1 ssa2* | RT + αf 2 hrs | 0.7 (0.4) | | |
| **3. *ste5Δ + STE5promSTE5-MYC9*** |  |  |  |  |
| *ste5Δ* | RT, 2% dextrose | 5.7 (3.2) |  |  |
| *ste5Δ* | RT, 2% galactose | 68.3 (11.7) |  |  |
| *ste5Δ* | RT, 2% raffinose | 73.8 (8.0) |  |  |
| *ste5Δ* | RT, 2% dextrose | 12.5 (4.5) |  |  |
| *ste5Δ* | RT, 2% galactose | 69.0 (10.5) |  |  |
| *ste5Δ* | RT, 2% raffinose | 79.0 (12.0) |  |  |
| **4. *STE5 + CUP1promSTE5-MYC9*** |  |  |  |  |
| WT | RT, 2% dextrose | 13.5 (5.5) |  |  |
| WT | RT, 2% galactose | 61.0 (21.0) |  |  |
| WT | RT, 2% raffinose | 66.0 (19.0) |  |  |
| **5. *STE5prom-STE5(1-242)-GFP2*** |  |  |  |  |
| WT | RT | 57.3 (35.9) |  |  |
| *ssa1 ssa2* | RT | 0.3 (0.3) |  |  |

<sup>1</sup>Nuclear accumulation: The %N>C is the percentage of total 9E10 positive cells with more intense staining in the nucleus compared to the cytoplasm. EY3136 (*SSA1 SSA2* WT) and EY3141 (*ssa1 ssa2*) cells harboring pSKM19 (*STE5-MYC9 2 $\mu$* ) or pSKM17 (*CUP1prom-STE5-MYC9 CEN*) in SC-uracil 2% dextrose (with copper sulfate for *CUP1prom*) were grown overnight at room temperature to A600 0.3-0.6 then adjusted for equal density, centrifuged and resuspended in fresh medium without or with 5 μM $\alpha$ factor with shaking for the indicated time. EY1775 (*ste5 $\Delta$* ) cells with pSKM19 and EY957 (*STE5* WT) cells with pSKM17 were grown overnight to logarithmic phase of A600 0.5 in SC-uracil medium containing either 2% dextrose, 2% galactose or 2% raffinose. Cells were then fixed and processed for indirect immunofluorescence with DAPI and 9E10 monoclonal antibody for Ste5-Myc9.
<sup>2</sup>Rim localization: Enriched 9E10 or GFP signal at cell cortex or projection tip. Ste5 localization at the cell periphery was assessed in EY3136 WT and EY3141 cells live with GFP-Ste5 and Ste5 nuclear accumulation was assessed with Ste5(1-242)-GFP2.
<sup>3</sup>The number of GFP+ cells with a strong enough signal to score was greatly reduced in *ssa1 ssa2* cells compared to wild type. For experiment 2, N=3 for WT RT WT 30°C, *ssa1 ssa2* 30°C; N=2 for *ssa1 ssa2* RT. SE is standard error of the mean.

### Ssa1 and Ssa2 enhance nuclear accumulation of Ste5

Ssa2-GFP is thought to localize in the cytoplasm and the nucleus whereas Ssa1-GFP is thought to localize only in the cytoplasm (94). We found Ssa1-GFP to localize throughout the cytoplasm and nucleus as shown by absence of nuclear exclusion (Fig 6B). Moreover, Ssa1-GFP was clearly enriched in the nucleus in addition to being in the cytoplasm in the *ssa1D ssa2D* strain that lacks endogenous Ssa1 and Ssa2 (Fig 6B). Therefore, Ssa1 is likely to have substrates in the cytoplasm and the nucleus.

Cytoplasmic Hsp70/Hsc70 proteins in yeast and mammals including Ssa1 and Ssa4 shuttle through the nucleus and are implicated in facilitating nuclear import of associated proteins (70, 95), although excess Ssa1 and Ssa2 inhibits nuclear accumulation of a substrate called Gts1 (96). We compared the localization of Ste5-Myc9 expressed at native levels in wild-type and *ssa1D ssa2D* strains and found that Ste5-Myc9 accumulated in fewer *ssa1D ssa2D* nuclei compared to wild-type before and during α factor treatment at room temperature (Table 1). Ste5(1-242)-GFP2 which has the NLS and RING-H2 also accumulated in fewer nuclei in *ssa1D ssa2D* cells (Table 1) whereas TAgNLS-GFP2 and TAgNLS-NES-GFP2 which have an additional NLS from SV40 TAg efficiently accumulated in nuclei of *ssa1D ssa2D* cells (Table S3). These results imply that Ssa1 and Ssa2 may stimulate nuclear accumulation or nuclear retention of Ste5 and may do so at least in part through the first 242 residues of Ste5.

Yeast cells grow most optimally at temperatures of 25°C (room temperature) to 30°C. The *S. cerevisiae* heat shock response is activated at temperatures of 37°C and higher with cell viability declining after long exposure to temperatures higher than 42°C (55). A heat shock at 55°C in the Craig lab ascospores induced cell death with no nuclear accumulation of Ste5-Myc9 in either the *SSA1 SSA2* or *ssa1D ssa2D* strains (Table 2), suggesting either that nuclear import was blocked or the nuclear pool was degraded or exported (Table 2).

**Table 2.** Effect of temperature on nuclear accumulation of Ste5-Myc9.

| Strain<sup>1</sup> | Temp | % N>C<sup>2</sup> | %N>C S.D |
| --- | --- | --- | --- |
| Strain *SSA1 SSA2* |  |  |  |
| + *STE5-MYC9* 2μ | RT 1-2 hr | 14,17,23 | 18 +/- 4.6 |
|  | 55°C 1 hr | 0,0, 0<sup>3</sup> | 0 +/- 0 |
| Strain *ssa1 ssa2* |  |  |  |
| + *STE5-MYC9* 2μ | RT 1-2 hr | 1,1,3 | 1.7 +/- 1.2 |
|  | 55°C 1 hr | 0,0, 0<sup>3</sup> | 0 +/- 0 |
| Strain *SSA1 SSA2 FY23*<sup>4</sup> |  |  |  |
| + *STE5-MYC9* 2μ | RT 3 hr | | 14.6 +/-2.8 |
|  | 30°C 3 hr |  | 21.4 +/-4.8 |
|  | 37°C 3 hr |  | 14.7 +/-4.6 |
|  | 42°C 3 hr |  | 18.9 +/-6.6 |
|  | 55°C 3 hr |  | 0 +/- 0<sup>3</sup> |
| Strain *cdc24-1 + CDC24* |  |  |  |
| + *STE5-MYC9* 2μ | RT 2-3 hr | 0.2, 2, 0.25, 2, 1.8 | 1.2 +/- 0.94 |
|  | 37°C 2-3 hr | 1, 6.8, 6, 1, 4 | 3.8 +/- 2.7 |
|  | 55°C 1-3 hr | 14, 24, 27, 17, 31 | 23 +/- 7.0 |

<sup>1</sup>Strains: *SSA1 SSA2* is EY3376 is EY3136 *ura3-* + (pSKM19 *STE5-MYC9 URA3* 2 $\mu$ ), *ssa1 ssa2* is EY3379 is EY3141 *ura3-ssa1::HIS3 ssa2::LEU2* + (pSKM19 *STE5-MYC9 URA3* 2 $\mu$ ). *SSA1 SSA2 FY23* + (pSKM19 *STE5-MYC9 URA3* 2 $\mu$ ), EYL2712 YMY467 (1,2,3) is EYL457 = YMY467 = [RAY914 *MATa cdc24-1::LoxPHIS5SpLoxP his3-200 leu3-3, 112 ura3-52 trp1-901 lys2-801 gal2 suc2-9* (pRS414 *TRP1-CEN-CDC24*)] + (pSKM10 *STE5-MYC9 URA3* 2 $\mu$ ). Cells were grown in SC-uracil with 2% dextrose to mid logarithmic phase (A600 0.4-0.6) then equal A600 units of cells were pelleted and resuspended in pre-warmed YEPD medium and incubated with shaking at the indicated temperature. Cells were collected and prepared for indirect immunofluorescence using 9E10 monoclonal antibody and DAPI.
<sup>2</sup>N>C means the 9E10 signal in the nucleus is stronger than in the cytoplasm. ~133-400 cells were counted for data points. The Ste5-Myc9 signal was weaker in the *ssa1 ssa2* strains than in the wild type strains.
<sup>3</sup>At 55°C (for 1-3 hours) cells of this strain background appear enlarged with a large vacuole devoid of Ste5-Myc9 signal and some Ste5-Myc9 signal beneath cell periphery. The cells appeared to be dying.
<sup>4</sup>A comparison of *FUS1prom-lacZ* $\beta$ galactosidase levels in S288c (FY23) grown at room temperature and 37°C was also done. Cells grown logarithmically at room temperature then exposed to + αF for 2 hours at room temperature had 135.6 +/- 1.0 S.E. FUS1-lacZ units whereas the same strain grown at 37°C for 3 hours then exposure to αF for 2 hours at 37°C had 122.5 +/- 9.0 S.E. FUS1-lacZ units.

In some S288c and W303a wild type strain backgrounds, elevated temperatures did induce more nuclear accumulation Ste5-Myc9. For example, a W303a background *MATa cdc24-1* (*CDC24-CEN-URA3)* strain from Rob Arkowitz accumulated more Ste5-Myc9 in nuclei at 37°C and 55°C compared to at room temperature (Table 2). More nuclear accumulation of Ste5(1-242)-GFP2 occurred in FY23 S288c cells grown at 37°C compared to at room temperature (nuclear accumulation at RT was 37.9% versus 80.4% at 37°C in one experiment and 69.7% versus 91.1% at 37°C in a second experiment). Thus, Ssa1 and Ssa2 together promote nuclear accumulation of Ste5 at room temperature and 30°C in the Craig strain background, consistent with less Ste5-Myc9 in 16,000 x g lysate pellets from *ssa1D ssa2D* cells compared to WT (Fig. 4A).

### Ste5 VWA domain mutant predicted to have greater disorder accumulates in punctate foci when Ssa1 is overexpressed

Proteins that are either soluble or aggregated with or without ubiquitinylation are corralled into distinct foci by Ssa1, Ssa2 and co-chaperones where they are refolded or routed for degradation in the vacuole or autophagy (65–67). We examined whether overexpression of Ssa1 altered the localization of wild type and mutant Ste5-Myc9 proteins that we first identified as having a propensity to form punctate foci. Wild type Ste5-Myc9 rarely localizes in inclusion bodies (i.e. punctate foci) by indirect immunofluorescence using an axioscope (Fig 6F, Table 3). The percentage of cells exhibiting punctate foci were: 0% in six independent experiments in W303, S288c backgrounds; and 0% and 1.5% after α factor treatment in two experiments in W303a background yielding an average of 0.19%. However, a Ste5(1-242)-GFP2 fusion that oligomerizes through the RING-H2 domain had more punctate foci in the cytoplasm (∼15% of cells) (Fig 6E). We screened several Ste5 mutants that might have less stable structures for greater propensity to form punctate foci. Interestingly, we found that a VWA domain quadruple point mutant, ste5L610/614/634/637A-Myc9 (Fig 6G-I, FigS2A-I), formed more punctate foci in the cytoplasm in ∼2.5% of cells at 30°C (Fig 6J). Overxpression of a TAgNLS derivative of this quadruple mutant, TAgNLS-ste5L610/614/634/637A-Myc9 (using the *GAL1* promoter) led to more punctate foci that were larger and more readily visible in 4% of cells and could be distinguished from the nuclear pool (Fig 6K). Thus, the TAgNLS-ste5L610/614/634/637A-Myc9 protein also forms foci outside of the nucleus.

**Table 3.**
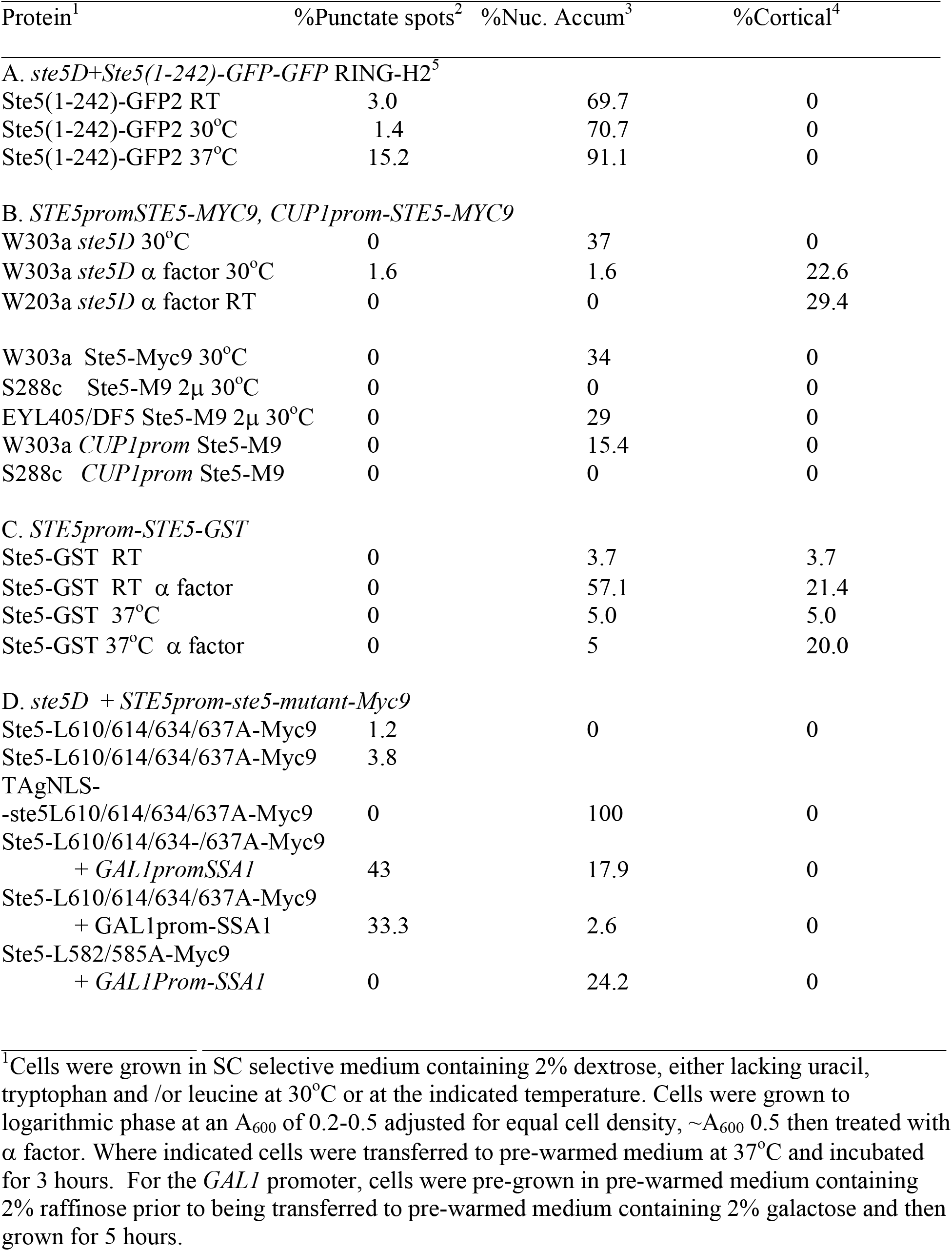
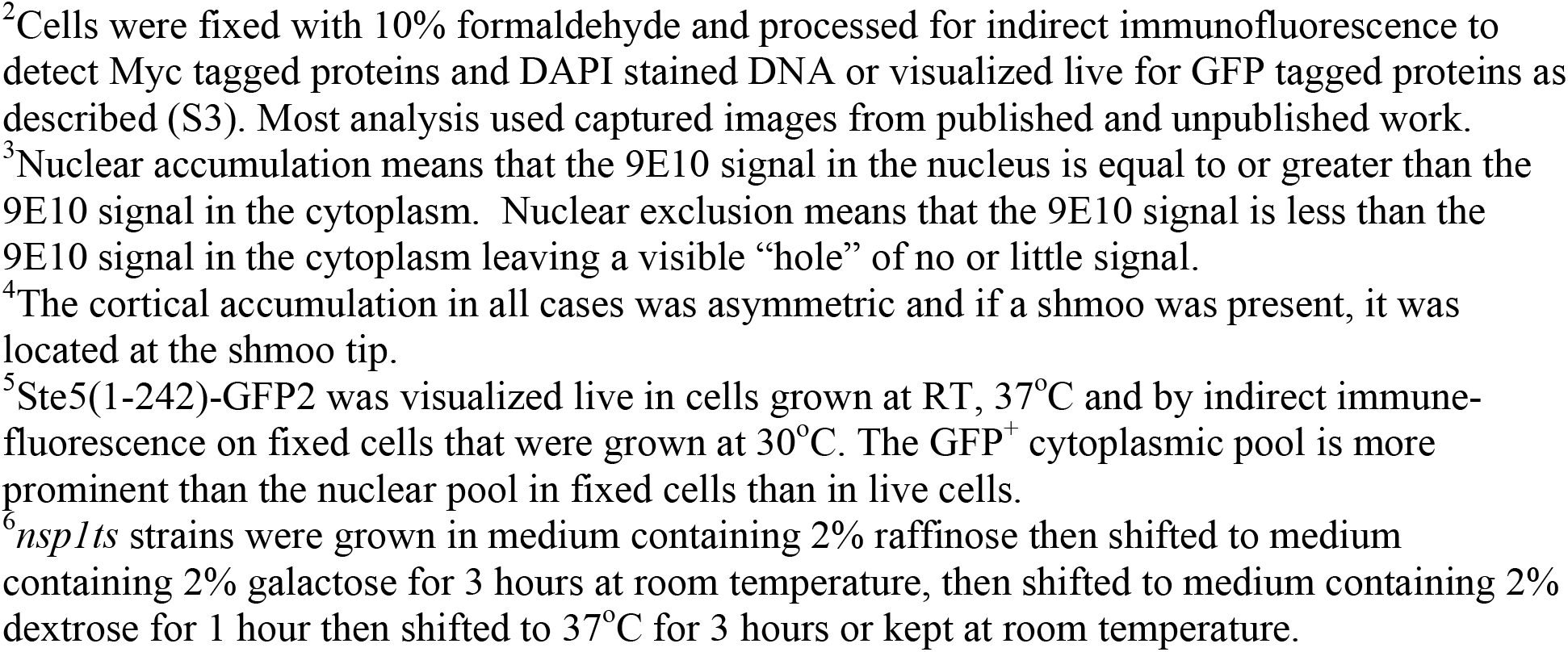
Prevalence of Ste5 in punctate foci.

| Protein<sup>1</sup> | %Punctate spots<sup>2</sup> | %Nuc. Accum<sup>3</sup> | %Cortical<sup>4</sup> |
| --- | --- | --- | --- |
| **A. *ste5D+Ste5(1-242)-GFP-GFP RING-H2*<sup>5</sup>** |  |  |  |
| Ste5(1-242)-GFP2 RT | 3.0 | 69.7 | 0 |
| Ste5(1-242)-GFP2 30°C | 1.4 | 70.7 | 0 |
| Ste5(1-242)-GFP2 37°C | 15.2 | 91.1 | 0 |
| **B. *STE5promSTE5-MYC9, CUP1prom-STE5-MYC9*** |  |  |  |
| W303a *ste5D* 30°C | 0 | 37 | 0 |
| W303a *ste5D* $\alpha$ factor 30°C | 1.6 | 1.6 | 22.6 |
| W203a *ste5D* $\alpha$ factor RT | 0 | 0 | 29.4 |
| W303a Ste5-Myc9 30°C | 0 | 34 | 0 |
| S288c Ste5-M9 2μ 30°C | 0 | 0 | 0 |
| EYL405/DF5 Ste5-M9 2μ 30°C | 0 | 29 | 0 |
| W303a *CUP1prom* Ste5-M9 | 0 | 15.4 | 0 |
| S288c *CUP1prom* Ste5-M9 | 0 | 0 | 0 |
| **C. *STE5prom-STE5-GST*** |  |  |  |
| Ste5-GST RT | 0 | 3.7 | 3.7 |
| Ste5-GST RT $\alpha$ factor | 0 | 57.1 | 21.4 |
| Ste5-GST 37°C | 0 | 5.0 | 5.0 |
| Ste5-GST 37°C $\alpha$ factor | 0 | 5 | 20.0 |
| **D. *ste5D + STE5prom-ste5-mutant-Myc9*** |  |  |  |
| Ste5-L610/614/634/637A-Myc9 | 1.2 | 0 | 0 |
| Ste5-L610/614/634/637A-Myc9 | 3.8 |  |  |
| TA<sub>g</sub>NLS- |  |  |  |
| -ste5L610/614/634/637A-Myc9 | 0 | 100 | 0 |
| Ste5-L610/614/634-/637A-Myc9 |  |  |  |
| + *GAL1promSSA1* | 43 | 17.9 | 0 |
| Ste5-L610/614/634/637A-Myc9 |  |  |  |
| + GAL1prom-SSA1 | 33.3 | 2.6 | 0 |
| Ste5-L582/585A-Myc9 |  |  |  |
| + *GAL1Prom-SSA1* | 0 | 24.2 | 0 |

<sup>1</sup>Cells were grown in SC selective medium containing 2% dextrose, either lacking uracil, tryptophan and /or leucine at 30°C or at the indicated temperature. Cells were grown to logarithmic phase at an A<sub>600</sub> of 0.2-0.5 adjusted for equal cell density, ~A<sub>600</sub> 0.5 then treated with $\alpha$ factor. Where indicated cells were transferred to pre-warmed medium at 37°C and incubated for 3 hours. For the *GAL1* promoter, cells were pre-grown in pre-warmed medium containing 2% raffinose prior to being transferred to pre-warmed medium containing 2% galactose and then grown for 5 hours.
<sup>2</sup>Cells were fixed with 10% formaldehyde and processed for indirect immunofluorescence to detect Myc tagged proteins and DAPI stained DNA or visualized live for GFP tagged proteins as described (S3). Most analysis used captured images from published and unpublished work.
<sup>3</sup>Nuclear accumulation means that the 9E10 signal in the nucleus is equal to or greater than the 9E10 signal in the cytoplasm. Nuclear exclusion means that the 9E10 signal is less than the 9E10 signal in the cytoplasm leaving a visible “hole” of no or little signal.
<sup>4</sup>The cortical accumulation in all cases was asymmetric and if a shmoo was present, it was located at the shmoo tip.
<sup>5</sup>Ste5(1-242)-GFP2 was visualized live in cells grown at RT, 37°C and by indirect immunofluorescence on fixed cells that were grown at 30°C. The GFP<sup>+</sup> cytoplasmic pool is more prominent than the nuclear pool in fixed cells than in live cells.
<sup>6</sup>*nsp1ts* strains were grown in medium containing 2% raffinose then shifted to medium containing 2% galactose for 3 hours at room temperature, then shifted to medium containing 2% dextrose for 1 hour then shifted to 37°C for 3 hours or kept at room temperature.

We overexpressed Ssa1 from the *GAL1* promoter to determine whether it affects Ste5 localization or formation of punctate foci. No obvious effect on punctate foci was found for wild type Ste5-Myc9. However, long exposures of films from the Ssa1-GFP and Ste5-Myc9 co-immunoprecipitation suggests that increased levels of Ssa1-GFP widens the width of the phosphorylated Ste5-Myc9 band, increases the amount of the high molecular weight species likely to be ubiquitinylated and/or aggregated and produces fragmented species not detected with the control GFP-Cdc24 fusion (Fig S3A, compare Lanes 1,2 with 3,4). Notably, when *STE5prom-ste5L610/614/634/637A-*was co-expressed with the *GAL1prom-SSA1* gene, the ste5L610/614/634/637A-Myc9 protein accumulated in many punctate foci and larger patches of varying sizes in ∼38% of cells compared to 2.6% of cells without excess Ssa1, as well as in nuclei in ∼15% of cells (Fig 6L-M, Table 3). Co-staining with DAPI revealed the punctate foci were not overlapping the nuclei (FigS3B-E). A second experiment with a different transformant confirmed that punctate foci did not overlap DAPI stained nuclear DNA and appear to be cytoplasmic (Fig 6N-O). By contrast, overexpression of Ssa1 did not induce either wild type Ste5-Myc9 or Ste5(1-242)-GFP2 to form more punctate foci. The effect of Ssa1 inducing the localization of ste5L610/614/634/637A-Myc9 into punctate foci is a hallmark of Hsp70 chaperones moving misfolded proteins into inclusion bodies for refolding or degradation (40,65–67). These data provide additional evidence of a role for Ssa1 in Ste5 homeostasis.

Ssa1 and Ssa2 bind misfolded proteins on hydrophobic patches of certain characteristics that flip to the aqueous surface. We therefore used bioinformatics to examine whether the L610A, L614A, L634A and L637A mutations are predicted to influence either the folding of Ste5 or the number of potential Ssa1 binding sites. The four mutations fall in evolutionarily conserved residues in Ste5 within the VWA-like domain and nearby oligomerization domain (Fig S2K) suggesting their effects on Ste5 have general implications. Atomic interaction predictions suggest all four alanine substitutions reduce intramolecular interactions (Fig S2F-I) that might normally stabilize the protein. The alanine substitutions in Ste5 are less hydrophobic and less bulky than leucines (FigS4A-C) and are predicted to be less buried (Fig S4D-E). The region overlapping the Ste5L610/614/634/637A mutations is predicted to have slightly different crystallization properties than wild type Ste5 based on XtalPred analysis (Table 4) and to have somewhat higher tendency for disorder based on IUPRED and IUPRED2/ANCHOR analysis; Fig S4F-J).

**Table 4.**
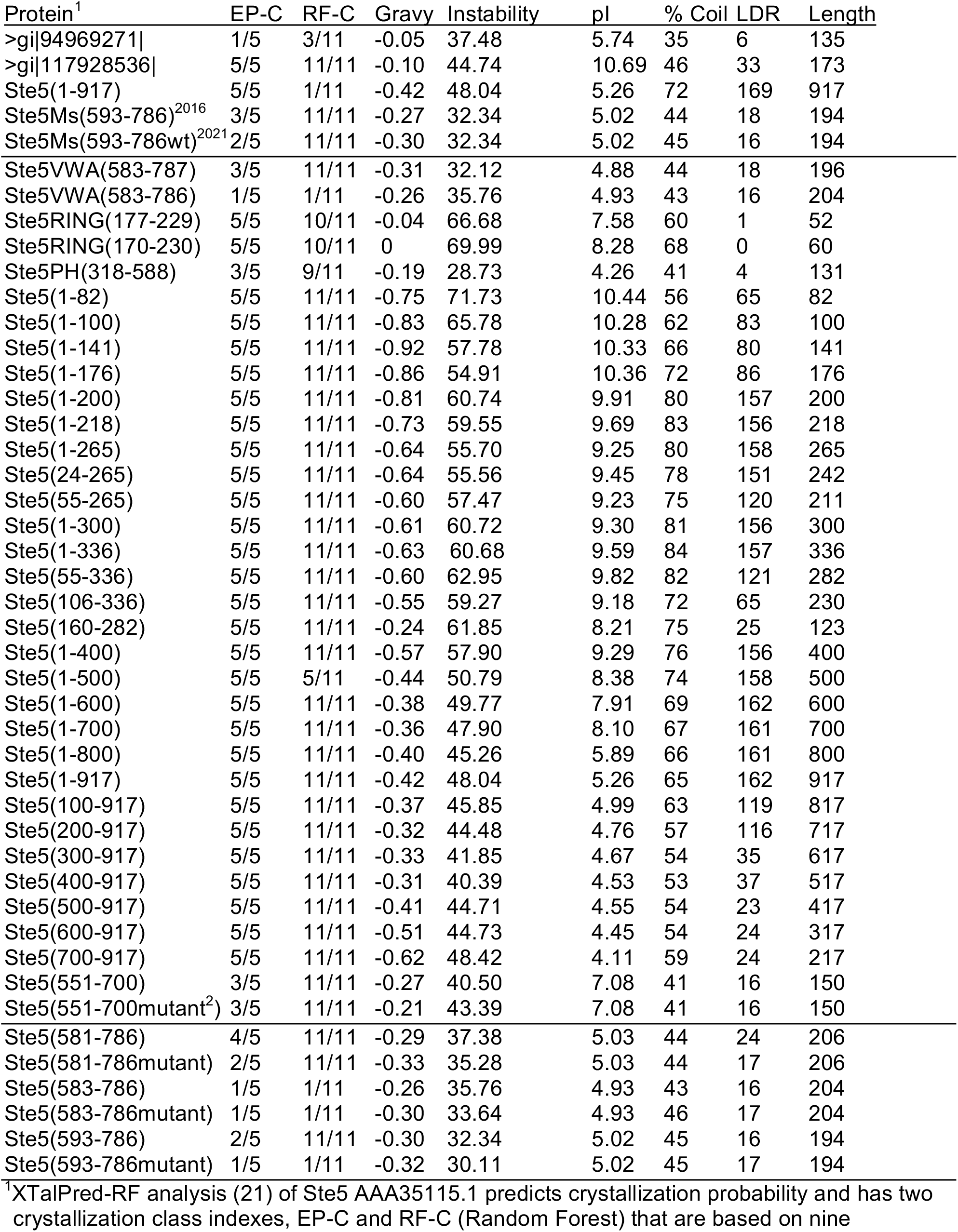
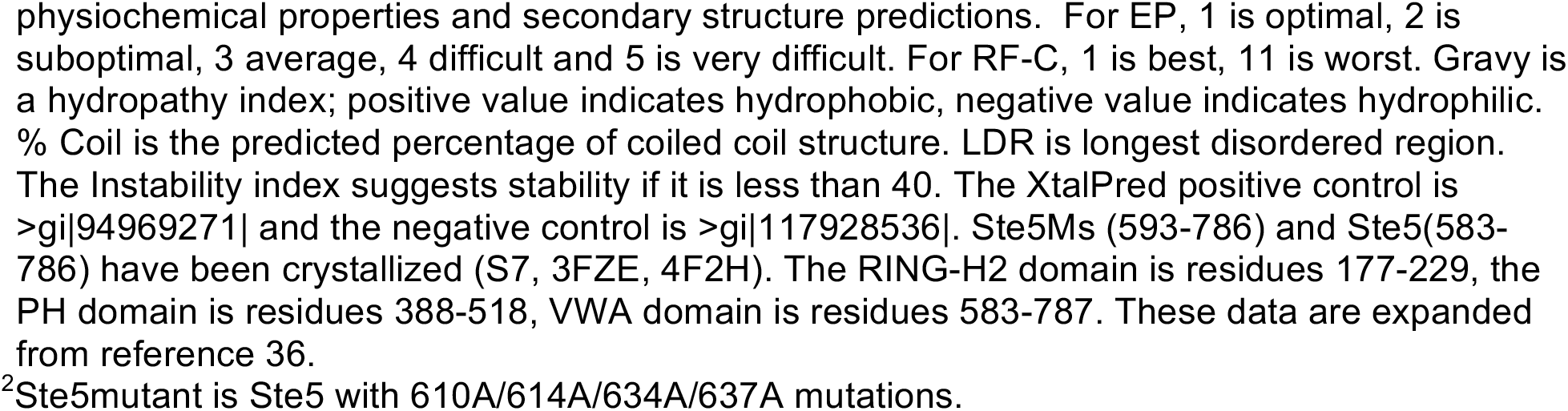
XtalPred analysis of Ste5 fragments.

| Protein<sup>1</sup> | EP-C | RF-C | Gravy | Instability | pI | % Coil | LDR | Length |
| --- | --- | --- | --- | --- | --- | --- | --- | --- |
| >gi\|94969271\| | 1/5 | 3/11 | -0.05 | 37.48 | 5.74 | 35 | 6 | 135 |
| >gi\|117928536\| | 5/5 | 11/11 | -0.10 | 44.74 | 10.69 | 46 | 33 | 173 |
| Ste5(1-917) | 5/5 | 1/11 | -0.42 | 48.04 | 5.26 | 72 | 169 | 917 |
| Ste5Ms(593-786)<sup>2016</sup> | 3/5 | 11/11 | -0.27 | 32.34 | 5.02 | 44 | 18 | 194 |
| Ste5Ms(593-786wt)<sup>2021</sup> | 2/5 | 11/11 | -0.30 | 32.34 | 5.02 | 45 | 16 | 194 |
| Ste5VWA(583-787) | 3/5 | 11/11 | -0.31 | 32.12 | 4.88 | 44 | 18 | 196 |
| Ste5VWA(583-786) | 1/5 | 1/11 | -0.26 | 35.76 | 4.93 | 43 | 16 | 204 |
| Ste5RING(177-229) | 5/5 | 10/11 | -0.04 | 66.68 | 7.58 | 60 | 1 | 52 |
| Ste5RING(170-230) | 5/5 | 10/11 | 0 | 69.99 | 8.28 | 68 | 0 | 60 |
| Ste5PH(318-588) | 3/5 | 9/11 | -0.19 | 28.73 | 4.26 | 41 | 4 | 131 |
| Ste5(1-82) | 5/5 | 11/11 | -0.75 | 71.73 | 10.44 | 56 | 65 | 82 |
| Ste5(1-100) | 5/5 | 11/11 | -0.83 | 65.78 | 10.28 | 62 | 83 | 100 |
| Ste5(1-141) | 5/5 | 11/11 | -0.92 | 57.78 | 10.33 | 66 | 80 | 141 |
| Ste5(1-176) | 5/5 | 11/11 | -0.86 | 54.91 | 10.36 | 72 | 86 | 176 |
| Ste5(1-200) | 5/5 | 11/11 | -0.81 | 60.74 | 9.91 | 80 | 157 | 200 |
| Ste5(1-218) | 5/5 | 11/11 | -0.73 | 59.55 | 9.69 | 83 | 156 | 218 |
| Ste5(1-265) | 5/5 | 11/11 | -0.64 | 55.70 | 9.25 | 80 | 158 | 265 |
| Ste5(24-265) | 5/5 | 11/11 | -0.64 | 55.56 | 9.45 | 78 | 151 | 242 |
| Ste5(55-265) | 5/5 | 11/11 | -0.60 | 57.47 | 9.23 | 75 | 120 | 211 |
| Ste5(1-300) | 5/5 | 11/11 | -0.61 | 60.72 | 9.30 | 81 | 156 | 300 |
| Ste5(1-336) | 5/5 | 11/11 | -0.63 | 60.68 | 9.59 | 84 | 157 | 336 |
| Ste5(55-336) | 5/5 | 11/11 | -0.60 | 62.95 | 9.82 | 82 | 121 | 282 |
| Ste5(106-336) | 5/5 | 11/11 | -0.55 | 59.27 | 9.18 | 72 | 65 | 230 |
| Ste5(160-282) | 5/5 | 11/11 | -0.24 | 61.85 | 8.21 | 75 | 25 | 123 |
| Ste5(1-400) | 5/5 | 11/11 | -0.57 | 57.90 | 9.29 | 76 | 156 | 400 |
| Ste5(1-500) | 5/5 | 5/11 | -0.44 | 50.79 | 8.38 | 74 | 158 | 500 |
| Ste5(1-600) | 5/5 | 11/11 | -0.38 | 49.77 | 7.91 | 69 | 162 | 600 |
| Ste5(1-700) | 5/5 | 11/11 | -0.36 | 47.90 | 8.10 | 67 | 161 | 700 |
| Ste5(1-800) | 5/5 | 11/11 | -0.40 | 45.26 | 5.89 | 66 | 161 | 800 |
| Ste5(1-917) | 5/5 | 11/11 | -0.42 | 48.04 | 5.26 | 65 | 162 | 917 |
| Ste5(100-917) | 5/5 | 11/11 | -0.37 | 45.85 | 4.99 | 63 | 119 | 817 |
| Ste5(200-917) | 5/5 | 11/11 | -0.32 | 44.48 | 4.76 | 57 | 116 | 717 |
| Ste5(300-917) | 5/5 | 11/11 | -0.33 | 41.85 | 4.67 | 54 | 35 | 617 |
| Ste5(400-917) | 5/5 | 11/11 | -0.31 | 40.39 | 4.53 | 53 | 37 | 517 |
| Ste5(500-917) | 5/5 | 11/11 | -0.41 | 44.71 | 4.55 | 54 | 23 | 417 |
| Ste5(600-917) | 5/5 | 11/11 | -0.51 | 44.73 | 4.45 | 54 | 24 | 317 |
| Ste5(700-917) | 5/5 | 11/11 | -0.62 | 48.42 | 4.11 | 59 | 24 | 217 |
| Ste5(551-700) | 3/5 | 11/11 | -0.27 | 40.50 | 7.08 | 41 | 16 | 150 |
| Ste5(551-700mutant<sup>2</sup>) | 3/5 | 11/11 | -0.21 | 43.39 | 7.08 | 41 | 16 | 150 |
| Ste5(581-786) | 4/5 | 11/11 | -0.29 | 37.38 | 5.03 | 44 | 24 | 206 |
| Ste5(581-786mutant) | 2/5 | 11/11 | -0.33 | 35.28 | 5.03 | 44 | 17 | 206 |
| Ste5(583-786) | 1/5 | 1/11 | -0.26 | 35.76 | 4.93 | 43 | 16 | 204 |
| Ste5(583-786mutant) | 1/5 | 1/11 | -0.30 | 33.64 | 4.93 | 46 | 17 | 204 |
| Ste5(593-786) | 2/5 | 11/11 | -0.30 | 32.34 | 5.02 | 45 | 16 | 194 |
| Ste5(593-786mutant) | 1/5 | 1/11 | -0.32 | 30.11 | 5.02 | 45 | 17 | 194 |

<sup>1</sup>XTalPred-RF analysis (21) of Ste5 AAA35115.1 predicts crystallization probability and has two crystallization class indexes, EP-C and RF-C (Random Forest) that are based on nine
physiochemical properties and secondary structure predictions. For EP, 1 is optimal, 2 is suboptimal, 3 average, 4 difficult and 5 is very difficult. For RF-C, 1 is best, 11 is worst. Gravy is a hydropathy index; positive value indicates hydrophobic, negative value indicates hydrophilic. % Coil is the predicted percentage of coiled coil structure. LDR is longest disordered region. The Instability index suggests stability if it is less than 40. The XtalPred positive control is >gi|94969271| and the negative control is >gi|117928536|. Ste5Ms (593-786) and Ste5(583-786) have been crystallized (S7, 3FZE, 4F2H). The RING-H2 domain is residues 177-229, the PH domain is residues 388-518, VWA domain is residues 583-787. These data are expanded from reference 36.
<sup>2</sup>Ste5mutant is Ste5 with 610A/614A/634A/637A mutations.

Hsp70 proteins, including Ssa1, recognize client proteins through short hydrophobic segments of 5-7 amino acids flanked by positively charged amino acids (53) which typically occur every 30-40 amino acids (97). BiPPred analysis has successfully predicted Hsp70 Kar2 binding sites (97) and a Ssa1 binding site in a peptide (97, 98). Interestingly, BiPPred predicted 13 fewer highly optimal recognition sequences for Hsp70 on the Ste5L610/614/634/637A polypeptide compared to wild type Ste5, which had 107 sites with highly optimal recognition scores of 0.9-1.0; (FigS4L). These observations are certainly consistent with a role for Ssa1 in Ste5 homeostasis and raise the possibility that Ste5L610/614/627/634 has less stable regions that are not as well recognized by Hsp70 chaperones for folding.

### Ssa1 and Ssa2 are required for vegetative growth and α factor-induced shmoo formation

To assay the contribution of Ssa1 and Ssa2 to mating responses, we constructed single, double and triple *ssa* mutants by sporulating a diploid from the Craig laboratory with heterozygous *ssa1, ssa2, ssa3* and *ssa4* null mutations (Table S1). The *ssa1*, *ssa2* single mutants as well as *ssa1 ssa3 ssa4* and *ssa2 ssa3 ssa4* triple mutants had only slightly slower growth than wild-type at room temperature (∼25°C) and 30°C. The *ssa1* and *ssa2* single mutants had colony sizes that were slightly smaller than wild type (Fig 7A, *ssa1*: 53% wt colony size N=22, *ssa2*:77% N=20). By contrast, the *ssa1D ssa2D* double mutant grew poorly at room temperature and 30°C and barely grew at all at 37°C (Fig 8D, Table S4). Spontaneous suppressors that permitted growth arose after 5 days incubation at 37°C at a rate of 10^-7^ similar to what has been found (83, 84,106,107). The *ssa1D ssa2D ssa4* triple and *ssa1D ssa2D ssa3 ssa4* quadruple mutants were inviable and died as unbudded haplospores as previously found. The *ssa1* and *ssa2* single mutants gave rise to colonies that were 1.96-fold and 2.75-fold larger, respectively, than the *ssa1D ssa2D* double mutant colonies on YEPD plates (Fig 7A, *ssa1D ssa2D* 28% wt colony size, N=25), indicating that the two mutations had additive deleterious effects on growth. The *ssa1D ssa2D* double mutants routinely yielded mainly small and a few bigger colonies in streakouts (in Fig 7A, the size range for *ssa1D ssa2D* is 0.12 to 0.47 compared to 0.71-1.3 for *SSA1D SSA2D*), but the small and big colonies grew at similar rates in liquid culture, suggesting varying delays in resuming growth on solid support.

**FIG 7.**
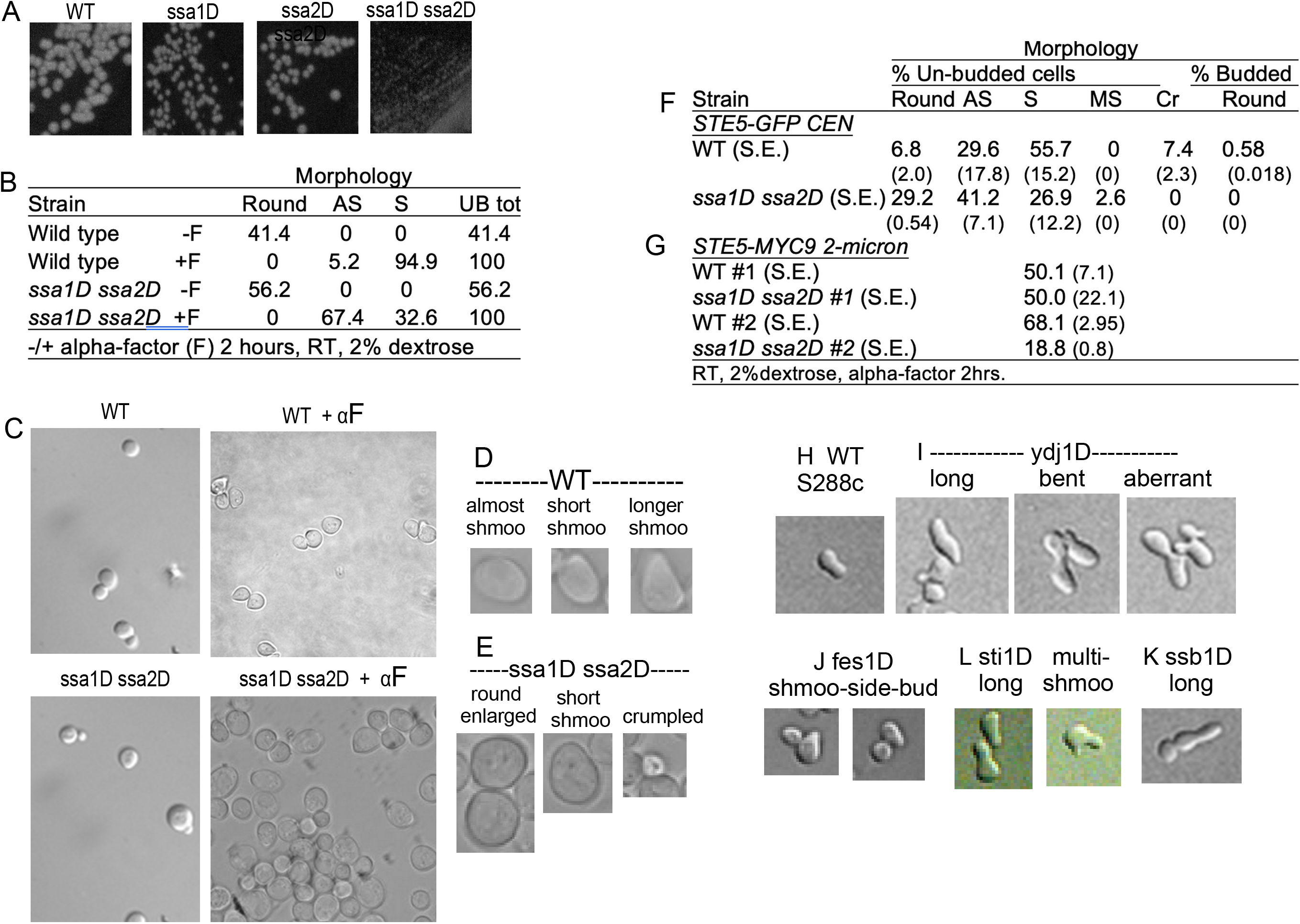
*ssa1D ssa2D* double mutants grow more slowly and are defective in shmoo formation. A. Streakouts of wild-type, *ssa1*, *ssa2*, *ssa1D ssa2D* strains. EY3136, EY3137, EY3138, EY3141 were streaked on a YEPD plate grown at room temperature and photographed after 3 days. B. Tally of morphology of WT and *ssa1D ssa2D* strains during vegetative growth and after α factor treatment (+αF) at room temperature. C. Images of wild-type and *ssa1D ssa2D* cells during logarithmic growth and after treatment with α factor. The field of α factor treated *ssa1D ssa2D* cells has 6 shmoos out of a total of 80 cells. D-E. Examples of wild-type and *ssa1D ssa2D* shmoo morphologies. F-G. Tally of morphologies of WT and *ssa1D ssa2D* cells with extra Ste5 (*STE5-GFP-CEN* and *STE5-Myc9-2μ).* H-K. Images of WT (H) and morphologically aberrant shmoos from Hsp70 network null mutants *ydj1D* (I), *fes1D* (J), *ssb1D* (K) and *sti1D* (L). Cells were grown logarithmically in YEPD at room temperature and treated with 5 μM α factor for 90 minutes.

**FIG 8.**
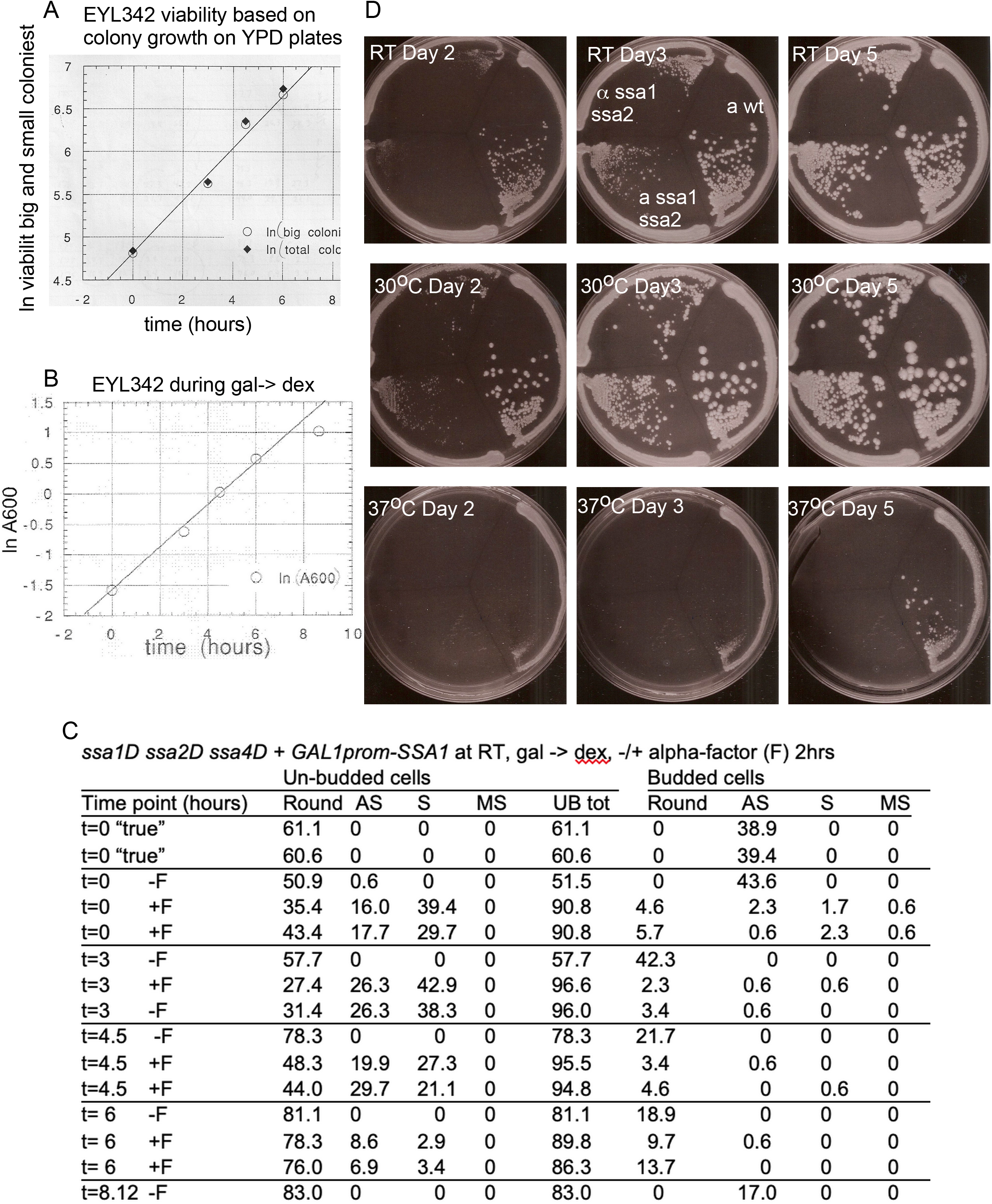
Effect of depletion of Ssa1 in a *ssa1D ssa2D ssa4* triple mutant on cell morphology. A. Liquid growth of small and big colonies of Craig lab strain EYL342 *MATa ssa1::HIS3 ssa2::LEU2 ssa4::LYS2 + pGAL1prom-SSA1* at 30°C in YEP-2%galactose medium. B. Growth rate of EYL342 during depletion of Ssa1. EYL342 was grown at 30°C overnight in YEP-2% galactose to logarithmic phase, washed once with YEP-2% dextrose (pH 5.0), then resuspended in same medium and shaken at 30°C without or with 5μM α factor for 2 hours, measured by growth by A600 and fixed at time intervals with formaldehyde and tallied for cell number after sonication. D. Tally of morphology of EYL342 during depletion of Ssa1. Cells were grown as in B and scored (N=175) as unbudded, budded, round, enlarged, almost like a shmoo (AS), pear-shaped shmoo (S), shmoo with two or more projections (MS, multi-shmoo). D. Growth of *WT* and *ssa1D ssa2D* strains at room temperature, 30°C, and 37°C. Strains EY3141 and EY3136 were streaked onto YEPD plates and incubated at different temperatures. Plates were photographed over the course of 5 days.

Stronger mating pathway activation is needed for shmoo formation than for G1 arrest (84). The morphologies of *ssa1D ssa2D* cells in vegetative growth were similar to wild-type but the cells appeared slightly larger and more were unbudded (e.g. 41.4% unbudded *SSA1D SSA2D* cells versus 56.2% unbudded *ssa1D ssa2D* cells with *ssa1D ssa2D* cells and ∼ 10% wider (Fig 7B-C, Table S5). The *ssa1D ssa2D* cells had a weaker morphological response to α factor than wild type *SSA1D SSA2D* cells. Although the *ssa1D ssa2D* double mutant cells could arrest in G1 phase, fewer cells formed shmoos compared to wild type (Fig 7B-G; FigS7). Most noteworthy was that many more *ssa1D ssa2D* cells were round enlarged rather than the classic pear shape with tapered projection. The cells that had some shmoo morphology had less emerged shorter projections that were broader (e.g. Fig 7C, Table S9, 29.2% round unbudded *ssa1D ssa2D* cells versus 6.8% *SSA1D SSA2D* cells. In Fig 7B, the cell width for *SSA1D SSA2D* in arbitrary units is 1 (size range 0.89-1.2, N=10) versus 1.43 for *ssa1D ssa2D* (size range 0.97-1.9, N=17). Therefore, Ssa1 and Ssa2 are needed for efficient α factor-induced polarized morphogenesis in addition to vegetative growth. In the absence of Ssa1 and Ssa2, more cells expand growth in all directions rather than towards a single polarization site.

We also looked at cells solely dependent on Ssa1 for survival for their ability to arrest in G1 phase and form shmoos. Ssa1 was overexpressed in a *ssa1D ssa2D ssa4* strain with a *GAL1prom-SSA1-CEN* (EYL342) by growth in medium containing 2% galactose, then Ssa1 was depleted through glucose repression and then cells were exposed to α factor or mock buffer and examined over time (Fig 8A-C). The cells were mainly unbudded upon Ssa1 depletion both in the absence and presence of α factor. At the earliest time point, 39.4%/29.7% of the cells could form shmoos, however, only 2.9%/3.4% formed shmoos after 6 hours of depletion of Ssa1. There was no loss of viability during the course of the experiment (Fig. 8B). Thus, Ssa1 is needed to transit from G1 to S phase, to bud and to form shmoos during mating.

### Ssa1 and Ssa2 are required for α factor-induced expression of FUS1::ubiYlacZ, growth inhibition and activation of MAPKs Fus3 and Kss1

To examine the ability of α factor to induce mating pathway transcription we monitored the expression of a *FUS1::ubiYlacZ* reporter gene that has a short half-life (88, 89). Ssa1 and Ssa2 provided additive contributions to the basal levels of FUS1:UbiYlacZ with bigger reduction in β-galactosidase activity in the *ssa1D ssa2D* double mutant (Fig 9A-B; *ssa1*: 3.6% WT, *ssa2*: 14% WT, *ssa1D ssa2D*: ∼1.6% WT). In the presence of α factor for 2 hours, FUS1::ubiYlacZ β-galactosidase activity was reduced in the *ssa1* single mutant to ∼46-55% of wild type, reduced in the *ssa2* single mutant to ∼46-63% of wild-type, and further reduced in the *ssa1D ssa2D* double mutant to ∼24% of wild type at all concentrations of α factor tested (Fig 9A-B). An *ssa1 ssa3 ssa4* triple mutant solely dependent on Ssa2 had 3.4% of wild type basal FUS1::ubiYlacZ activity and, after α factor stimulation, had ∼45-49% wild type FUS1::ubiYlacZ activity (Fig 9B; lack of viability prevented analysis of *ssa2 ssa3 ssa4* triple and a *ssa1D ssa2D ssa3 ssa4* quadruple mutants). Therefore, Ssa1 and Ssa2 are equivalently required for full expression of FUS1:UbiYlacZ, with little contribution from Ssa3 and Ssa4.

**FIG 9.**
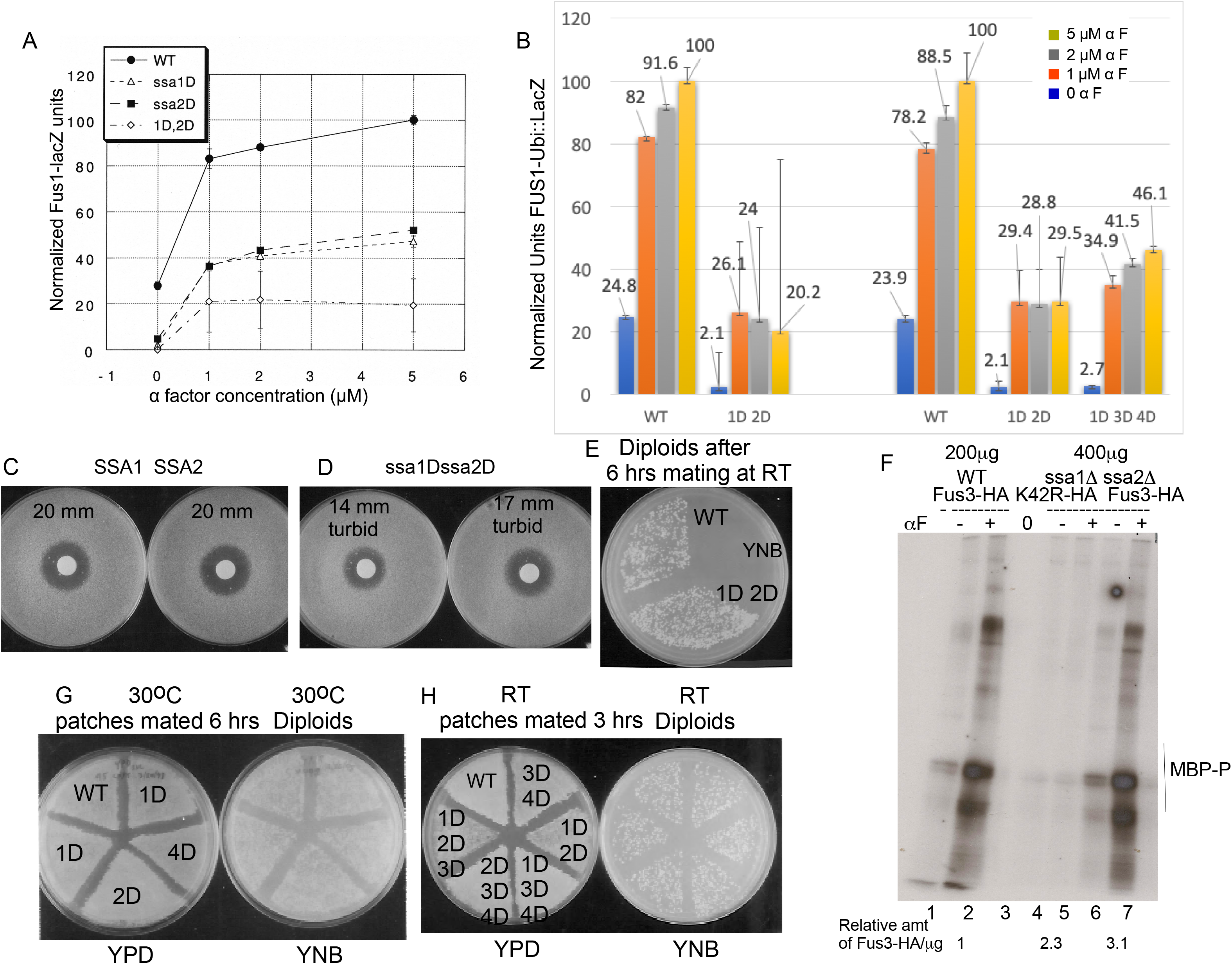
*ssa1D ssa2D* double mutants are defective in FUS1::ubiYlacZ activity and G1 arrest but can still mate. A-B. Quantification of β-galactosidase activity in wild-type and *ssa1*, *ssa2*, *ssa1D ssa2D* and *ssa1 ssa3 ssa4* strains expressing FUS1::ubiYlacZ. EY3136, EY3137, EY3138, EY3141 and EY3148 (and non reverting *ura3^-^*derivatives) harboring the FUS1::ubiYlacZ plasmid pDL1460 were grown at room temperature and exposed to the indicated amount of α factor for 2 hours. Extracts were prepared as described in Materials and Methods. C-D. Duplicate halo assays for WT (C) and *ssa1D ssa2D* (D) strains done at RT. E. Mating of wild type and *ssa1D ssa2D* cells. F. Fus3 kinase assay. A MATa bar1 *ssa1D ssa2D* strain was assessed for FUS3-HA and fus3K42R-HA activity. 400 μg of cell extract was used for the bar1 *ssa1D ssa2D* strain and 200 μg cell extract was used for the *bar1* strain. - /+ is 50 nM α factor for 60 minutes. MBP is myelin basic protein exogenously added substrate. Levels of Fus3-HA are in Fig 10D. –G-H. *ssa1*, *ssa2*, *ssa3*, and *ssa4* single, double and triple deletion mutants can mate. Patches of mutants on YEP-2% dextrose agarose plates were mated against lawns of *MATa lys9* wild-type cells for the indicated time and then transferred to YNB 2% dextrose medium agarose plates to select for prototrophs. Testing of individual colonies showed they were diploids and not haploid suppressors. E. 30oC, 6 hour mating for WT, *ssa1*, *ssa2*, *ssa4*. F. RT, 3 hour mating for WT, *ssa1D ssa2D*, *ssa3 ssa4*, *ssa1 ssa3 ssa4*, *ssa2 ssa3 ssa4*, *ssa1D ssa2D ssa3*. G RT, 6 hour mating for *WT, ssa2 ssa2*.

To assess ability to cell cycle arrest in presence of α factor, we performed halo assays. The *ssa1D ssa2D* strain was also less efficient at being inhibited by α factor in halo assays and formed smaller more turbid halos of growth inhibition compared to wild type (Fig 9C-D). Collectively, the reduced shmoo response, FUS1::ubiYlacZ activity and G1 arrest reveal that mating pathway activation is impaired in the *ssa1D ssa2D* double mutant.

We assessed the ability of α factor activated Fus3 to phosphorylate associated substrates in vitro, by compareing Fus3-HA activity in immunoprecipitation kinase assays of extracts from *bar1 SSA1D SSA2D* cells to those of *bar1 ssa1D ssa2D* cells that had been exposed to 50 nM α factor for one hour. The profile of Fus3-HA phosphorylated co-immunoprecipitated substrates and exogenously added myelin basic protein (MBP) was similar for both strains (Fig 9F) and was dependent on Fus3 catalytic activity (Fig.9 fus3K42R-HA, lanes 4,5). Twice as much *ssa1D ssa2D* WCE was immunoprecipitated and the level of the Fus3-HA protein was 3-fold higher in the *ssa1D ssa2D* extracts compared to wild type (Fig. 9F; TableS2). Therefore, ∼6-fold less Fus3-HA activity was detected *in vitro* in the *ssa1D ssa2D* extracts. Thus, less of the pool of Fus3-HA enzyme is active in the *ssa1D ssa2D* strain.

We assayed activation of Fus3 and Kss1 with an anti-active p44/p42 MAPK antibody made against a conserved peptide in human ERK2 in BAR1+ *ssa1D ssa2D* and BAR1+ strains (Fig 10A). Basal levels of active Fus3 and Kss1 were not detected in this Craig lab strain background compared to W303a for our standard amount of whole cell extract. Strikingly, the levels of α factor-induced active Fus3 and Kss1 were reduced in a *ssa1D ssa2D* mutant during vegetative growth and at all time points after addition of 50nM α factor (Fig 10A, Table S2). The decrease in the band signal was not from less protein loaded based on ribosomal protein Tcm1 and Ponceau S staining (data not shown). Analysis of more *ssa1D ssa2D* cell extract did not reveal basal Fus3-P, whereas basal Kss1-P was detected in both wild type and *ssa1D ssa2D* whole cell extracts (Fig 10C). Densitometry of these representative immunoblots confirmed that the *ssa1D ssa2D* strain had undetectable basal levels of active Fus3 and less active Fus3 during α factor stimulation compared to wild type (Table S2). Kss1 activation during α factor stimulation was reduced, although less impaired than that of Fus3 in the *ssa1D ssa2D* strain (Fig 10B-C), but the abundance of Kss1 was not reduced (Fig 10E, Table S2).

**FIG 10.**
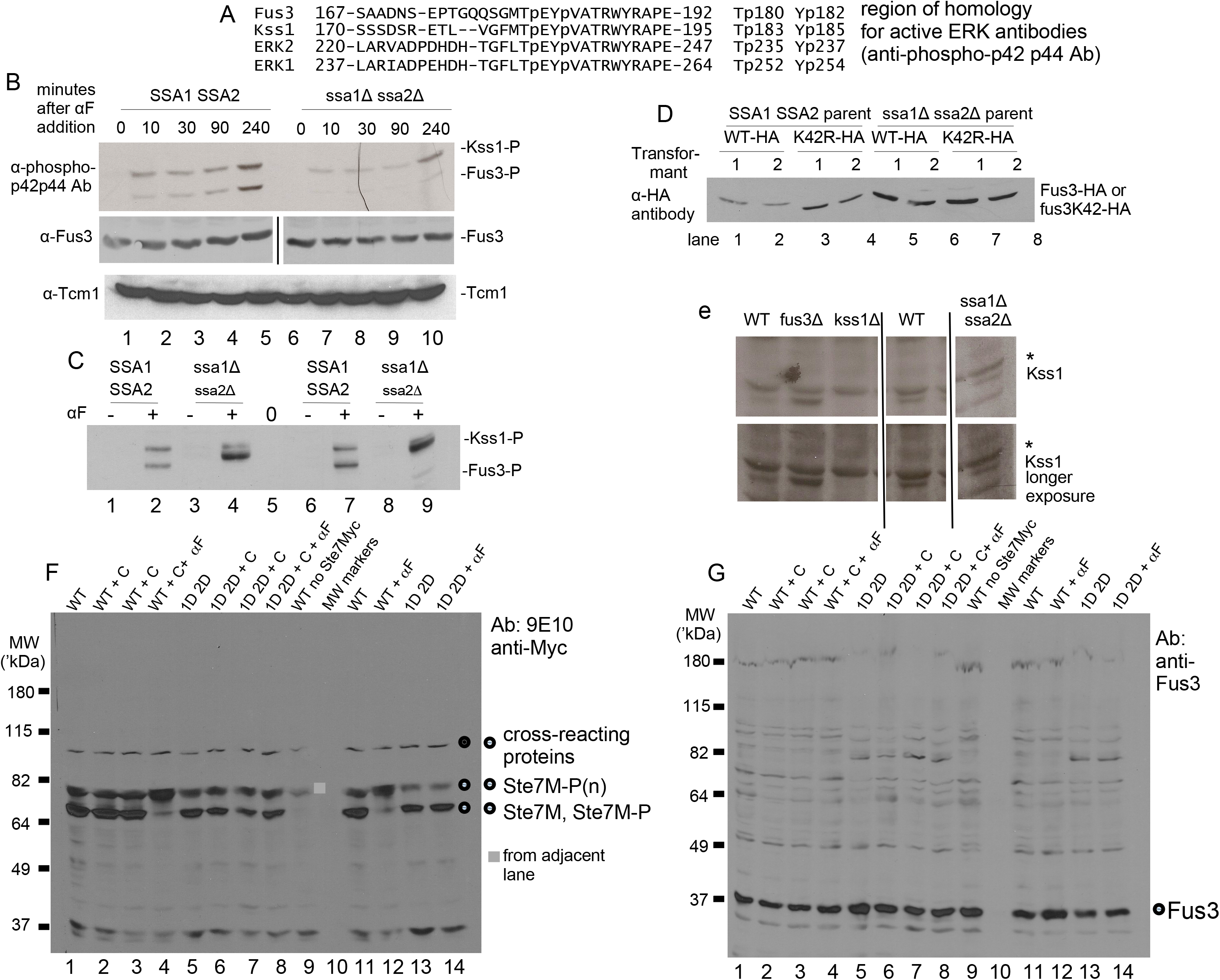
Effect of *ssa1D ssa2D* mutations on Fus3 and Kss1 activity, Ste7 feedback phosphorylation and abundance of Fus3, Kss1, Ste7 and Tcm1. A. Amino acid alignment of Fus3 and Kss1 with the Erk1, Erk2 region used to prepare anti-phospho-p42, p44 antibodies. B. Fus3 and Kss1 activation are reduced in a *ssa1D ssa2D* strain. Immunoblot of WCEs from wild-type (EY3136) and *ssa1D ssa2D* (EY3141) exposed to 2.5 μM α factor for the indicated length of time. Active Fus3 and Kss1 were detected with anti-phospho-p42p44 antibody. The immunoblot was stripped and reprobed with Tcm1 monoclonal antibody. Fus3 was assessed separately using anti-Fus3 polyclonal antibodies. C. Immunoblot of large amounts of WCE to detect basal Fus3 and Kss1 activation. Lanes 1-4 have 150 μg of WCE, lanes 5-8 have 300 μg WCE. The WCEs were precipitated with trichloroacetic acid to concentrate. D. Abundance of Fus3-HA and fus3K42R-HA in EY3136 and EY3141 harboring *FUS3-HA-CEN* (pYEE102) and *fus3K42R-HA-CEN* (pYEE106) plasmids. Two transformants are shown, 12CA5 was used for detection. Equal total protein was loaded on the gel. E. Abundance of Kss1 in EY3136 WT, EY3141 *ssa1D ssa2D*, EY*fus3* and *kss1* strains. Kss1 was detected with anti-Kss1 polyclonal 6775-kss1-yc-19a. The asterisks indicate cross-reacting proteins. All lanes are from the same gel. F-G. Abundance of hypo- and hyper-phosphorylated Ste7 and Fus3. Wild-type and *ssa1D ssa2D* cells harboring *CYC1-STE7-MYC-CEN* were grown to mid-logarithmic phase then treated with 10 μg/ml cycloheximide for 10 minutes where indicated (+). 1 mg of WCEs were concentrated by 40% ammonium precipitation for analysis. Blots were probed with 9E10 (G) and anti-Fus3 polyclonal antibodies (H).

Ste7-Myc is hyperphosphorylated in the presence of α factor through feedback phosphorylation by Fus3 and Kss1. We looked at basal feedback phosphorylation of Ste7-Myc by Fus3 and Kss1 using a *CYC1prom-STE7-MYC* gene that is not regulated by either the mating pathway or heat shock proteins Ssa1, Ssa2, Hsf1. The ratio of phosphorylated Ste7-Myc to hypophosphorylated Ste7-Myc was nearly identical in wild-type and *ssa1D ssa2D* strains with or without cycloheximide to reveal primary signaling events (Fig 10F, Table S2, slower mobility protein band is compared to faster mobility phosphorylated protein bands). Kss1 is likely responsible for most of the basal feedback phosphorylation of Ste7-Myc in the *ssa1D ssa2D* strain, since we only detect Kss1 as basally active (Fig, 10B,E). Thus, Ssa1 and Ssa2 are essential for basal activation of Fus3 and full basal activation of Kss1 and essential for α factor-induced activation of both Fus3 and Kss1. Ssa1 and Ssa2 are therefore required for efficient response to mating pheromone in addition to regulating Ste5 abundance and integrity.

### Abundance of Fus3, Ste7 and Kss1 in *ssa1D ssa2D* mutant

The large effect of the *ssa1* and *ssa2* mutations on Ste5 abundance was specific to Ste5, based on comparative analysis of Fus3, Ste7, Kss1 and Tcm1. The abundance of Fus3 was not obviously reduced in the *ssa1D ssa2D* double mutant (Fig 4C, Fig 10 B,D, TableS2) even when Ste5 was overexpressed (Table S2). Fus3 abundance appeared to be approximately the same or slightly higher in a *ssa1D ssa2D* double mutant compared to wild-type (TableS2). The level of Fus3 appeared elevated in a *ssa1D ssa2D* double mutant when Fus3 was catalytically inactive (i.e. Fus3K42R-HA and Fus3-HA (Fig 10D). Fus3 was slightly elevated in a *GAL1prom-STE5-MYC9* shut-off experiment (data not shown), and when the *ssa1D ssa2D* strain co-expressed a *CYC1prom-STE7-MYC* gene, either before and after blocking translation with cycloheximide (Fig 10G, TableS2). The increase in endogenous Fus3 ranged from 1.1 to 1.3-fold for nontagged Fus3 to several-fold for HA-tagged Fus3 (TableS2) Thus, Ssa1 and Ssa2 have an inhibitory effect on Fus3 abundance that is independent of Fus3 catalytic activity. This is consistent with unconfirmed high throughput data suggesting Ssa1 may associate with Fus3 and suggests the Hsp70 proteins may promote its degradation. An increase in abundance of Fus3 in the *ssa1D ssa2D* double mutant might counteract somewhat a reduction in the level of Ste5 protein.

The abundance of Kss1 was also slightly elevated in the *ssa1D ssa2D* double mutant to ∼1.24 the level in the isogenic wild type strain. A *fus3D* control strain had a larger 2.59-fold increase in Kss1, consistent with the known repressive effect of Fus3 on Kss1 expression (Fig 10E, Table S2). By contrast, Ste7-Myc abundance was not obviously altered in the *ssa1D ssa2D* mutant, (Fig 10F, Table S2). The relative abundance of phosphorylated and unphosphorylated species of Ste7-Myc was the same in wild type and *ssa1D ssa2D* strains (Fig 10E, Table S2, *ssa1D ssa2D/*WT Ste7-Myc 1.02 +/- 0.14 s.d., Ste7-Myc-P 1.0 +/- 0 s.d.). Collectively, these findings suggest that Ssa1 and Ssa2 negatively regulate the abundance of Fus3 and potentially may also negatively regulate Kss1 abundance (high background from the anti-Kss1 antibodies interfered with densitometry on replicates). By contrast Ssa1 and Ssa2 do not appear to regulate Ste7-Myc in an obvious way, although minor changes would not be obvious by immunoblot analysis.

### The *ssa1D ssa2D* double mutant can mate

We compared the mating of wild-type, *ssa1*, *ssa2*, *ssa3*, *ssa4* single, double and triple mutants in a standard plate assay in which patches of cells are mated to a lawn of wild-type cells. Surprisingly, there was no obvious decrease in mating ability for any of the mutants compared to wild-type at room temperature or 30°C in *BAR1* or *bar1* backgrounds (Fig 8D, Table S4). Given the poor growth, reduced Ste5 abundance, MAPK activation and morphogenesis in the *ssa1D ssa2D* double mutant, the *ssa1 and ssa2* mutations must induce additional changes that compensate for the signaling defects. For example, Ssb1 and Ssb2 are predicted to interact with Ste5 (99) and may generate a pool of Ste5 that is folded and functional in the *ssa1D ssa2D* strain along with other proteins. In addition, counteractive effects of increased Fus3 abundance may relieve the reduction in the level of Ste5. In addition, genetic analysis of interspecies chimeras suggests that weaker binding of Ste5 to kinases may enhance mating (100).

### Interaction data suggest Ssa1 is the chaperone most dedicated to the mating pathway

To determine which Hsp70 proteins are likely to be important for mating pathway and invasive growth pathway signaling, we looked at the distribution of chaperone network interactions with ∼227 mating and invasive growth pathway and polarity proteins (TableS5) using published interaction data (TableS5). We included 12 Hsp70 proteins (9 cytosolic, 2 mitochondrial, 1 ER), 2 Hsp110 proteins, 2 Hsp40/DnaJ family proteins (Ydj1, Sis1), 2 nucleotide exchange factors (Fes1/HspBP1 homolog, Snl1/Bag-1 homolog), 2 Hsp70-associated disaggregation chaperones (Hsp104, Hsp42), an Hsp90 co-chaperone (Cdc37), and Sti1, a co-chaperone that bridges Hsp70and Hsp90) and for comparison 2 Hsp90 chaperones (Hsc82, Hsp80). Most data came from unconfirmed high throughput experiments. All cells in the published experiments had been grown at either RT or 30°C in logarithmic phase in either YEPD, SCD or SILAC medium and were not subjected to heat shock (TableS5).

Ssa1 is predicted to have the largest number of mating pathway clients among the classic Hsp70 proteins (Ssa1,2,3,4) and may regulate over 79% of the proteins examined; however, only a small number have been identified from α factor treatment experiments (TableS6, FigS5). Ssa2 exhibits substantial overlap and is predicted to regulate 55% of the proteins compared to only 13% for Ssa3 and 14% for Ssa4. Interestingly, 89% of proteins predicted to be regulated by Ssa2 are also predicted to be regulated by Ssa1, and likewise for Ssa3 (84%) and Ssa4 (86%) (FigS5A). A similar pattern was found for invasive growth pathway proteins (FigS5B). The bias towards Ssa1 is even more pronounced among core mating pathway signaling components (95%) and core invasive growth pathway signaling components (94%) (FigS5C-D). The Hsp110 protein Sse1 which cooperates with Hsp70 and Hsp90 proteins (46) is predicted to interact with many components in the mating and invasive growth pathways (50%, 57% respectively) (Table S6). None of the other regulators we examined are predicted to interact with a significant number of proteins in the mating and invasive growth pathways (Table S6).

### Mutations in Ssa1 and Ssa2 regulators Fes1, Ydj1 and Sse2 cause unique defects in shmoo morphogenesis

To explore compensatory functions that might regulate in parallel with Ssa1 and Ssa2, we determined whether mutations in other Hsp70 network proteins affect the ability of cells to respond to α factor and arrest in G1 phase and form shmoos. A *fes1* mutant (39, 40) was defective in shmoo formation and the few shmoos that formed often had a small bud on the side (Fig 7J, Table S6,S7, Fig S6). Mutant *ydj1* cells (101) did not efficiently arrest in G1 phase and were often severely misshapen (both unbudded and budded cells) and formed fewer shmoos that were often enlarged and aberrant, with projections that were longer, wider, curved or bent (Fig 7I, FigS6E-F, Table S6, Tables S6, S8), adding to known defects in Axl1 biogenesis and a factor processing (102). The Sse1 Hsp110 protein (40, 103) is predicted to interact with many mating and invasive growth proteins, whereas its paralog Sse2 is predicted to interact with few (Tables S6). Surprisingly, the *sse1* mutant had a nearly normal pheromone response for G1 arrest and shmoo morphogenesis, whereas the *sse2* mutant failed to form shmoos although it could undergo G1 arrest and cell enlargement (Fig S7, Table S8). Perhaps its paralog Sse2 compensates in the *sse1* mutant but not *vice versa*. Mutation of Ssz1 Hsp70 that modulates Zuo1, a ribosome-associated J protein, caused defects in both G1 arrest and shmoo formation (Fig S7, Table S8). Therefore, Ssz1 may provide unique functions for mating morphogenesis although it is not predicted to interact with many mating pathway proteins (Table S6). Surprisingly, the *ssb1* mutant underwent nearly normal G1 arrest and shmoo formation with some shmoos longer than wild-type (Fig 7K; Table S8), contrasting its predicted interactions with many mating pathway proteins (Table S6). Perhaps its paralog Ssb2 compensates for loss of Ssb1. Interestingly, the *sti1* mutation in the Hsp90 cochaperone (104) increased both the number of shmoos and the length of shmoo projections compared to wild type (Fig 7G, Fig S7, Table S8), indicating either enhanced shmoo formation or reduced downregulation of shmoo formation. Currently, Sti1 has no known interactions with mating proteins (Table S6). In summary, Fes1, Ydj1, Sse2, and Ssz1 are needed for proper shmoo morphogenesis with their loss causing distinct changes in morphology, whereas Sti1 may inhibit shmoo formation. Thus, the Hsp70 network provides additional redundant and unique contributions to the mating response together with Ssa1 and Ssa2.

## Discussion

Faulty removal of damaged proteins that arise from misfolding, chemical insults, aggregation and translational truncations disrupts cell function and promotes aging and disease. We have found Ssa1 and Ssa2 to have key roles in regulating Ste5 integrity (Fig 2). Ssa1 co-purifies with Ste5 (Fig 1) and this association can be confirmed using tags on either end of Ste5 and Ssa1 in co-immunoprecipitations (Fig 2). The large amount of Ssa1 associating with Ste5 is likely due to its large size and many potential Ssa1 binding sites based on BIPPRed analysis (FigS4L). Ssa1 is in high abundance compared to Ste5 and MAPK pathway signaling components based on mean averages of molecules per cell from quantitative mass spectrometry studies (i.e. Ssa1-45137 to 314830, Ste5-814, Ste7-615, Ste11-1023, Ste20-3959, Fus3-5006, Kss1-3653 (105)). Loss of Ssa1 and Ssa2 is detrimental to Ste5-Myc9 integrity and results in over 2-fold reduction in abundance (Fig 3), greater vulnerability to proteolysis at all temperatures examined and during α factor stimulation (Fig 3,4) and greater availability of epitope-tags on the N- and C-termini to monoclonal antibodies (Fig 5). By contrast, Ste7, Fus3, Kss1, Tcm1 and many cross-reacting proteins to antibodies showed little (e.g. 1.1-1.3-fold increase for Fus3, slight increase for Kss1) or no change (i.e. Ste7, Tcm1) in abundance in the *ssa1D ssa2D* double mutant. Ste5 normally accumulates largely as monomers in the nucleus and as dimers at the cell cortex. Ste5 localization was impaired in the *ssa1D ssa2D* mutant: less Ste5-Myc9 and several other derivatives accumulated at the cell cortex (Fig 6, TableS3), suggesting the capacity of dimers of Ste5 to bind to anchors at the plasma membrane is impaired. Less Ste5 accumulated in nuclei including Ste5-Myc9, Ste5(1-242)-GFP2, which has the bipartite NLS (Table 1), GFP-Ste5 and TAgNLSK128T-Ste5-Myc9 (Table S3). We infer from these findings that Ssa1 and Ssa2 chaperone functions may increase directly or indirectly the pool of Ste5 competent to enter the nucleus or protect and sequester a nuclear pool from either export or degradation by SCF^Cdc4^ in the nucleus. Since the cytoplasmic pool of Ste5-Myc9 was also reduced in the *ssa1 ssa2* strain, Ssa1 and possibly Ssa2 also appear to protect Ste5 from degradation in the cytoplasm. Further work is needed to understand how the chaperones protect Ste5 from degradation. Moreover, overexpression of Ssa1 induced a VWA-like domain mutant Ste5L610/614/634/637-Myc9 to localize in numerous punctate foci and patches in the cytoplasm (Fig 6; Table 3) supporting a role for Ssa1 in quality control of Ste5. Further work is needed to define these foci and their function. Collectively, multiple lines of evidence support an active role for Ssa1 in maintaining Ste5 abundance, protection from potentially less functional conformers and degradation and proper localization in the cell.

Ssa1/Ssa2 exert refolding and aggregation functions in the cytoplasm which produces either soluble biologically active protein or, conversely, soluble misfolded protein that can be recognized for ubiquitinylation and degradation (39-41,63,65,67,73). The simplest interpretation is that Ste5 is protected by Ssa1 and possibly Ssa2 to stay properly folded for biological activity. This function of the chaperones may block unwanted degradation of Ste5 in the cytoplasm and the nucleus and potentially influence accessibility to binding partners. On the other hand, the more misfolded conformers or those molecules regulated to be turned over are expected to be targeted for degradation either by the proteasome and/or through the vacuole. It is tempting to speculate that Ssa1 (and possibly Ssa2) may also positively regulate Ste5 degradation by the SCF^Cdc4^ ubiquitin ligase complex in the nucleus after it has been phosphorylated by Cln/Cdc28. However, such a function would be expected to yield an increase in Ste5 abundance in the *ssa1 ssa2* double mutant, and we currently lack this type of evidence. On the other hand, we repeatedly found that Ssa1 and Ssa2 promoted overexpressed Ste5 to accumulate as high molecular weight species that are known to be ubiquitinylated (Fig. 2; 37). Thus, we do not rule out the possibility and think further work would be needed to determine whether Ssa1 and or Ssa2 have direct or indirect roles in promoting ubiquitin-dependent degradation of Ste5

Ssa1 and Ssa2 are known to sequester misfolded and aggregated proteins into a variety of inclusion bodies. The Ste5L610/614/634/637A-Myc9 foci/inclusion bodies induced by Ssa1 appear to be in the cytoplasm and near the nucleus (Fig 6, FigS2, Table 4). The conditions of vegetative growth, cytoplasmic location and shapes of the Ste5L610/614/634/637A-Myc9 inclusion bodies tend to rule out IPOD, cytoplasmic granules (106), and aggresomes (107), but they could conceivably be JUNQ/INQ (65, 67), Q-bodies/CytoQ bodies (107), stress foci (67) or sequestrosomes (60). We compared localization data for heat shock proteins, aggregated proteins, various inclusion bodies, and proteins that are markers for organelles and vesicles. This analysis suggests that the pattern of foci for Ste5L610/614/634/637A-Myc9 foci is most similar to aggregated Ste11DNK444R (66, 106, 108), and punctate foci of misfolded Ssa1-GFP and von Hippel-Landau protein (VHL-GFP) that accumulate in *hsp104* and *hsp82* mutants (108, 109) and Q bodies visualized with Ssa1-GFP, Hsp104-mCherry and Hsp82-GFP (41). Further work is needed to classify the Ste5L610/614/634/637A-Myc9 foci and determine their function.

The published high throughput data support potential interaction between Fus3 and Ssa1, but not with Kss1 or Ste7, although Kss1 is predicted to interact with Sse2 which binds Ssa1 and Ssa2 (Table S7, S8). We found that Fus3 and Kss1 may be inhibited by Ssa1 and/or Ssa2, because their abundance was slightly elevated 10-30% in the *ssa1D ssa2D* strain. Ste7 and Tcm1 appeared unaffected (Fig 8, Table S2), but minor effects might not be detected. The nearly equimolar amounts of Ste5-Myc9 and Ssa1 in the co-ips and reduced MAPK Fus3 activation raises the possibility that Ssa1 regulates Ste5 while in complex with additional signaling components. Further work may distinguish whether Ssa1 regulates the conversion of Ste5 into forms that can bind signaling partners or directly regulates Ste5-kinase signaling complexes and whether Ssa1 and or Ssa2 promote degradation of Fus3 and Kss1..

Of the 25 HSP70 network proteins examined for potential interactions in published databases, we found a strong bias towards Ssa1 followed by Ssa2 and Sse1 among mating and invasive growth proteins. Ssa1 is predicted to regulate over 79% of the 227 proteins examined and 95% of core signaling proteins compared to ∼55% for Ssa2 and Sse1, 13% for Ssa3 and 14% for Ssa4, with similar trends for invasive growth (Table S6, FigS5).

Despite the numerous proposed mating pathway targets for the Ssa proteins, we found, surprisingly, that mating still occurs in all *ssa1-ssa4* combinations tested (Fig 7, Table S6). Therefore, compensatory mechanisms permit mating during reduced signaling and loss of Ssa quality control. How might loss of Ssa1 and Ssa2 be compensated? One possibility is that a threshold level of active Ste5 conformers permits mating, perhaps with folding assistance from Ssb1 and Ssb2 during translation; similar reasoning can be applied to other proteins. Ste5 is very rate-limiting and low levels can be sufficient for function (9). Perhaps unfolding leads to a pool of active conformers of Ste5. Perhaps other mating pathway proteins are derepressed, expressed at higher levels (e.g. Fus3) or have fewer inhibitors. Additionally, genetic analysis suggests weakened binding of Ste5 to kinase may enhance mating (100). In addition, stress induced from changes in cell wall integrity from *ssa1D ssa2D* mutations may trigger activation of the invasive growth pathway to activate Kss1 which can partially substitute for Fus3 (Fig 1C). Perhaps the unfolded response (UPR) becomes activated triggering the cell surface mucin Msb2, which stimulates signaling through Ste11 to Kss1 (110). Redundant chaperone folding systems may provide compensatory functions that promote mating, such as compensatory increases in Ssa4, Hsf1 and Msn2/Msn4 that occur in *ssa* mutants although the current data on viability would argue against this (40), or compensatory changes in ubiquitin-dependent degradation (44,64,68). Notably, we found that Fes1, Ydj1, Sse2, and Ssz1 were needed for shmoo morphogenesis in addition to Ssa1 and Ssa2, and that Sti1 may inhibit shmoo formation (Tables S9, S9), suggesting multiple compensatory mechanisms preserve mating during normal growth and stress. Fes1 and both Sse1/Sse2 are also needed for invasive growth “mat” formation (111), indicating functional overlap for mating and invasive growth.

## Supporting information

SupportingInformationText

## Acknowledgments

We greatly thank Yuanyi Feng (Professor, Uniformed Services University of the Health Sciences, Bethesda, MD) and Yunmei Wang (Professor, Case Western Reserve School of Medicine, Cleveland, OH) for research contributions when they were in the lab and William S. Lane, John Neveu for the mass spectrometry (Harvard Microchemistry/Mass Spectrometry and Proteomics Resource Laboratory of Harvard University), Maosong Qi (Physician, Lowell General Hospital, MA) for teaching and doing the kinase assays with R.M., and Yunmei and Bill for helpful comments. We thank E. Craig (U. Wisconsin), B. Errede (U. North Carolina, Chapel Hill), P. Pryciak (U. Massachusetts Worcester Medical School), A. Truman (U. North Carolina, Charlotte), A. Kumar (U. Michigan), Z. Moqtaderi and K. Struhl (Dept. BCMP, HMS) for strains, plasmids, protocol details or archived published data. The authors declare they have no conflicts of interest with the contents of this manuscript. This work was supported by a grant from the N.I.H. (GM46962). A portion of the methods are published in a thesis by Homayoun Sadeghi in the field of Biology for a Master of Liberal Arts in Extension Studies from Harvard Extension School, June 1999. Y.F., Y,W., F.W.F., R.M., M.A.S., W.S.L,, H.S., E.A.E. did the experiments. E.A.E. designed the project, analyzed the data, wrote the paper and made the figures and tables. Coauthors provided input on manuscript drafts. We thank H.D. and colleagues for careful review of this paper.

## Supporting Information

### Supplemental Methods

#### Whole cell extract and pull-down methods in Fig 2

The large-scale co-immunoprecipitations of Ste5-Myc9 for mass spectrometry were done in 125 mM potassium acetate and no NaCl, whereas the co-immunoprecipitations of Ste5-Myc9 and HA3-Ste5 with Ssa1 and signaling partners were done with 250 mM NaCl and no potassium acetate. In Fig 1B, cells expressing Ste5-Myc9 were grown in SC-selective medium containing 2% dextrose to logarithmic phase to an A600 of 1.0 with or without treatment with 50 nM α factor for 2 hrs. Cells were pelleted, washed and immediately broken using liquid nitrogen and grinding with a mortar and pestle to break the cells, keeping samples in an ice bath in a cold room. Protein concentration was determined by the Biorad method and a portion of the samples were run on SDS-PAGE gels and the other portion was frozen in dry ice-ethanol and stored at −80°C. The extraction and immunoprecipitation (IP) buffers contained 20 mM Tris-HCl pH 7.2, 125 mM potassium acetate, 0.5 mM EGTA, 0.5 mM EDTA, 1 mM DTT, 0.1% Tween 20, 12.5% Glycerol (vol/vol) 1 mM 4-(2-aminoethyl)-benzenesulfonylfluoride, 1 mM sodium ortho-vanadate, 25 mM oo-glycerophosphate, 10 t g/ml pepstatin A.

In Fig 2, cells expressing Ssa1-GFP and either Ste5-Myc9 or HA3-Ste5 were grown in SC-selective medium containing 2% dextrose to mid-logarithmic phase of A600 of 0.8-1.0 and induced with α factor for 15 minutes (where indicated, 50 nM α factor for *bar1D/sst1D* cells, 5 μM for *BAR1/SST1* cells), pelleted, washed, frozen in an dry ice/ethanol bath, then thawed on ice and broken with six 30-s pulses with glass beads using a vortex after resuspending cells in modified-H buffer containing 25 mM Tris-HCl pH 7.4; 10% glycerol (vol/vol); 250 mM NaCl; 15 mM EGTA; 0.1% Triton-X-100; 1 mM sodium azide; 0.25 mM each meta- and orthovanadate; 1 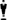 g/ml PMSF; 2 mM benzamidine; 4 mM 1,10-phenanthroline; 50 mM NaF; 5 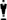 g/ml each of pepstatin A, leupeptin, peptin and chymostatin. Lysates were clarified with a 10 min centrifugation at 3,000 RPM to remove cellular debris followed by centrifugation for 10 min at 10,000 RPM at 4°C. Protein concentration was determined by the BioRad method and samples were frozen in dry ice ethanol bath and stored at −80°C. IPs were done with similar buffer with 200 mM NaCl and 7% glycerol.

## Supplemental tables

**Table S1 Plasmids and strains used in this study.**

**TableS2 Summary of ImageJ densitometry values.**

**Table S3 Localization of Ste5 derivatives to cell periphery and nucleus in wild type and *ssa1D ssa2D* cells.**

**Table S4 Summary of growth and mating of *ssa1*, *ssa2*, *ssa3*, *ssa4* single, double and triple mutants.**

**Table S5. List of positive and negative regulators of mating and invasive growth proteins screened for interaction with Ssa1, Ssa2 and other HSP70 network proteins in published databases.**

**Table S6. Summary of interaction data on 227mating pathway and invasive growth pathway proteins with Hsp70chaperonses Ssa1, Ssa2, Ssa3, Ssa4 and other Hsp70 networkproteins.**

**Table S7. Morphology of wild type and fes1 mutant strains before and after exposure to α factor.**

**Table S8 Cell morphology of Hsp70 family mutants before and after α factor treatment.**

## Supplemental figure titles and legends

**Fig S1. Analysis of full-length Ste5 (1-917) coding sequence with bioinformatics algorithms.** A. Cartoon of Ste5 with structural domains, domains of interaction with proteins and lipids and dimerization domains. B. PLAAC plot of Ste5. PLAAC (S11, S12) displays the propensity of a protein to adopt a prion or prion-like conformation along with intrinsic unfoldedness shown as the fold index (S13). The regions predicted to be most intrinsically unfolded are underlined in the amino acid sequence below the plot. C. RCS PDB plot (S14) of Ste5. D. Disopred3 analysis of Ste5 (S15). Di shows the amino acid sequence with residues predicted to be disordered enclosed in green boxes. Dii shows a confidence score for disorder (blue line) with regions predicted to interact with protein binding domains (orange line). E. IUPred plot (S16, S17) of disorder tendency for Ste5.

**Fig S2. Analysis of Ste5ms crystal structures 3FZE and 4F2H overlapping the VWA domain (residues 583-786).** A-E iCn3D renderings of Ste5 crystal structures. A-C. 3FZE ribbon, tube and amino acid side chain views. The 3FZE Ste5ms (residues 593-786) structure is at 1.16 angstrom resolution (S18). The polypeptide backbone is in magenta with the ste5L610/614/634/637A leucine to alanine mutation residues and part of an N-terminal helix are in yellow. D-E. Ribbon and tube views of 4F2H. The 4F2H Ste5(583-786) structure is at 3.18 angstrom (S19). The views show an anti-parallel homodimer with monomers in magenta and blue with the ste5L610/614/634/637A mutation residues and the entire N-terminal helix that dimerizes (residues 583-592) in yellow. F-I. Analysis of amino acid interactions of wild type and mutant Ste5 minimal scaffold dimer 4FH2 amino acid residues 589-798 with iCn3D 3.1.2 at NCBI (S20). Shown are predicted amino acid contacts with the wild type leucine residues and the mutant alanine residues. K. NCBI-BLASTP of *S. cerevisiae* Ste5 minimal scaffold. Positions 610, 614, 634 and 637 in Ste5 are highlighted in grey for identical and pink for similar. I. BIPPRED comparison (S22) of Ste5 wild type versus L610/614/634/637A mutant. Thirteen of the 107 predicted high maximal score (i.e. 0.9-0.99, where 1.0 is a perfect score) for Hsp70 binding in Ste5 have a lower score in the Ste5_L610/614/634/637A mutant than for the Ste5 wild type sequence. They are shown as bars overlapping the mutations. The difference in score between mutant and wild type is shown for 4 potential binding sites.

**Fig S3. Long exposure of Ste5-Myc9 and Ssa1-GFP immunoblot and tally of image showing punctate foci in cells expressing ste5L610/614/634/637A.** A. Long exposures of Fig. 2 immunoblot of Ste5-Myc9 from whole cell lysates. Lanes 1,2 are Ste5-Myc9 with HA3-GFP-Cdc24 and lanes 3, 4 are Ste5-Myc9 with Ssa1-GFP. Ste5-Myc9 was expressed from its own promoter on a 2 micron plasmid (pSKM90). Ssa1-GFP was expressed from the *ADH1* promoter on a *CEN* plasmid (pYMW122) and GFP-Cdc24 was expressed from the *MET25* promoter on a *CEN* plasmid (EBL664 415MET(GFPS65T)-A8-CDC24). 9E10 and DAPI colocalization in RGB color. C. 9E10 localization of Ste5-Myc9 in greyscale. In addition to the many punctate foci and patches visible in the photo, some cells exhibited what appears to be a network of globule like circles of Ste5-Myc9 which is indicated by two cartoon in line with two cells. D. Key of arrows for A. and B. E. Tallies of inclusion body characteristics.

**Fig S4. Comparison of hydrophobicity, bulkiness, average buried and disorder of amino acid residues of wild type Ste5 and mutant Ste5 L610/614/634/637A.** A. Alanine versus leucine characteristics. B-C. Bulkiness plots of Ste5(551-700) and Ste5mut(551-700) using algorithm based on Rose et al. (S23). D-E. Average buried plots on Ste5(551-700) and Ste5mut(551-700) using algorithm based on Zimmerman et al (S24, S25). The wild type 601-660 sequence is 601SLKREKPDN**<u>L</u>** AII**<u>L</u>**QIDFTK LKEEDSLIVV YNS**<u>L</u>**KA**<u>L</u>**TIK FARLQFCFVD RNNYVLDYGS 660 and the mutant sequence is 601SLKREKPDN**<u>A</u>** AII**<u>A</u>**QIDFTK LKEEDSLIVV YNS**<u>A</u>**KA**<u>A</u>**TIK FARLQFCFVD RNNYVLDYGS 660. F. IUPRED plot (S26-S28) of Ste5(551-700). G. IUPRED plot of Ste5L610/614/634/637A (551-700). H. IUPRED plot of known disordered protein that aggregates alpha synuclein. I. IUPRED2 and ANCHOR plots (S28) of Ste5(1-551-700). J. IUPRED2 and ANCHOR plot of Ste5L610/614/634/637A (551-700). Amino acid sequences were analyzed at the IUPRED server and the ISOPRED2 server //iupred2a.elite.hu/ (S26-S28). Other analyses were done at the EXPasy website with default 9 amino acid scanning window (S29).

**Fig S5. Summary of predicted chaperone interactions with the mating and invasive growth pathways reveals bias towards Ssa1.** A comparison of predicted interactions of Ssa1, Ssa2, Ssa3, Ssa4, Sse1 and Sse2 for mating pathway and invasive growth pathway proteins. A-B Distribution among 227 mating and invasive growth pathway proteins. C-D. Distribution among core signaling components of mating and invasive growth pathways. A literature search was done to compile potential interactions between Hsp70 chaperones and the mating and invasive growth pathways from published data bases. See Table S5, Table S6 and databases of interactions (S30-S39). The majority of the reported interactions come from unconfirmed high-throughput experiments.

**Fig S6. Representative fields of wild type and Hsp70 mutants treated with 2.5 μM α factor.** Cells were grown at room temperature to logarithmic phase and treated with 2.5 μM αF for 90 minutes. A. WT EYL1740 (S288c-BY4741), B. *fes1*, C. *ydj1*. See Table S7 and Table S8 for tallies of morphology.

**Fig S7. Representative fields of wild type and hsp70 mutants treated with 5 μM αF.** Cells were grown at room temperature to logarithmic phase and treated with 5 μM αF for 90 minutes. A. WT EYL1740 (S288c-BY4741), B. *ydj1*, C. *ssb1*, D. *sse1*, E. *sse2*, F. *ssz1*, G. *sti1*. See Table S8 for tallies of morphology.

