## Supplementary material for "Effects of HSP70 chaperones Ssa1 and Ssa2 on Ste5 scaffold and the mating mitogen-activated protein kinase (MAPK) Pathway in *Saccharomyces cerevisiae*": FigS1

A.

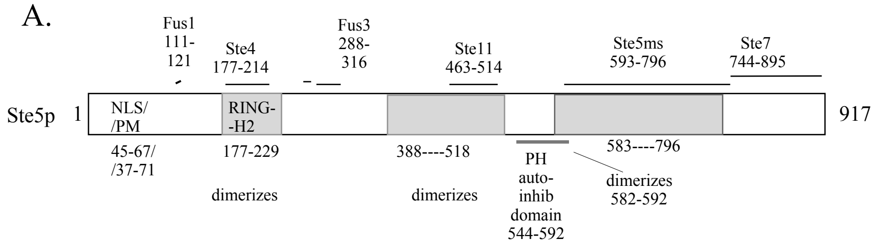


B. PLAAC Ste5(1-917) full length

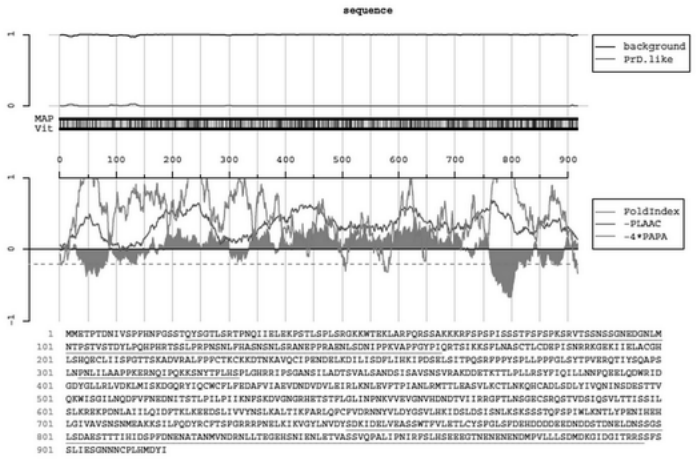


C. RSC PDB Ste5(1-917) full length

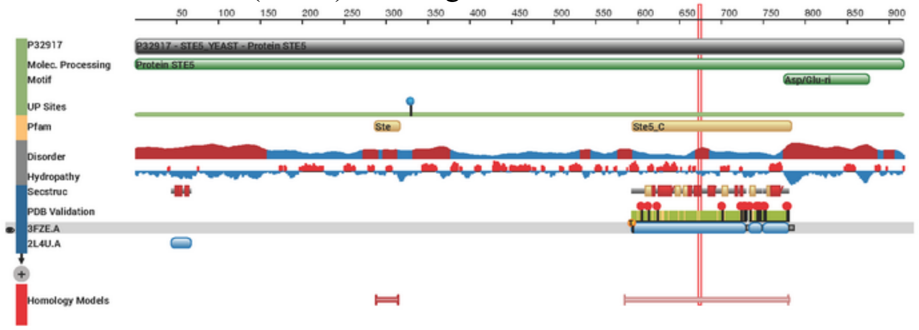


Di.

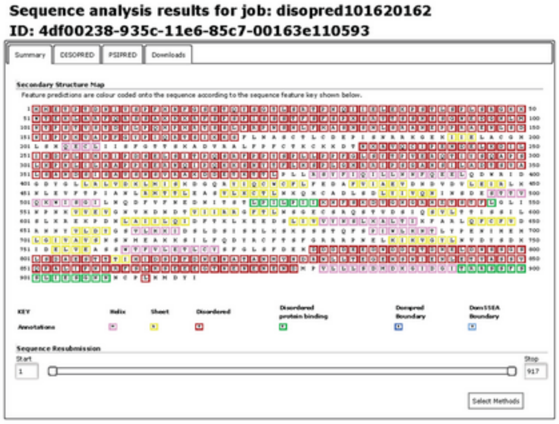


Dii.

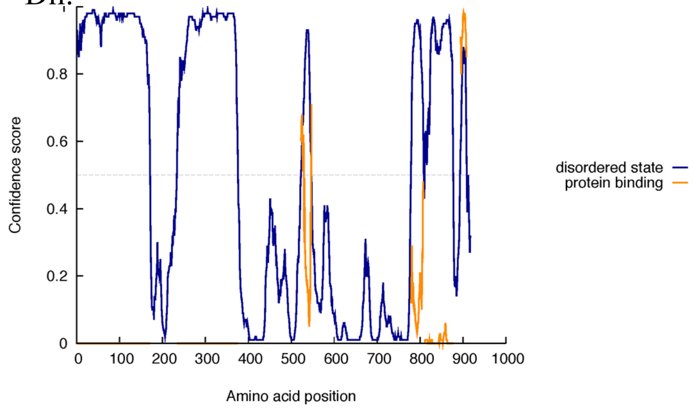


Intrinsic disorder profile

E. IUPRED
Ste5(1-917) full length

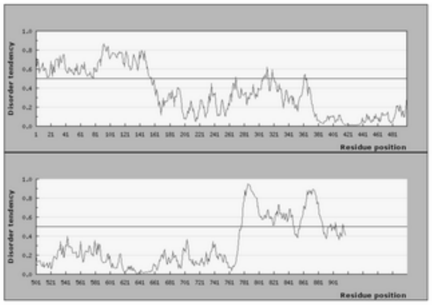
