## Supplementary material for "Effects of HSP70 chaperones Ssa1 and Ssa2 on Ste5 scaffold and the mating mitogen-activated protein kinase (MAPK) Pathway in *Saccharomyces cerevisiae*": FigS2

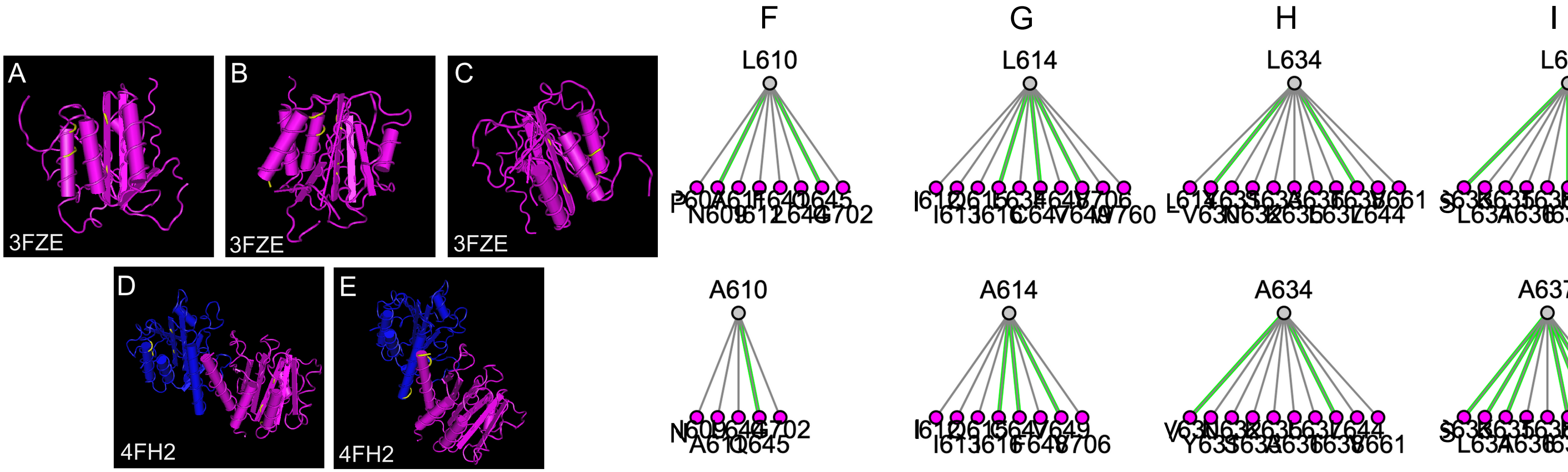


J KEY: iCn3D 3.1.2 Interactions

**Green**: H-Bonds; **Cyan**: Salt Bridge/Ionic; **Grey**: contacts
**Magenta**: Halogen Bonds; **Red**: π-Cation; **Blue**: π-Stacking

K BLASTP

```
                                       610  614                       634 637
Ste5 593:    1  LTTISSILSLKREKPDNLAIILQIDFTKLKEE--DSLIVVYNSLKALTIKFARLQFCFVD 650
S. cer 1:  593  LTTISSILSLKREKPDNLAIILQIDFTKLKEE--DSLIVVYNSLKALTIKFARLQFCFVD 650
S. cer 2:  593  LTTISSILSLKREKPDNLAIILQIDFTKLKEE--DSLIVIYNSLKALTIKFARLQFCFVD 650
S. cer 3:  593  LTTISSILSLKREKPDNLAIILQXDFTKLKEE--DSLIVVYNSLKALTIKFXRLQFCFVD 650
k. lac     498  LTSISSILSIKRATPNELNLVLQLDKRKTNE---DNIVSIKNVILALTWKFSTFNCCIVD 554
k. afr     585  MTTISSILSLKRERPDDLVIIFQIDFFKVKN--NDDILNFYNILKALLLKFPECKFCAVN 642
N. dai     641  LTTISSILSLKRERPDDIIIVLQLDLNKLKL--GNNSTTLYNSLTALNMKFPNFKICIVD 698
N. cas     614  MTTISSILSLKRERPDDVVIVLQIDFKKINNE--DDYTTITNTLKALTFKFPRLKACIVD 671
V. pol     598  MTSVSSILSLKRERPEGIVIILQLDFRKLKD---DSYIIIYNSLQALVTKYPNLIYCTVD 654
Z. rou     565  MTNISSILSLKRERPDELVVVLQLDFEKIRT--KNNSLVIFNSLKALVLKFPDMKICIVD 622
a. gos     513  RASINSILSLKRRHPDELCLFFQIDTAKVQA---VDYKILANTCKAILWKFNTAKVALAD 569
c. gla     370  NTITSSVLSIKKEKPSDILLLIQIDEQKITDQKDQIKQVIKNTLMALKLKYPEYFLVILS 429
```

610 614 634 637

STE5 593 LTTISSILSLKREKPDN^L^AII^L^QIDFTKLEEDSLIVVYNS^L^KA^L^TIKFARLQFCFD 650

L BIPPRED MUTANT VS WT MAX SCORE

KPDNLAI -7.2214
ILQIDFT -0.1197
IVVYNSL -0.1299
NSLKALT -0.186
