## Supplementary material for "Effects of HSP70 chaperones Ssa1 and Ssa2 on Ste5 scaffold and the mating mitogen-activated protein kinase (MAPK) Pathway in *Saccharomyces cerevisiae*": FigS3

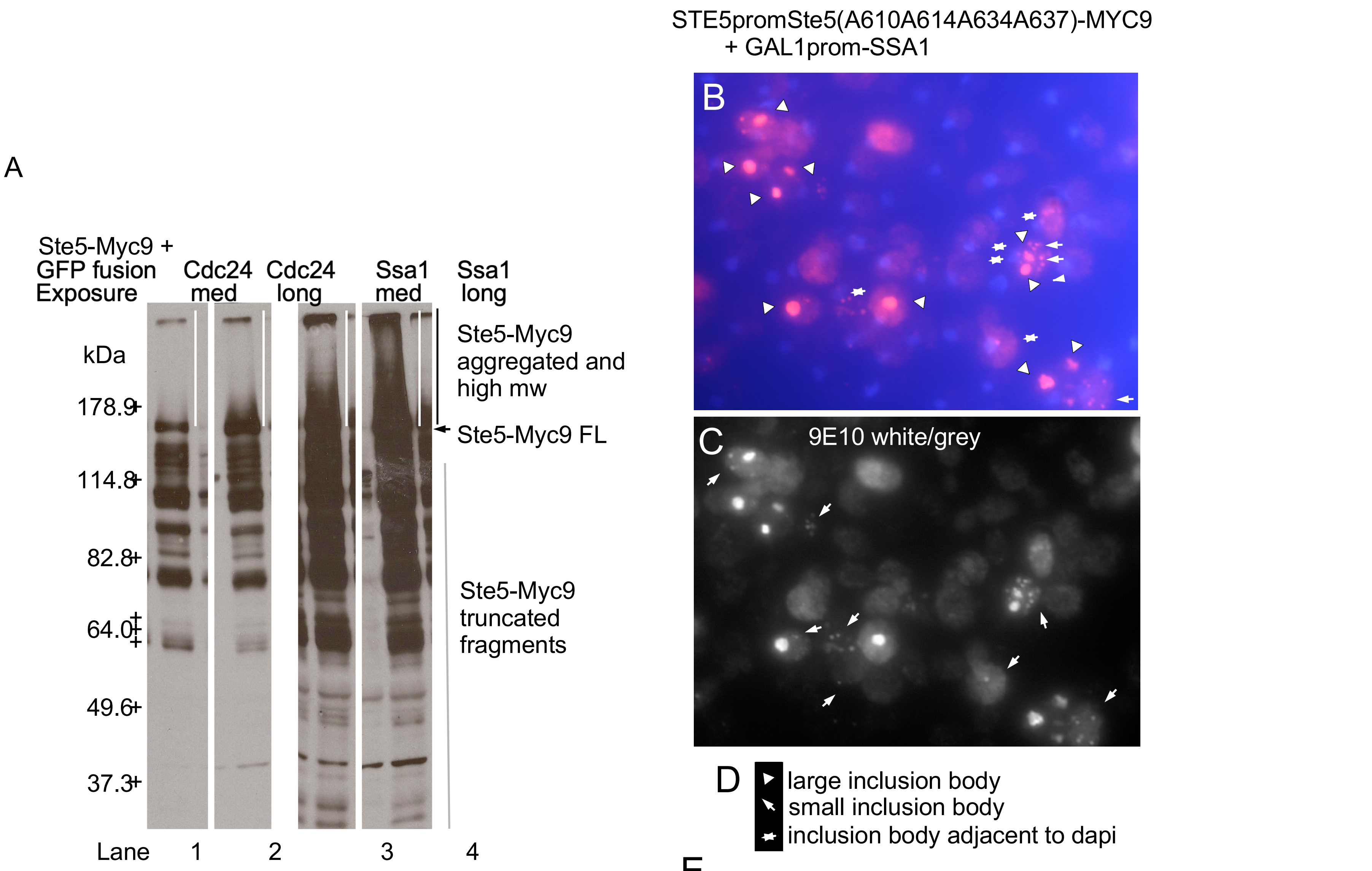


E

59: #Dapi staining nuclei
41: #9E10 staining cells
17: #9E10 staining cells with inclusion bodies
10: #9E10 cells with large inclusion bodies
12: #9E10 cells with small inclusion bodies
0: #9E10 cells with inclusion bodies overlapping dapi
2: #9E10 cells with small inclusion bodies next to dapi
