## Supplementary material for "Effects of HSP70 chaperones Ssa1 and Ssa2 on Ste5 scaffold and the mating mitogen-activated protein kinase (MAPK) Pathway in *Saccharomyces cerevisiae*": FigS4

A Alanine vs Leucine

Bulkiness:
Ala: 11.500 Leu: 21.400

Tendency to be buried:
Ala: -0.29 Leu: -0.02

Average Flexibility:
Ala: 0.360 Leu: 0.370

% Accessible residues:
Ala: 6.600 Leu: 4.800

B Bulkiness
Ste5(551-600)

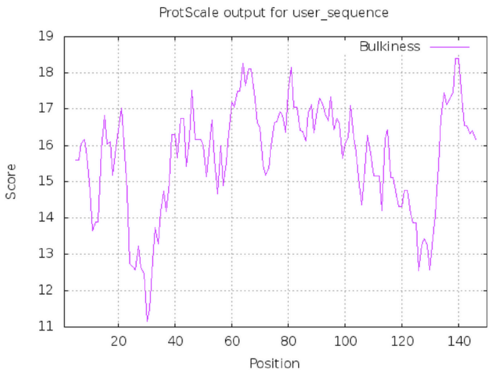


C Bulkiness
Ste5Leu610,614,634,637toAla(550-600)

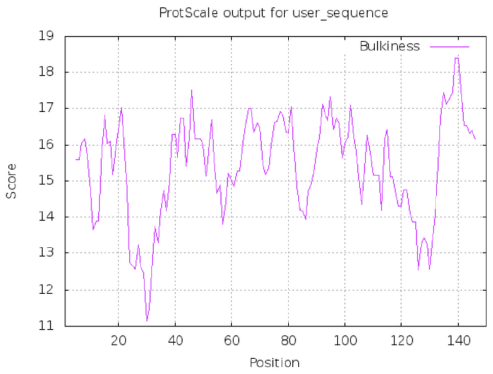


D Average Buried
Ste5(551700)

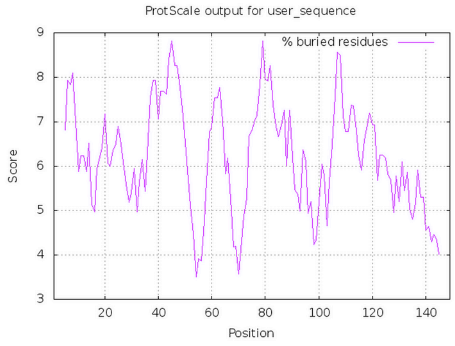


E Average Buried
Ste5Leu610,614,634,637toAla(551-700)

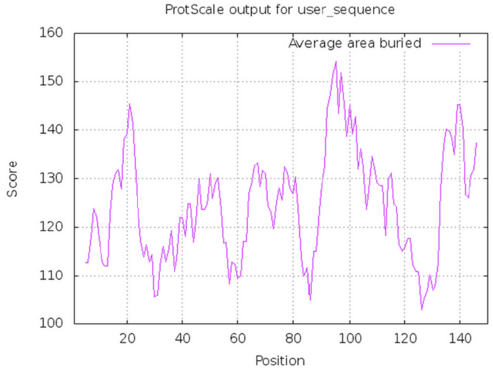


F. IUPRED STE5 (551-700)

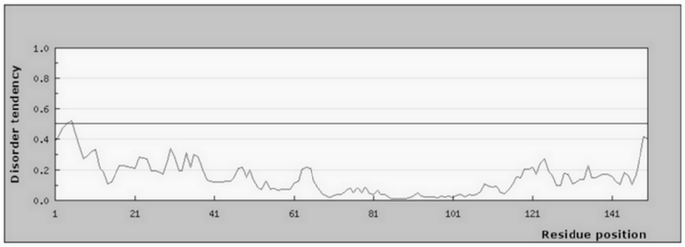


G. IUPRED
STE5 L610/614/634/637A (551-700)

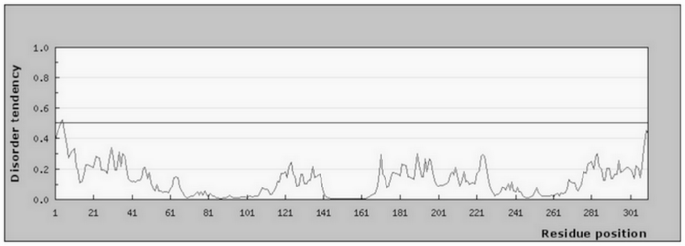


H. IUPRED control
alpha synuclein (human P37480)

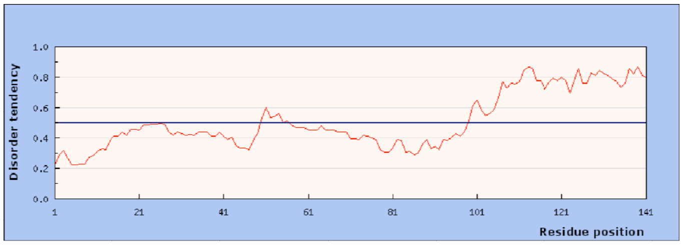


I. IUPRED2, ANCHOR
STE5 (551-700)

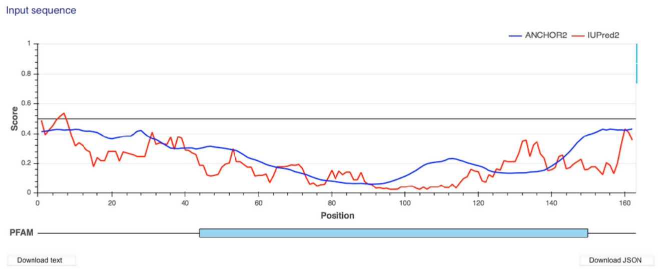


J. IUPRED2, ANCHOR
STE5 L610/614/634/637A (551-700)

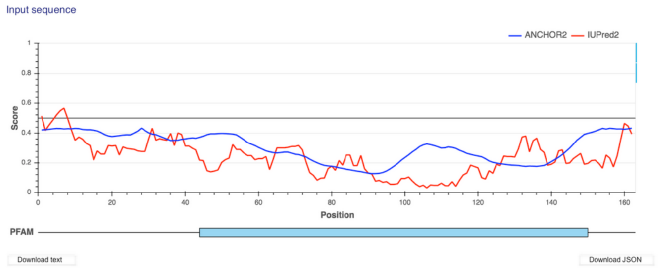
