## Supplementary figures and images for "Effects of HSP70 chaperones Ssa1 and Ssa2 on Ste5 scaffold and the mating mitogen-activated protein kinase (MAPK) Pathway in *Saccharomyces cerevisiae*"

### FigS5

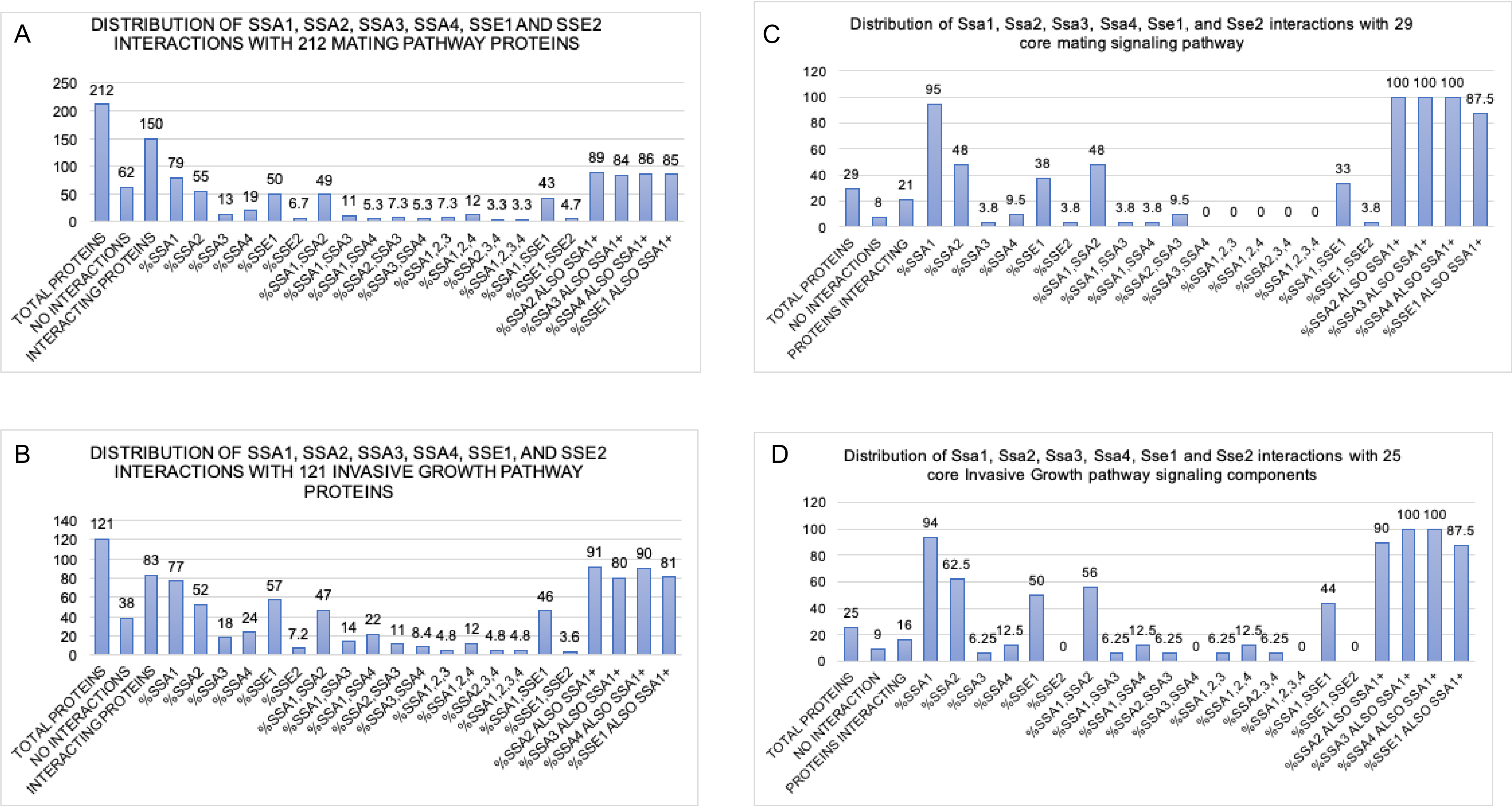

### FigS6

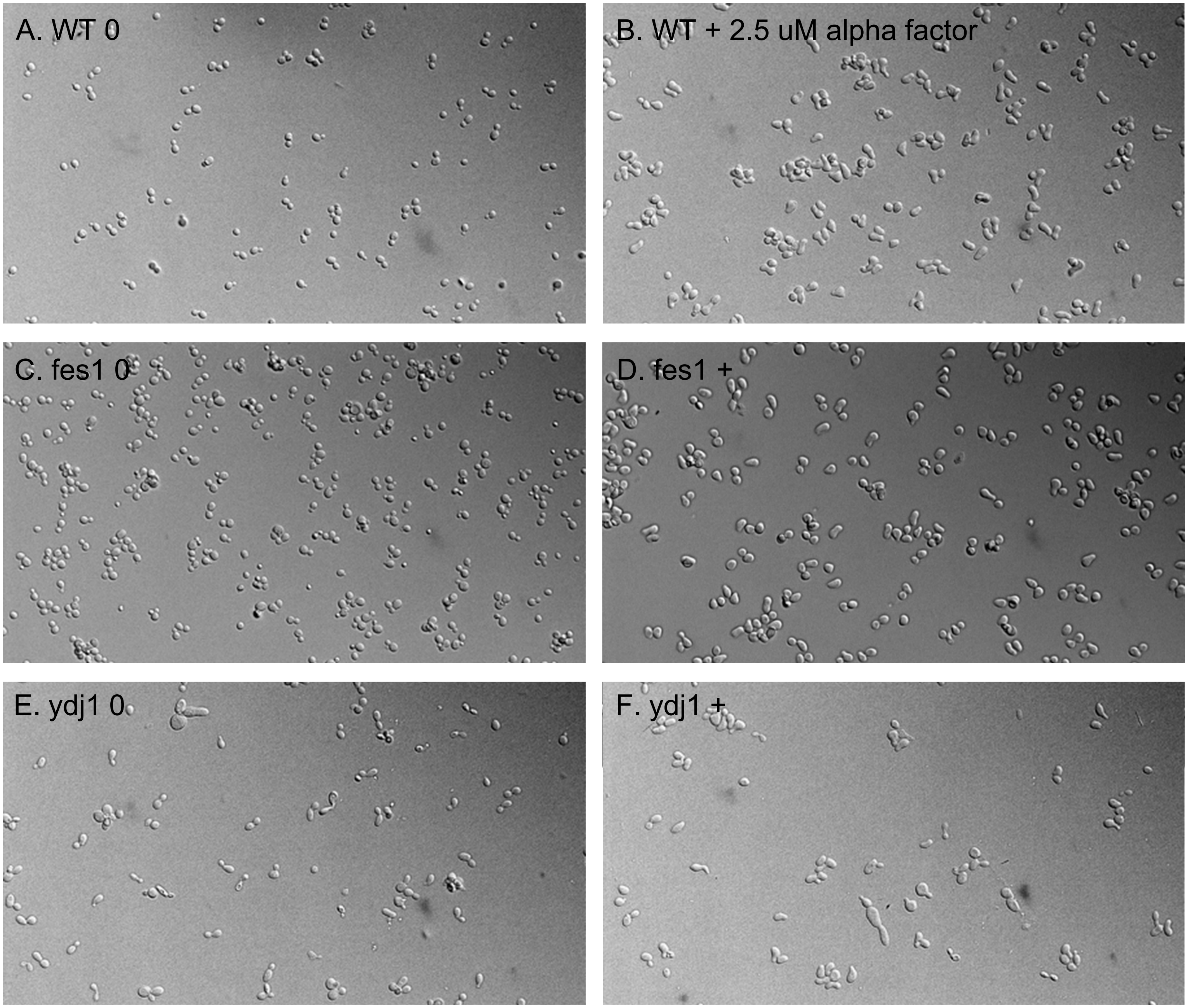

### FigS7

A. WT +5mM aF

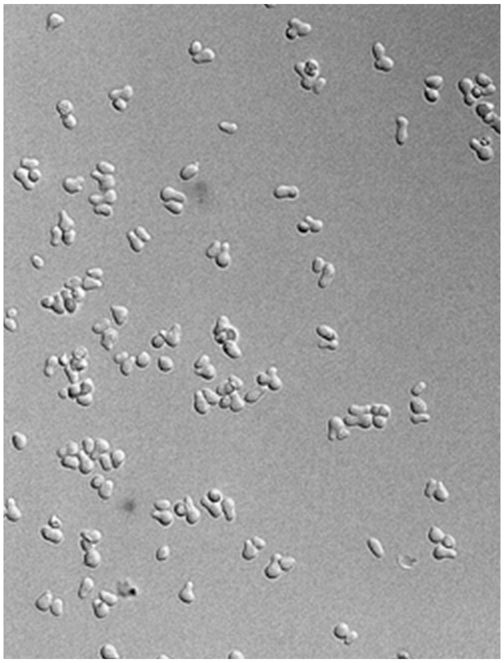

B. ydj1 +5mM aF

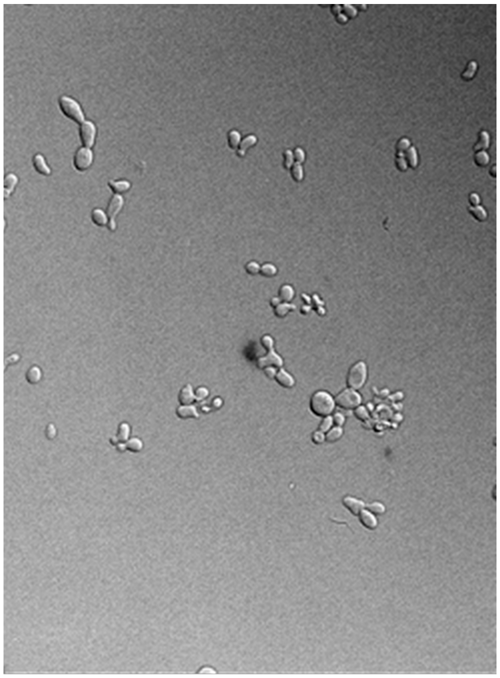

C. ssb1 +5mM aF

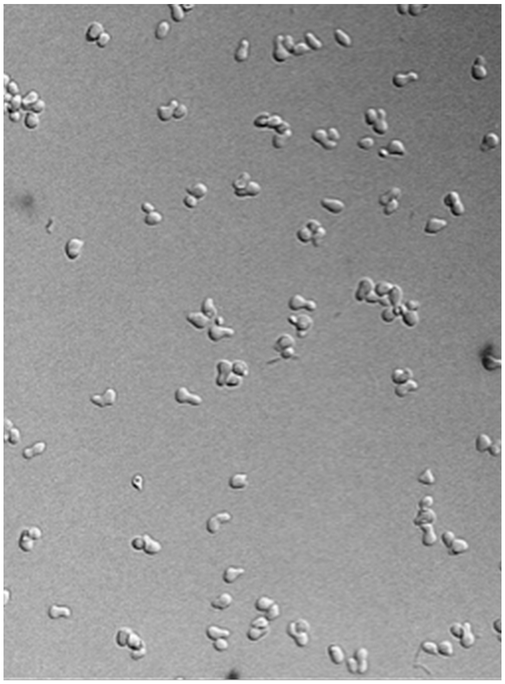

D. sse1 +5mM aF

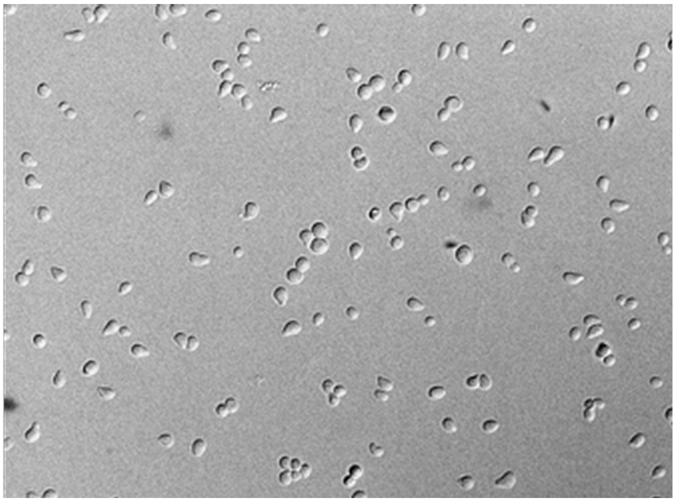

E. sse2 +5mM aF

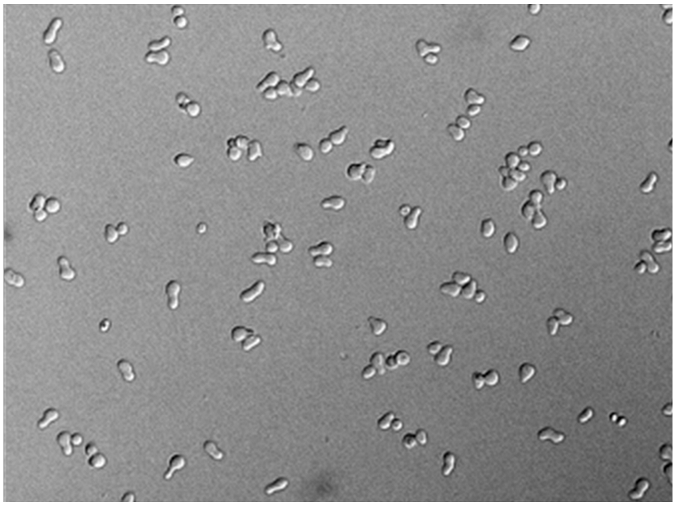

F. ssz1 +5mM aF

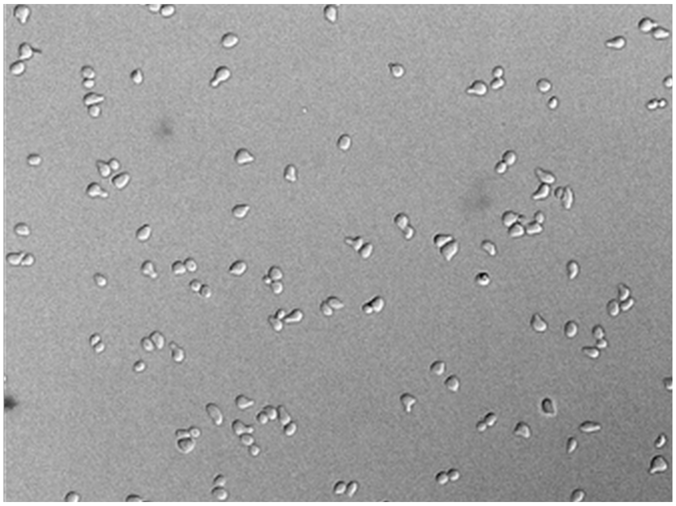

G. sti1 +5mM aF

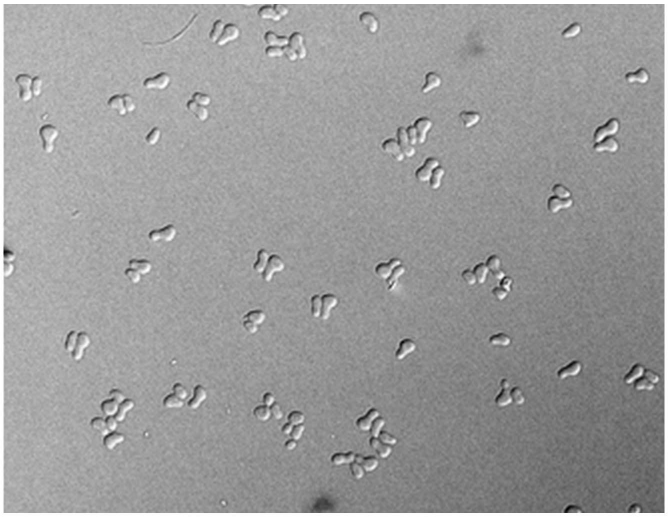
