## Supplementary material for "Effects of HSP70 chaperones Ssa1 and Ssa2 on Ste5 scaffold and the mating mitogen-activated protein kinase (MAPK) Pathway in *Saccharomyces cerevisiae*": TableS1

Table S1. Plasmids and strains used in this study.

| Plasmid or Strain | Description | | | Source1 |
|---|---|---|---|---|
| PLASMIDS | | | | |
| EBL95 | pDL1460/pGA1815 *FUS1::ubiYlacZ* | *URA3* | *CEN* | (S1) B. Errede |
| pRM1 | *STE5-Myc9* | *TRP1* | *2μ* | R.M., this study |
| pYMW122, 123 | *ADH1p-SSA1-GFP* | *URA3* | *CEN* | Y.W., this study |
| EBL530 | *GAL1prom-SSA1* | *LEU2* | *CEN* | Y.W., this study |
| EBL531, 536 | *GAL1prom-SSA1* | *TRP1* | *CEN* | Y.W. this study |
| EBL532, 535 | *GAL1prom-SSA1* | *URA3* | *CEN* | Y.W. this study |
| pYMW5 | *ste5Δ522-527-Myc9* | *URA3* | *2μ* | (S2) EE lab |
| pYMW15 | *ste5L634A/637A-Myc9* | *URA3* | *2μ* | (S2) EE lab |
| pYMW37 | *ste5L482A/485A*-Myc9* * L483/486A with both Met1Met2* | *URA3* | *2μ* | (S3) EE lab |
| pYMW38 | *ste5L610/614A-Myc9* | *URA3* | *2μ* | (S2) EE lab |
| pYMW65 | *ste5D522-527-Myc9* | *URA3* | *2μ* | (S4) EE lab |
| pYMW107.8 | *TAgNLS-ste5L610A/614A/634A/637A-Myc9* | *URA3* | *2μ* | Y.W. EE lab |
| pYMW110.29 | *ste5L610A/614A/634A/637A-Myc9* | *URA3* | *2μ* | Y.W. EE lab |
| pYMW120 | *GAL1prom-TAgNLS- ste5L610A/614A/634A/637A-Myc9* | *URA3* | *CEN* | Y.W. EE lab |
| pSKM12 | *STE5-Myc9* | *URA3* | *CEN* | (S4) EE lab |
| pSKM17 | *CUP1prom-STE5-Myc9* | *URA3* | *CEN* | (S4) EE lab |
| pSKM19 | *STE5-MYC9* | *URA3* | *2μ* | (S4) EE lab |
| pSKM21 | *CUP1prom-GFP-STE5* | *URA3* | *CEN* | (S4) EE lab |
| pSKM30 | *GAL1p-STE5-MYC9* | *URA3* | *CEN* | (S4) EE lab |
| pSKM72 | *ste5-K49A50A-Myc9* | *URA3* | *2μ* | (S4) EE lab |
| pSKM74 | *ste5-K64-66A-Myc9* | *URA3* | *2μ* | (S4) EE lab |
| pSKM76 | *ste5-K49A50A/64-66A-Myc9* | *URA3* | *2μ* | (S4) EE lab |
| pSKM87 | *HA3-STE5* | *URA3* | *2μ* | (S4) EE lab |
| pSKM90 | *STE5-Myc9* | *HIS3* | *2μ* | (S4) EE lab |
| pSKM98 | *TAgNLS*$^{K128T}$*-STE5-Myc9* | *URA3* | *2μ* | (S4) EE lab |

| | | | | |
|---|---|---|---|---|
| pSKM102 | *pSKM102[pGal TAgNLS-Ste5 M9 CEN]* | *URA3* | *CEN* | (S4) EE lab |
| pSKM71 | *EBL418 ste5K49,50A-Myc9* | *URA3* | *2μ* | (S4) EE lab |
| pSKM73 | EBL420 *ste5K64-66A-Myc9* | *URA3* | *2μ* | (S4) EE lab |
| pSKM76 | EBL423 *ste5K49,50,D64-66A-Myc9* | *URA3* | *2μ* | (S4) EE lab |
| pSKM113 | *STE5(1-242)-GFP2* | *URA3* | *CEN* | (S2) EE lab |
| pSKM115 | *STE5(1-242Δ49-66)-GFP2* | *URA3* | *CEN* | (S2) EE lab |
| pNC318 | *CYC1promSTE7-MYC* | *TRP1* | *CEN* | (S5) B. Errede |
| EBL664 | *415MET25(GFPS65T)-A8-CDC24 (MET25 promoter)* | *LEU2* | *CEN* | (S6) D. Johnson |
| YCp403 | *HIS3 CEN4* | *HIS3* | *CEN* | Y.W. EE lab |
| YCplac22 | *TRP1 CEN4* | *TRP1* | *CEN* | Y.W. EE lab |
| pYBS305 | *GAL1prom-GST URA3* | *LEU2* | *2μ* | (S7) EE lab |
| ZM43 | *URA3 CEN* | *URA3* | *CEN* | Z.Moqtaderi, K. Struhl |
| B1818 | *TRP1 CEN* | *TRP1* | *CEN* | EE lab |

STRAINS E. coli:

| | | |
|---|---|---|
| DH5α | F – endA1 glnV44 thi-1 recA1 relA1 gyrA96 deoR nupG purB20φ80dlacZΔM15 Δ(lacZYA-argF)U169 hsdR17(rK –mK+), λ lacIq lacY+ | EE lab |
| HB101 | hsdR17(rK –mK+), λ lacI$^q$ lacY$^+$ | EE lab |
| XL10 Gold | endA1 glnV44 recA1 thi-1 gyrA96 relA1 lac Hte Δ(mcrA)183 Δ(mcrCB-hsdSMR-mrr)173 tet R F'[proAB lacI qZΔM15 Tn10(Tet R Amy Cm R)] | EE lab |

STRAINS *S. cerevisiae*
W303a background

| | | |
|---|---|---|
| EY699 | W303a *MATa ura3-1 leu2-3,112 trp1-1 his3-11,15 ade2-1 can1-100 Gal+* | R. Rothstein |
| EY957 | *MATa bar1D::hisg ura3-1 leu2-3,112 trp1-1 ade2-101 his3-11, 15 can1-100* | EE lab |
| EY1457 | EY1411 (*MATa bar1D*::hisg *ura3-1 leu2-3,112 trp1-1 ade2-101 his3-11, 15 can1-100*+pYBS305) + YCp403 | (S8) EE lab |

| | | |
|---|---|---|
| EY1775 | *MATa bar1D::hisg ste5D:TRP1 ura3-1 leu2-3,112 trp1-1 ade2-101 his3-11, 15 can1-100* | EE lab |
| EY3387 | BY58 *ste5D::TRP1. + YEplac195 URA3 2µ* | P.M. , this study |
| EY3390 | EY940 + *YEplac195 URA3 2µ* | P.M., this study |
| EY3392 | EY957 + EBL432 *pCUP1prom-STE5-Myc9 CEN* | P.M., this study |
| EY3393 | EY957 + EBL444 *TAgNLSK128T-STE5-Myc9* | P.M., this study |
| EY3397(1-6) | *EYK365 ste5D::ADE2 + EBL536 pGAL1prom-SSA1-TRP1-CEN* | P.M., this study |
| EY3398 (1-3) | *EYL365 ste5D::ADE2 + EBL418 ste5K49,50A-Myc9 +* EBL536 *pGAL1prom-SSA1-TRP1-CEN* | P.M., this study |
| EY3399 (1-3) | EYL365 *ste5D::ADE2 + EBL536 pGAL1prom-SSA1-TRP1 +* EBL418 *ste5K64A,K66A-Myc9-URA3-2µ* | P.M., this study |
| EY3400 (1-3) | EYL365 *ste5D::ADE2 + EBL536 pGAL1prom-SSA1-TRP1 +* EBL418 *ste5K49A,K50A, K64-66A-Myc9-URA3-2µ* | P.M., this study |
| EY3401 (1-3) | EYL365 *ste5D::ADE2 +EBL536 pGAL1prom-SSA1-TRP1 +* EBL418 *STE5(1-242)-GFP-GFP-URA3-CEN* | P.M., this study |
| EY3402 (1-3) | EYL365 *ste5D::ADE2 + EBL536 pGAL1prom-SSA1-TRP1 + pADHprom-STE5(1-242D49-66)-GFP-GFP URA3* | P.M., this study |
| EY3408 | EYL365 *ste5D::ADE2 + EBL365 pSKM19 STE5-Myc9-URA3-CEN* | P.M., this study |
| EY3411 | EYL365 *ste5D::ADE2 + B1818 TRP1-CEN* | P.M., this study |
| EYL357 | EY699 *msn5D::HIS3* | (S2) EE lab |
| EYL457, RAY914 | *leu2-3,112 trp1-D901 his3-D200 lys2-801 ura3-52 gal2 suc2-D9 (pRS414 CDC24) cdc24D-1::LoxP HIS5 SpLoxP* | R. Arkowitz |
| EYL2619 | YMY374: EY1775 *ste5D::TRP1 +* pSKM19 *STE5-Myc9-URA3-2µ* | Y.W., this study |
| EYL2620 | YMY375: EY1775 *ste5D::TRP1 +* pSKM107 *STE5-Myc9 -LEU2-2µ* | Y.W. this study |
| EYL2765 | YMY520: EY1775 *ste5D::TRP1+* pYMW106.2 *TAgNLSK128T-ste5L610A/614A/634A/637A-Myc9 URA3-2µ* | Y.W. this study |
| EYL2766 | YMY521: EY1775 *ste5D::TRP1 +* | Y.W., this study |

| | | |
|---|---|---|
| | pYMW107.8 *TAgNLS-ste5L610A/614A/634A/637A-Myc9-URA3-2μ* | |
| EYL2769 | YMY524: EY1775 *ste5D::TRP1* + pYMW110.29 *ste5L610A/614A/634A/637A-Myc9-URA3-2μ* | Y.W., this study |
| EYL2771 | YMY526: EYL365 (*MATa ste5D::ADE2*) + EBL531 *GAL1prom-SSA1-TRP1-CEN* | Y.W., this study |
| EYL2778 | YMY533: YMY526 (EYL365 *ste5D::ADE2* EBL531 *GAL1prom-SSA1-URA3-CEN)* + pYMW37 *ste5L482/485A-Myc9-URA3-2μ* | Y.W., this study |
| EYL2795 | YMY550: YMY526 (EYL365 *ste5D::ADE*2) + pYMW110.29 *ste5L610A/614A/634A/637A-Myc9-URA3-2μ* | Y.W., this study |
| EYL2812 | YMY567: EY957 + pYMW118 *pSTE5prom-TAgNLS-ste5L610A/614A/634A/637A-LEU2-CEN* | Y.W., this study |
| EYL2816 | YMY571: EYL957 *ste5D::ADE2* + pYMW120 *pGAL1prom-TAgNLS-ste5L610A/614A/634A/637A-Myc9 URA3-2μ* | Y.W., this study |
| EYL2823 | YMY578: YMY570 (EY957 + pYMW123 *pADHprom-SSA1-GFP-URA-2μ)* + pSKM90 *STE5-Myc9-HIS2-2μ* | Y.W., this study |
| EYL3022 | YMY784: EY699 + pSKM19 *STE5-Myc9-URA3-2μ* | Y.W., this study |
| EYL3023 | YMY785: EYL357 *msn5D::HIS3* + pSKM19 *STE5-Myc9-2μ-URA3* | Y.W., this study |
| EY3411(1-4) | EYL365 *MATa ste5D::ADE2 + B1818-TRP1-CEN* | P.M., this study |
| EY3423 | EYL365 *MATa ste5D::ADE2 + B1818-TRP1-CEN* + pSKM113 (EBL459) *STE5(1-242)-GFP2* | P.M., this study |
| EY1775/98 | EY1775 + pSKM98 *TAgNLS*$^{K128T}$*-STE5-Myc9-URA3-2μ* | Y.W., this study |
| YMW467 | EYL457 + pSKM19 *STE5-Myc9 URA3-2μ* | Y.W., this study |
| EYL2778/YMY533 | YMY526: EYL365 (*MATa ste5D::ADE2*) + EBL531 *GAL1prom-SSA1-TRP1-CEN* ) + *ste5L482A,L485A-Myc9-URA3-2μ* | Y.W., this study |
| EYL2779/YMY534 | YMY526: EYL365 (*MATa ste5D::ADE2*) + | Y.W., this study |

| | | |
|---|---|---|
| | EBL531 *GAL1prom-SSA1-TRP1-CEN* ) + pYMW65 *ste5D522-527-Myc9-URA3-2µ* | |
| EYL2780/YMY535 | YMY526: EYL365 (*MATa ste5D::ADE2*) + EBL531 *GAL1prom-SSA1-TRP1-CEN* ) + pYMW15 *ste5L634A/637A-Myc9-URA3-2µ* | Y.W., this study |
| EYL2781/YMY536 | YMY526: EYL365 (*MATa ste5D::ADE2*) + EBL531 *GAL1prom-SSA1-TRP1-CEN* ) + pYMW110.29 *ste5L610A/614A/634A/637A-Myc9-URA3-2µ* | Y.W., this study |
| STRAINS-FY23 background (isogenic to S288c) | | |
| FY23 | *MATa ura3-52 leu2D1 trp1D63 GAL2* | F. Winston |
| EYL275 | YMY505: FY23 + *STE5-GST CEN URA3* | (S2) EE lab |
| EYL308 | YMY851: FY23 + pSKM113 *Ste5(-1-242)-GFP-GFP* | Y.W., this study |
| FY23/98 | FY23 (S288c) + pSKM98 *TAgNLSK128T-Ste5-Myc9* | Y.W., this study |
| STRAINS-Craig lab | | |
| MW143 | *MATa leu2-3,112 Dtrp1 ura3-52 his3-11,15 lys2 GAL2 ssa1::HIS3 ssa2D::URA3* | (S9) E. Craig |
| EYL339 | MW142 vial Y234 M*ATα GAL2 his3-11,15 leu2-3,112 Dtrp1 lys2 ssa1D:HIS3 ssa2D::URA3* (not *MATa*) | (S9) E. Craig |
| EYL340 | MW143 *MATa ssa1::HIS3 ssa2::URA3 his3-11, 15 leu2-3,112 lys2 Dtrp1 ura3-52* | (S9) E. Craig |
| EYL341, EYL344 | *T211, DS10 vial Y166 MATa his3-11,15, leu2-3,112 lys1 lys2 Dtrp1 ura3-52* | E. Craig |
| EYL342 | MW331 *MATa his3-11,15 leu2-3,112 lys2 Dtrp1 ura3-52 ssa1::HIS3 ssa2::LEU2 ssa4::LYS2 + pGAL1prom-SSA1-URA3-CEN4* | E. Craig |
| EYL349/MW63 | EY3135 *MATa/Matα his3-11,15/his3-11,15 lys2/lys2 ura3-52/ura3-52 leu2-3, 112/leu2-3,112 Dtrp1/Dtrp1 GAL2/GAL2 ssa1::HIS3/+ ssa2::LEU2/+ +/ssa3::TRP1 +/ssa4::URA3* | E. Craig |

| STRAINS-Haploid strains derived from EYL349 | | |
|---|---|---|
| EY3136 | *MATa his3-11,15 leu2-3,112 lys2 Dtrp1 ura3-52 GAL2* | F. F., this study |
| EY3137 | *MATa ssa1::HIS3* | F. F., this study |
| EY3138 | *MATa ssa2::LEU2* | F. F., this study |
| EY3141 | *MATa ssa1::HIS3 ssa2::LEU2* | F. F., this study |
| EY3143 | *MATa ssa1::HIS3 ssa4::URA3* | F. F., this study |
| EY3144 | *MATa ssa2::LEU2 ssa3::TRP1* | F. F., this study |
| EY3145 | *MATa ssa2::LEU2 ssa4::URA3* | F. F., this study |
| EY3146 | *MATa ssa3::TRP1 ssa4::URA3* | F. F., this study |
| EY3147 | *MATa ssa1::HIS3, ssa2::LEU2 ssa3::TRP1* | F. F., this study |
| EY3148 | *MATa ssa1::HIS3 ssa3::TRP1 ssa4::URA3* | F. F., this study |
| EY3158 | EY3136 *WT+ EBL295 pDL1460/pGA1815 FUS1::ubiYlacZ* | F. F., this study |
| EY3159 | EY3141 *MATa ssa1::HIS3 ssa2::LEU2 +* EBL295 *pDL1460/pGA1815 FUS1::ubiYlacZ* | F. F., this study |
| EY3160 | EY3147 *MATa ssa1::HIS3, ssa2::LEU2 ssa3::TRP1 +* EBL295 *pDL1460/pGA1815 FUS1::ubiYlacZ* | F. F., this study |
| EY3149 | *MATa ssa2::LEU2 ssa3::TRP1. ssa4::URA3* | F. F., this study |
| EY3156 | EY3136 + EBL295 *pGA1815/pDL1460 FUS1::ubiYlacZ* | F. F., this study |
| EY3158 | EY3136 + EBL295 *pGA1815/pDL1460 FUS1::ubiYlacZ* | F. F., this study |
| EY3159 | EY3141 + EBL295 pGA1815/pDL1460 FUS1::ubiYlacZ | F. F., this study |
| EY3160 | EY3147 + EBL295 pGA1815/pDL1460 FUS1::ubiYlacZ | F. F., this study |
| EY3181 | EY3141 + EB238 FUS3-HA | P.M., this study |
| EY3311 | EY3136 + pSKM30 *pGAL1prom-STE5-Myc9-URA3-CEN* | F. F., this study |
| EY3312 | EY3136 + ZM43 *URA3-CEN* | F. F., this study |
| EY3313 | EY3141 + pSKM30 *pGAL1prom-STE5-Myc9-URA3-CEN* | F. F., this study |
| EY3314 | EY3141+ ZM43 *URA3-CEN* | F. F., this study |
| EY3315 | EY3148 5-FOA$^R$ *MATa ssa1::HIS3 ssa3::TRP1 ssa4::ura3-* Non-reverting ura3- 5FOA resistant derivative of EY3148 | F. F., this study |

| | | |
|---|---|---|
| EY3335 | *MATa ssa1::HIS3 ssa3::TRP1 ssa4::ura3-* + EBL295 pGA1815/pDL1460 FUS1::ubiYlacZ | F. F., this study |
| EY3350 | EY3136 + pNC245 *pCYC1prom-STE7-MYC-TRP1-CEN* | F. F., this study |
| EY3353 | EY3141 + pNC245 *pCYC1prom-STE7-MYC-TRP1-CEN* | F. F., this study |
| EY3376 | EY3136 + pSKM19 *STE5-Myc9* | P.M., this study |
| EY3377 | EY3136 + pSKM98 *TAgNLSK128T- STE5-Myc9* | P.M., this study |
| EY3378 | EY3136 + EB238 *FUS3-HA* | P.M., this study |
| EY3379 | EY3141 + pSKM19 *STE5-Myc9* | P.M., this study |
| EY3380 | EY3141 + pSKM98 *TAgNLSK128T-STE5-Myc9* | P.M., this study |
| EY3384 (1,2) | EY3376 (EY3136 + *pSKM19 STE5-Myc9*) + EBL231 *GAL1prom-SSA1-TRP1-CEN* | P.M., this study |
| EY3391 (1,2) | EY3379 (EY3141 + pSKM19 STE5-Myc9) + EBL231 *GAL1prom-SSA1-TRP1-CEN* | P.M., this study |
| EY3406 | EYL365 *ste5D::ADE2 + EBL365 pSKM19 STE5-Myc9-URA3-CEN* | P.M., this study |
| EY3407 (1-3) | EY3136 + pSKM12 *STE5-Myc9-URA3-CEN* | P.M., this study |
| EY3409 (1-3) | EY3141 + pSKM12 *STE5-Myc9-URA3-CEN* | P.M., this study |
| EY3410 | YMY117 *((EYL394 ran1-1 ) + pSKM19 STE5-Myc9-URA3-2$\mu$)+* EBL536 *pGAL1prom-SSA1-TRP1-CEN* | P.M., this study |
| LY53,1-3 | EY3136 +pSKM21 *[CUP1promGFP-STE5 URA3 CEN4]* | L.Y., rhis study |
| LY54, 1-3 | EY3141 + pSKM21 *[CUP1promGFP-STE5 URA3 CEN4]* | L.Y., this study |
| EYL1797, 1798 | *bar1D::*$KAN^R$ derivatives of EY3136 and EY3141 *MATa ssa1::HIS3 ssa2::LEU2* | R.M., this study |
| STRAINS-S288C background | | |
| EYL1740 | BY4741 *MATa his3D1 leu2D0 met15D0 ura3D0* | Res Genetics/K/Struhl |
| EYL4860 | *MATa his3D1 leu2D0 met15D0 ura3D0 ydj1D::KanR* | Res Genetics/K.Struhl |
| EYL4862 | *MATa his3D1 leu2D0 met15D0 ura3D0 sti1D::kanR* | Res Genetics/K. Struhl |
| EYL4863 | *MATa his3D1 leu2D0 met15D0 ura3D0 sse1D::kanR* | Res Genetics/K. Struhl |
| EYL4864 | *MATa his3D1 leu2D0 met15D0 ura3D0 sse2D::kanR* | Res Genetics/K. Struhl |

| | | |
|---|---|---|
| EYL4865 | *MATa his3D1 leu2D0 met15D0 ura3D0 ssb1D::kanR* | Res Genetics/K. Struhl<br>l |
| EYL4866 | *MATa his3D1 leu2D0 met15D0 ura3D0 ssz1D::kanR* | Res Genetics/K. Struhl |
| EYL4867 | *MATa his3D1 leu2D0 met15D0 ura3D0 fes1D::kanR* | Res Genetics/K. Struhl |
| EYL BY4038 | *MATa his3D1 leu2D0 met15D0 ura3D0 ste5D::kanR* | Res Genetics/K. Struhl |
| EYL BY3694 | *MATa his3D1 leu2D0 met15D0 ura3D0 msn5D::kanR* | Res Genetics/K. Struhl |
| EYL2809 | YMY564(-1,-2,-3): EYL442 *MATa nsp1ts::URA3 ade2 leu2 lys2 +*<br>*pSKM102[pGal TAgNLS-Ste5 M9 CEN]* | Y.W. this study<br>Y.W. this study |
| EYL2817 | YMY564(-1,-2,-3): EYL442 *MATa nsp1ts::URA3 ade2*<br>*Ura3 leu2 lys2*+pYMW120<br>*pGAL1prom TAgNLS-ste5L610/614/634/637A-Myc9-URA3 -CEN* | Y.W. this study |
| EY3412 | BY4741 *MATa his3D0 leu2D0 met15D0 ura3D0* +<br>pSKM19 *STE5Myc9-URA3-CEN* | P.M. this study |
| EY3417 | EYL1689 BY4741s*te5D0* + pSKM19 *STE5Myc9-CEN* | P.M. this study |
| BY4742 | *RG1104 MATa sst1D0::KANR ura3D0 his3D0 leu2D0 lys2D0* | Research Genetics |
