## Supplementary material for "Effects of HSP70 chaperones Ssa1 and Ssa2 on Ste5 scaffold and the mating mitogen-activated protein kinase (MAPK) Pathway in *Saccharomyces cerevisiae*": TableS2

TableS2 Summary of ImageJ densitometry values relative to wild type.

| Experiment[1] | *ssa1 ssa2* wce[2] / *SSA1 SSA2* wce | IP/input wce |
|---|---|---|
| 1. Fig2A Relative densitometry of ip'd Ste5-Myc9 to co-ip'd 70 kDa band (normalized with IgG) | | |
| –αF 0.67 | | |
| + αF 1.45 | | |
| 2. Relative densitometry of Ste5-Myc9 In wce from *ssa1 ssa2* mutant relative to *MATa* wild type (normalized with Tcm1) | | |
| Fig. 3C Ste5-Myc9 *CEN* MATα *ssa1 ssa2* | 0.20+/-0.03 | |
| Fig.3C Ste5-Myc9 *2μ MATα ssa1 ssa2* | 0.57 +/-0.048 | |
| Fig.3E Ste5-Myc9 CEN *MATa* WT RT | 1 | |
| Fig.3E Ste5-Myc9 CEN *MATa* WT 30$^{o}$C | 0.59 | |
| Fig.3E Ste5-Myc9 CEN *MATa* WT 37$^{o}$C | 0.93 | |
| Fig.3E Ste5-Myc9 CEN *MATa* WT 42$^{o}$C | 0.58 | |
| Fig.3E Ste5-Myc9 CEN *MATa ssa1 ssa2* RT | 0.26 | |
| Fig.3E Ste5-Myc9 CEN *MATa ssa1 ssa2* 30$^{o}$C | ~0.00016 | |
| Fig.3E Ste5-Myc9 CEN *MATa ssa1 ssa2* 37$^{o}$C | ~0.00037 | |
| Fig.3E Ste5-Myc9 CEN *MATa ssa1 ssa2* 42$^{o}$C | ~0.00027 | |
| Fig.3F Ste5-Myc9 CEN *MATa* WT RT | 1 | |
| Fig.3F Ste5-Myc9 CEN *MATa* WT 30$^{o}$C | 1 | |
| Fig.3F Ste5-Myc9 CEN *MATa* WT 37$^{o}$C | 1.5 | |
| Fig.3F Ste5-Myc9 CEN *MATa ssa1 ssa2* RT | 0.29 | |
| Fig.3F Ste5-Myc9 CEN *MATa ssa1 ssa2* 30$^{o}$C | 0.38 | |
| Fig.3F Ste5-Myc9 CEN *MATa ssa1 ssa2* 37$^{o}$C | 0.36 | |
| Fig.3F Ste5-Myc9 CEN *MATa* WT RT[3] | 0.76 | |
| Fig.3F Ste5-Myc9 CEN *MATa* WT 30$^{o}$C[3] | 1.14 | |
| Fig.3F Ste5-Myc9 CEN *MATa* WT 37$^{o}$C[3] | 2.6 | |
| Fig.3F Ste5-Myc9 CEN *MATa ssa1 ssa2* RT[3] | ~0.031 | |
| Fig.3F Ste5-Myc9 CEN *MATa ssa1 ssa2* 30$^{o}$C[3] | ~0.00027 | |
| Fig.3F Ste5-Myc9 CEN *MATa ssa1 ssa2* 37$^{o}$C[3] | ~0.00016 | |
| 3. Relative densitometry of Ste5-Myc9 wce and ip from *ssa1 ssa2* mutant relative to wild type | | |
| Fig.5A Ste5-Myc9 *2μ* | 0.37 +/-0.061 | 1.7 +/-0.17 |
| Fig.5A Ste5-Myc9 *2μ* | 0.29+/-0.044 | 1.9 +/-0.26 |
| Fig.5B Ste5-Myc9 *2μ* | 0.38 | 1.7 |
| Fig.5D Ste5-Myc9 *CEN* | 0.28 | 2.4 |

4. Ratio of HA3-Ste5 in wce and ip'd

from *ssa1 ssa2* mutant relative to wild type

| | | |
|---|---|---|
| Fig. 5C HA3-Ste5 | 0.14 | 3.0 |
| Fig. 5D HA3-Ste5 | 0.2 | 5.6 |

5. Relative densitometry of Fus3 in wce

From *ssa1 ssa2* compared to wt
(normalized with Tcm1 or cross-reactive band)

| | |
|---|---|
| Fig 4C Fus3 | 1.5 +/-0.16 |
| Not shown | |
| Fus3 | 1.2 +/- 0.14 |

6. Relative densitometry of active Fus3

and active Kss1 in wce from *ssa1 ssa2*
compared to wild type normalized with Tcm1

| | |
|---|---|
| Fig.10B Fus3-P WT 0 min αF | 1 |
| WT 240 min αF | 7.7 |
| *ssa1D ssa2D* 0 min αF | 0.36 |
| *ssa1D ssa2D* 240 min αF | 1.4 |
| Fig.10B Kss1-P WT 0 min αF | 1 |
| WT 240 min αF | 6.9 |
| *ssa1D ssa2D* 0 min αF | 0.02 |
| *ssa1D ssa2D* 240 min αF | 1.2[3] |

| | |
|---|---|
| Fig.10C[4] WT Fus3-P 50 μg wce | 1 |
| Fig.10C *ssa1D ssa2D* Fus3-P 50 μg wce | 0.27 |
| Fig.10C WT Fus3-P 300μg wce | 1.7 |
| Fig.10C *ssa1D ssa2D* Fus3-P300 μg wce | 0.14 |

7. Relative densitometry of Ste7-Myc and Fus3 in wce from *ssa1 ssa2* compared to wild type (normalized with Tcm1 and Ponceau S)

| | | |
|---|---|---|
| Fig.10F | Ste7-Myc | 1.02 +/- 0.14 |
| Fig.10F | Ste7-Myc-P | 1.0+/-0 |
| Fig.10G WT: | | |
| Fig. 10G Fus3 | | 1.02 |
| Fig. 10G Fus3 + αF | | 1.74 |
| Fig. 10G Fus3 + cyh | | 0.87 |
| Fig. 10G Fus3 + cyh + no αF | | 0.97 |
| Fig. 10G Fus3 + cyh + αF | | 0.99 |
| Fig. 10G *ssa1D ssa2D*: | | |
| Fig. 10G Fus3 | | 1.20 |
| Fig. 10G Fus3 + α factor | | 1.37 |
| Fig. 10G Fus3 + cyh | | 1.44 |
| Fig. 10G Fus3 + cyh + no αF | | 0.97 |

| | |
|---|---|
| Fig. 10G Fus3 + cyh + αF | 1.06 |

8. Fig. 10D Relative densitometry of Fus3-HA and Fus3K42R-HA in wce from *ssa1 ssa2* compared to wild type (normalized with total protein input)

| | |
|---|---|
| Fus3-HA in *SSA1 SSA2* | 1.0 +/- 0.0027 |
| Fus3K42R-HA in *SSA1 SSA2* | 1.36 +/- 0.57 |
| Fus3-HA in *ssa1 ssa2* | 3.06 +/- 0.40 |
| Fus3K42R-HA in *ssa1 ssa2* | 2.26 +/- 0.88 |

9. Relative densitometry of Kss1 in wce from *ssa1 ssa2*, *fus3*, *kss1*, compared to wild type of same background (normalized with cross-reactive band)[4]

| | |
|---|---|
| Fig.10E Kss1 in *ssa1 ssa2* | 1.24 |
| Fig.10E Kss1 in *fus3D* | 2.59 |
| Fig.10E Kss1 in *kss1D* | 0 |

---

[1] Summary of relative ImageJ densitometry values of Ste5, Fus3 and Ste7, and 70kDa protein in extracts from *ssa1 ssa2* mutants compared to wild type *SSA1 SSA2.* Experiments are from the figures in the main text using images shown and shorter or longer exposure images that are not shown. See Materials and Methods for densitometry methods with ImageJ, photoshop and canvasXdraw. Some values from the *ssa1 ssa2* mutant were equal to or below the background. Negative values were multiplied by -1000 to yield a positive integer equal to or greater than ~1 for the calculations. ImageJ densitometry was done according to the directions from the NIH and from SYBIL (European Community's Seventh Framework Programme FP7/2007-2013 at www.sybil-fp7.eu/node/95) and reference S10.

[2]Abbreviations: WCE is whole cell extract, IP is immunoprecipitated, co-ip is co-immunoprecipitated, Ste7-Myc-P is the phosphorylated slower migrating species.

[3]These values are from the second set of lanes in Fig.3C. Due to degradation, the entire lane was used for quantification.

[4]These values reflect the reduction of detectable active Fus3 when large amount of whole cell extract is run on SDS-polyacrylamide gel.

[4]The pattern appeared similar for repeat experiments, however, antibody background precluded reliable densitometry. Higher levels of background led to a lower fold increase.
