## Supplementary material for "Effects of HSP70 chaperones Ssa1 and Ssa2 on Ste5 scaffold and the mating mitogen-activated protein kinase (MAPK) Pathway in *Saccharomyces cerevisiae*": TableS3

Table S3. Localization of Ste5 derivatives to cell periphery and nucleus in wild type and *ssa1 ssa2* cells.

| Strain[1] | %Rim[2] | %N>C[3] |
|---|---|---|
| 1. *STE5promSTE5-MYC9, RT, 2μ,* 50 nM αF 45 min. | | |
| WT | 29.3 (N~150) | |
| *ssa1 ssa2* | 16.8 (N~150) | |
| 2. *STE5promTAgNLS$^{K128T}$-STE5-MYC9 2μ, RT,* 50 nM αF 20 min. | | |
| WT | 40.7 (N~250) | |
| *ssa1 ssa2* | 14.2 (N~250) | |
| 3. *TAgNLS-GFP2, TAgNLS-NES-GFP2*, RT, no αF | | |
| *WT* TAgNLS-GFP2 | | 95.0 (N=60) |
| *ssa1 ssa2* TAgNLS-GFP2 | | 96.0 (N=200) |
| *WT* TAgNLS-NES-GFP2 | | 1.5 (N=200) |
| *ssa1 ssa2* TAgNLS-NES-GFP2 | | 0 (N=200) |
| 4. *TAgNLSNESGFP2*, live cells fixed, RT, 55°C 2 hr | | |
| WT RT | | 1.5 (N=200) |
| WT 55°C | | 0[4] |
| *ssa1 ssa2* RT | | 0 (N=200) |
| *ssa1 ssa2* 55°C | | 0[4] |
| 5. *Ste5(1-242)-GFP2,* live cells fixed, RT, 55°C 2 hr | | |
| WT RT | | 93.2 (N=206) |
| WT 55°C | | 0[4] |
| *ssa1 ssa2* RT | | 0.0068 (N =148) |
| *ssa1 ssa2* 55°C | | 0[4] |

[1]All strains were grown and prepared for microscopy by PBM. Data in 1-3 were tallied by PBM, data in 4-5 were tallied by RM. Strains EY3136 *MATa SSA1 SSA2* ("wild type", WT) and EY3141 *MATa ssa1::HS3 ssa2::LEU2* harboring plasmids pSKM19 STE5-MYC9, TAgNLSK128T-STE5-MYC9, EBL663 TAgNLS-NES-GFP2 or pYM27.1 TAgNLS-GFP2, Ste5(1-242)-GFP2, were grown in SC-uracil medium with 2% dextrose at room temperature (RT) to logarithmic phase and monitored live for GFP fluorescence. Cells harboring pSKM19 (*STE5-MYC9 URA3 2μ)* and pSKM98 *TAgNLSK128T-STE5=MYC9* were grown with shaking at room temperature in SC-uracil with 2% dextrose overnight to an $A_{600}$ of 0.3-0.6, then cells were adjusted to $A_{600}$ ~0.5 with or without α factor. Cells were then fixed and prepared for indirect immunofluorescence with 9E10 monoclonal antibody and DAPI. Aproximately 211-379 cells were tallied per sample. S.E. is standard error. (N) is the number of cells counted in the individual experiments

[2]The %Rim is the percentage of 9E10 positive cells with a signal that is more intense at the cell periphery than the surrounding cytoplasm.
[3]The %N>C is the percentage of total 9E10 positive cells and GFP positive cells with more intense staining in the nucleus compared to the cytoplasm.
[4] At 55$^{o}$C the green GFP signal started to disappear and become a yellow-green signal that matched the weak background yellow-green signal of plasmid control cells that did not harbor any GFP. The fields of cells permitted counting over 100 cells but it was difficult to get a good count.
