## Supplementary material for "Effects of HSP70 chaperones Ssa1 and Ssa2 on Ste5 scaffold and the mating mitogen-activated protein kinase (MAPK) Pathway in *Saccharomyces cerevisiae*": TableS4

Table S4. Summary of growth and mating of *ssa1*, *ssa2*, *ssa3*, and *ssa4* single, double and triple mutants.

| | Growth | | Mating | |
|---|---|---|---|---|
| Strain[1] | RT | 30$^{o}$C | RT | 30$^{o}$C |
| EY3136 *WT SSA1SSA2 SSA3 SSA4* | ++ | ++++ | +++ | +++ |
| EY3137 *ssa1* | + | +++ | | +++ |
| EY3138 *ssa2* | + | +++ | | +++ |
| EY3139 *ssa3* | + | +++ | | +++ |
| EY3140 *ssa4* | + | +++ | | +++ |
| EY3141 *ssa1 ssa2* | +/- | +/- | +++ | +++ |
| EY3143 *ssa1 ssa4* | + | | | |
| EY3144 *ssa2 ssa3* | + | | | |
| EY3145 *ssa2 ssa4* | + | | | |
| EY3146 *ssa3 ssa4* | + | | +++ | |
| EY3147 *ssa1 ssa2 ssa3* | +/-- | | +++ | |
| EY3148 *ssa1 ssa3 ssa4* | + to ++ | | +++ | |
| EY3149 *ssa2 ssa3 ssa4* | + | | +++ | |

[1] Streakouts and patch mating of WT and *ssa1-ssa4* single, double and triple mutant strains. WT and *ssa1-ssa4* single mutant and *ssa1 ssa2* double mutant strains were tested for mating at 30$^{o}$C after mating for 6 hours on YPD followed by recovery of diploid prototrophs on YNB 2% dextrose at 30$^{o}$C. The. WT, *ssa1-ssa4* double mutant and triple mutant strains were quite sick and were mated at room temperature for 3 hours on YPD. In these experiments extra cells of the two slowest growing strains, *ssa1 ssa2* and *ssa1 ssa2 ssa3,* so that these patches would be of equal density to wild type patches after overnight growth at either 30$^{o}$C or room temperature. WT and *ssa1 ssa2* strains were tested in mating assays four times and the ssa1-ssa4 single mutants and all other double and triple mutants were tested twice. Strains tested in these assays were: *EY3136 MATa wild type, EY3137 MATa ssa1::HIS3, EY3138 MATa ssa2::LEU2, EY3139 MATa ssa3::TRP1, EY3140 MATa ssa4::URA3,, EY3141 MATa ssa1::HIS3 ssa2::LEU2, EY3143 MATa ssa1::HIS3 ssa4::URA3, EY3144 MATa ssa2::LEU2 ssa3::TRP1, EY3145 MATa ssa2::LEU2 ssa4::URA3,, EY3146 MATa ssa3::TRP1 ssa4::URA3, EY3147 MATa ssa1::HIS3, ssa2::LEU2 ssa3::TRP1, EY3148 MATa ssa1::HIS3 ssa3::TRP1 ssa4::URA3,, EY3149 MATa ssa2::LEU2 ssa3::TRP1. ssa4::URA3.* Plates were photographe every day for 4 days.
