## Supplementary material for "Effects of HSP70 chaperones Ssa1 and Ssa2 on Ste5 scaffold and the mating mitogen-activated protein kinase (MAPK) Pathway in *Saccharomyces cerevisiae*": TableS5

Supplemental Table 5. List of positive and negative regulators of mating and invasive growth screened for interaction with Ssa1, Ssa2 and other HSP70 network proteins in the literature and published databases.

PROTEIN

Act1/YFL039C***,
Adf1/YCL058W-A*
Aga1/YNR044W*
Aga2/YGL032C*
Akr1/ YDR264C***
Arr4/Get3/YDL100C*
Ark1/YNL020C***
Ash1/YKL185W**
Axl1/YIL140W*
Axl2/Bud10/YPR122W***
Bck1/Slk1/YJL095W***
Bem1/YBR200W***
Bem2/YER155C***
Bem3/YPL115C***
Bem4/YPL161C***
Bni1/YNL271C***
Bos1/Sec32/YLR078C*
Bud2/YKL092C***
Bck1/Slk1/YJL095W***
Bem1/YBR200W***
Bem2/YER155C***
Bem3/YPL115C***
Bem4/YPL161C***
Bni1/YNL271C***
Bos1/Sec32/YLR078C*
Bud2/YKL092C***
Bud3/YCL014C**
Bud5/YCR038C***
Bud6/YLR318C***
Bud8/YLR353W***
Ccw12/YLR110C*
Cdc4/YFL009W***
Cdc24/YAL041W***
Cdc28/YBR160W***
Cdc34/YDR054C***

Cdc36/YDL165W***
Cdc37/YDR168W***
Cdc39/YCR093W***
Cdc42/YLR229C***
Cdc48/YDL126C*+
Cdc53/YDL132W***
Cdc55/YGL190C***
Chs3/YBR023C*
Chs5/YLR330W*+
Cik1/YMR198W*
Cin2/YPL241C*
Cin4/YPL241C*
Cla4/YNL298W***
Cla4/YNL298W***
Cln1/YMR199W***
Cln2/YPL256C***
Cln3/YAL040C***+
Cmd1/YBR109C*
Cmp2/YML057W*
Cna1/YLR433C*
Cnb1/YKL190W*
Csi1/YMR025W*
Csn9/YDR179C*
Csn12/YJR084W*
Dia1/YMR316W**
Dia3/YDL024C*
Dig1/Rst1/ YPL049C***+
Dig2/Rst2/ YDR480W***
Dnf1/YER166W*
Dnf2/YDR093W*
Dnf3/YMR162C*
Drs2/YAL026C*
Elm1/YKL048C***
Far1/YJL157C*
Far3/YMR052W*
Far7/YFR008W*
Far8/YMR029C*
Far9/VPS64/YDR200C*
Far10/YLR238W*
Far11/YNL127W*

Fig1/YBR040W*
Fig2/YCR089W
Fig3/Kar5/YMR065W
Fig4/YNL325C*
Fks1/YLR342W***+
Fks2/YGR032W***
Flo8/YER109C**
Fps1/YLL043W***+
Fpk1/YNR047W***
Fus1/YCL027W*
Fus2/YMR232W
Fus3/YBL016W***
Fyv5/YCL085C***
Gic1/Ira1/YBR140C***,
Glc7/YER133W***+
Gpa1/Cdc70/Dac1/Scg1/YHR005C***
Gpa2/YER020W**
Hbt1/YDL223C*
HO/YDL227C*
Hkr1/YDR420W***
Hub1/YNR032C***
Hyp2/TIF51A/YEL034W***
Kar1/YNL188W*
Kar2/YJL034W***+
Kar3/ YPR141C*
Kar4/YCL055W*
Kar5/Fig3/YMR065W*
Kar7/Sec66/ YBR171W***
Kar8/Jem1/YJL073W*
Kar9/YPL269W*
Kel1/YHR158C***+
Kel2/YGR138C***
Kem1/Xrm1/YGL173C***+
Kin28/YDL108W**
Kin82/YCR091W*
Kss1/YGR040W***
Lsg1/YHL003C*
Mdg1/YNL137C*
Mdy2/YOL111C*+
Mfa1/YDR461W*

Mfa2/YNL145W*
Mf(alpha1)/YPL187W***
Mf(alpha2)/YNL145W***
Mfg1/YDL233W*
Mid2/YLR322W*
Mkk1/YOR231W*
Mkk2/Ypl143C*
Mpk1/Slt2/YHR030C*
Mps3/YJL019C***
Mpt5/YGL178W***
Msb1/YOR188W***
Msb2/YGR014W***
Msg5/YNL053W***
Mss11/YMR164C**
Myo2/YOR326W***
Myo4/YAL029C*
Myo5/YMR109W***
Nfg1/YLR042C**
Och1/YGL038C**
Opy2/YGR014W**
Osh3/YHR073W*
Pea2/YER149C**
Pkc1/YBL105C***
Pog1/YIL122W***
Pph21/YDL134C
Pph22/YDL188C***
Prm1/YNL279W*
Prm2/YIL037C*
Prm3/YPL192C*
Prm4/YPL156C*
Prm5/YIL117C*
Prm6/KCH1/YML047C*
Prm7/YDL039C*
Prm8/YGL053W*
Prm9/YAR031W*
Prm10/YJL108C*
Prm15/YMR278W*
Prr1/YKL116C*
Prr2/YDL214C*
Ptc1/YDL006W***

Ptc2/YER089C***
Ptp1/YDL230W***
Ptp2/YOR208W***
Ptp3/YER075C***
Ras2/YNL098C***
Rck1/YGL158W*
Rck2/YLR248W***
Reg1/YDR028C*
Rga1/YOR127W***
Rga2/YDR379W***
Rgd1/YBR260C*
Rho1/YPR165W***
Rod1/YOR018W***
Rog3/YFR022W*
Rri1/YDL216C*
Rri2/YOL117*
Rsp5/YER125W***+
Rsr1/Bud1/YGR151C***
Rvs161/YCR009C***+
Rsv167/YDR388W***+
Rxt2/YBR095C***+
Sag1/YJR004C*
Sak1/YER129W*
Scp160/YJL080C***+
Scw10/YMR305C*
Skp1/YDR328C***+
Sey1/YOR165W*
Sfg1/YOR315W***
Sho1/YER118C**
Spa2/ YLL021C***
Spc110/YDR356W*+
Sph1/ YLR313C***
Spo14/YKR031C*
Spt7/YBR081C***
Ssf1/YHR066W*
Ssf2/YDR312W***
Sst1/Bar1/YIL015W*
Sst2/YLR452C*
Ste2/YFL026W*
Ste3/YKL178C***

Ste4/YOR212W***
Ste5/YDR103W***
Ste6/YKL209C*
Ste7/YDL159W***
Ste11/YLR362W***
Ste12/YHR084W***
Ste13/YOR219C*
Ste14/YDR410C*
Ste16/Ram1/Fus8/YDL090C*
Ste18/YJR086W*
Ste20/YHL007C***
Ste21/Msn5/YDR335W***
Ste23/YLR389C*
Ste24/YJR117W*
Ste50/YCL032W***
Ste24/YJR117W*
Ste50/YCL032W***
Sut1/YGL162W***
Sut2/YPR009W***
Swi4/YER111C***
Swi6/YLR182W***
Tec1/ YBR083W**
Tem1/YML064C***
Tip1/YBR067C***
Tos3/YGL179C*
Ubc1/YDR177C***+
Ubi1/RPL40A***
Whi2/YOR043C***
YNL058C*
Ypk1/YKL126W*+
Ypk2/YMR104C***
Yps1YLR120CC/***
Yps2/Mck2/YDR144C***
Ysp1/YHR155W***

---

[1]This is a list of ~ 212 mating pathway proteins* and ~121 invasive growth pathway proteins** compiled from SGD, the literature and Integrative Multi-species Prediction (IMP) (S30-S39). The list does not include every protein identified by SGD GO terms, but does include the majority of

core signaling proteins, transcription factors, cell cycle, agglutination, morphogenesis, nuclear fusion and cell fusion, and interaction analysis.
[2]Asteriks: Mating pathway*, invasive growth pathway**, both***, interacts with Ssa1 + alpha-factor.
