## Supplementary material for "Effects of HSP70 chaperones Ssa1 and Ssa2 on Ste5 scaffold and the mating mitogen-activated protein kinase (MAPK) Pathway in *Saccharomyces cerevisiae*": TableS6

Table S6. Summary of interaction data on 227 mating pathway and invasive growth pathway proteins with Hsp70 chaperones Ssa1, Ssa2, Ssa3, Ssa4 and other Hsp70 network proteins.

| Protein | Description/Putative interacting proteins[1] |
|---|---|
| Ste5: | Ssa1, Ssb1, Ssb2 |
| Ssa1: | *Classic Hsp70 chaperone in cytoplasm*<br>Act1, Akr1, Arr4/Get3, Ash1, Axl1, Akr1, Bar1/Sst1, Ben1, Bem2, Bem4, Bud3, Bni1, Bos1/Sec32, Ccw12, Cdc4, Cdc24, Cdc28, Cdc34, Cdc37, Cdc39, Cdc42, Cdc48, Cdc55, Chs3, Chs5, Cik1, Cin2, Cla4, Cln3, Cmp2, Cna1, Cnb1, Cnb2, Csn9, Dia1, Dia2, Dia3, Dig1, Dig2, Dnf3, Elm1, Far1, Far3, Far7, Far8, Far11, Fig2, Fig4, Fks1, Fus1, Fus2, Fus3, Gic1, Glc7, Hbt1, HO, Kar2, Kar3, Kar4, Kel1, Kem1/Xrm1, Kin28, Kin82, Lsg1, Mdy2, Mkk1, Mpt5, Msg5, Mss11, Myo2, Myo5, Osh2, Osh3, Pkc1, Pog1, Pph22, Prm9, Prr1, Prr2, Ras2, Rck2, Reg1, Rga1, Rga2, Rgd1, Rod1, Rsp5, Rsr1/Bud1, Rsv161, Rvs167, Rxt2, Sac7, Sak1, Scp160, Sey1, Sfg1, Spc110, Sph1, Spo14, Spt7, Ssf1, Sst2, Ste2, Ste4, Ste5, Ste6, Ste11, Ste12, Ste13, Ste18, Ste20, Ste21/Msn5, Ste24, Ste50, Sut1, Tec1, Tip1, Tos3, YNL058C, Ypk1, Ysp1, [Fes1, Ssa2, Ssa3, Ssa4, Ssb1, Ssb2, Ssc1, Sse1, Sse2, Ssz1, Kar2, Sil1, Sti1,Whi2, Ydj1, Hsp26, Hsc82, Hsp82, Hsp104] |
| Ssa1 + $\alpha$F: | Dig1, Glc7, Kel1, Cdc28, Cdc48, Cdc53, Cdc55, Chs5, Cln3, Dig1, Fks1, Fps1, Glc7, Kel1, Kem1/Xrm1, Mdy2, Rsp5, Rvs161, Rvs167, Rxt2, Scp160, Skp1, Spc110, Ste12, Ubc1, Whi2, Ypk1, [Ssb1, Ssc1, Sse2, Ssz1, Kar2, Sis1, Ydj1, Fes1, Sti1, Hsp26, Hsc82, Hsp82, Hsp104] |
| Ssa2: | *Classic Hsp70 chaperone in cytoplasm*<br>Akr1, Axl2/Bud10, Bem2, Bni1, Bos1/Sec32, Bud3, Cdc4, Cdc28, Cdc24, Cdc28, Kel1, Cdc34, Cdc48, Cdc55, Chs3, Cik1, Cin2, Cla4, Cln2, Cln3, Cmp2, Cnb1, Cnb2, Csn9, Dia2, Dia3, Dig1, Dnf1, Dnf3, Elm1, Far1, Far7, Far8, Far11, Fig2, Fig4, Fks1, Fus1, Fus2, Fus3, Gic1, Hbt1, HO, Kar1, Kar2, Kem1/Xrm1, Kin82, Mpt5, Msb1, Mss11, Myo2, Myo5, Osh3, Pea2, Pkc1, Pog1, Prm9, Prr1, Ras2, Rck1, Rck2, Rga1, Rga2, Rod1, Rog3, Rri2, Rsp5, Rvs161, Rvs167, Rxt2, Sak1, Scp160, Scw10, Spc110, Sph1, Spo14, Spt7, Ssf1, Sst2, Ste2, Ste6, Ste11, Ste12, Ste13, Ste21/Msn5, Ste50, Tec1, Tos3, YNL058C, Whi2, Ypk1 [Ssa1, Ssa2, Ssa3, Ssa4, Ssb1, Ssb2, Sse1, Sse2, Fes1, Ydj1, = Kar2, Hsp42] |
| Ssa3: | *Classic Hsp70 chaperone in cytoplasm*<br>Bni1, Bos1/Sec32, Bud2, Bud3, Cdc34, Cdc39, Cln3, Dia2, Elm1, Gic1, Glc7, Kar1, Kar2, Kar3, Osh2, Pph22, Spa2, Spc110, Ste2, Ste11, Ste24, Swi4 [Ssa1, Ssa2, Ssa3, Ssa4, Ssb1, Ssb2, Sse1, Sse2, Ydj1, Kar2] |
| Ssa4: | *Classic Hsp70 chaperone in cytoplasm*<br>Bud3, Cdc28, Cdc37, Cla4, Cln2, Cln3, Cna1, Far3, Fig2, Fus1, Gic1, Kar2, Kel1, Kem1/Xrm1, Msg5, Osh2, Pkc1, Pph22, Rod1, Rri1, Rvs161, Spa2, Spc110, Sst2, Ste12, Ste16/Ram1, Ste18, Ste21/Msn5, Sst2, Swi4, Tec1, Ysp1, [Ssa1, Ssa2, Ssa3, Ssb1, Sse1, Sse2, Fes1, Ydj1] |
| Ssb1: | *Hsp70 chaperone associated with ribosomes (paralog of Ssb1)*<br>Act1, Ark1, Bar1/Sst1, Bck1/Sik1, Bem2, Bem4, Bni1, Bos1, Bud3, Bud4, Bud6, Ccw12, Cct6, Cct8, Ccw12, Cdc4, Cdc24, Cdc28, Cdc34, Cdc37, Cdc39, Cdc42, |

Cdc43, Cdc48, Cdc53, Cdc55, Chd1, Chs3, Chs5, Cik1, Cka1, Cka2, Cin2, Cin4, Cla4, Clb2, Cln2, Cln3, Cmd1, Cmp2, Cmp2, Cna1, Cnb1, Csn9, Csn12, Dia2, Dia3, Dig1, Dig2, Dnf1, Dnf3, Elm1, Far1, Far7, Far8, Far11, Fig2, Fig4, Flo8, Fks3 , Fus3, Gic1, Glc7, Gpa2, Hub1, Hyp2, Kar2, Kar3, Kel1, Kel2, Kem1, Kin28, Lsg1, Mid2, Mkk1, Mkk2, Mpk1/Slt2, Mpt5, Msb1, Msg5, Mss11, Myo3, Myo4, Myo5, Osh2, Osh3, Pea2, Pkc1, Pkh1, Pog1, Prm9, Pph21, Pph22, Prr1, Prr2, Ptc2, Ptk1, Ptp1, Ras2, Rck1, Rck2, Rga1, Rga2, Reg1, Rgd1, Rgr1, Rod1, Rri1, Rri2, Rvs161, Rvs167, Rsp5, Rxt2, Sak1, Scp160, Scw10, Sfg1, Spc110, Spo14, Ssf1, Sst2, Spo14, Spt7, Ste2, Ste3, Ste5, Ste6, Ste7, Ste11, Ste13, Ste20, Ste21/Msn5, Ste24, Ste50, Sut1, Tec1, Tem1, Tip1, Tos3, Ubc1, Whi2, Ypk2, Ysp1, [ Fes1, Hsc82, Hsp42, Hsp104, Hsp82, Kar2, Sis1, Snl1, Ssa1, Ssa2, Ssa3, Ssa4, Ssb1, Ssb2, Sse1, Sse2, Ssc1, Ssz1, Sti1, Ydj1]

Ssb2: *Hsp70 chaperone associated with ribosomes, paralog of Ssb1*
Act1, Aga2, Bem1, Bem2, Bem4, Bni1, Bud3, Ccw12, Cdc4, Cdc24, Cdc28, Cdc34, Cdc36, Cdc37, Cdc39, Cdc42, Cdc48, Cdc53, Cdc55, Chs3, Cik1, Cin2, Cin4, Cla4, Clb2, Cln1, Cln2, Cmd1, Cmp2, Cna1, Cnb1, Csn9, Dia2, Dig1, Dnf3, Far1, Far10, Far11, Gic1, Glc7, Hub1, Kar4, Kel1, Kin28, Kss1, Mdy2, Mid2, Mkk1, Mkk2, Mpt5, Msb2, Myo3, Myo4, Myo5, Nfg1, Opy2, Osh2, Pea2, Pkc1, Pog1, Pph21, Pph22, Ptc1, Ptc2, Ptc3, Ptp1, Ptp3, Reg1, Rho1, Far1, Far10, Fus3, Rck1, Reg1, Rho1, Rod1, Rri1, Rsp5, Rsr1/Bud1, Rvs161, Rvs167, Sac1, Scp160, Scw10, Skp1, Sph1, Spo14, Sst2, Ste2, Ste4, Ste5, Ste7, Ste11, Ste20, Ste21/Msn5, Ste23, Sut2, Tec1, Tem1, Tos3, Ubc1, Whi2, Ysp1 [ Fes1, Hsp26, Hsp104, Hsp82, Sis1, Snl1, Ssa1, Ssa2, Ssa3, Ssb1, Ssb2, Ssc1, Sse1, Sse2, Ssz1, Ydj1]

Ssz1: *Hsp70 chaperone in cytoplasm*
Bem2, Bud2, Bud3, Cdc4, Cdc39, Cla4, Cna1, Dia2, Dnf3, Ecm22, Elm1, Far1, Far3, Far7, Kem1/Xrm1, Kar5, Kem1/Xrm1, Kss1, Lsg1, Mkk2, Osh2, Osh3, Pkc1, Prr1, Ptc1, Rck1, Reg1, Scp160, Spt7, Sst2, Tec1, Ubi1/RPL40A, Ysp1, Ysp2, [Ssb1, Ssb2, Ssc1, Sse1, Ssq1, Hsc82, Hsp82, Sti1]

Sse1: *Hsp110/ATPase component/nucleotide exchange factor for Hsp70 and Hsp90*
Act1, Bck1/Sik1, Bem2, Bni1, Bos1/Sec32, Bud2, Bud3, Ccw12, Cdc4, Cdc24, Cdc34, Cdc39, Cdc42, Cdc48, Cdc55, Chs3, Chs5, Cin2, Cln2, Cla4, Cmd1, Dia1, Dia2, Dig2, Dnf3, Drs2, Elm1, Far1, Fus1, Far1, Far7, Far8, Gic1, Glc7, Hbt1, Hub1, Kar2, Kel1, Kel2, Kem1/Xrm1, Kin28, Kss1, Mkk1, Mpk1/Slt2, Msb1, Msg5, Msn5, Mss11, Myo2, Myo5, Osh3, Pkk1, Pog1, Pph21, Prm1, Prm5, Prr1, Prr2, Prs1, Ptp3, Ras2, Reg1, Rga1, Rga2, Rod1, Rri2, Rvs161, Rvs167, Rxt2, Sak1, Scp160, Spa2, Sph1, Spo14, Ste2, Ste20, Ste21/Msn5, Ste50, Sut1, Tem1, Tip1, Tos3, Whi2, YNL058C, Ypc1, Ypk1, [Ssa1, Ssa2, Ssa4, Ssb1, Ssb2, Sse1, Sse2, Hsp42, Hsc82, Hsp80, Hsp104, Sis1, Sti1]

Sse2: *Hsp110/putative ATPase component/nucleotide exchange factor for Hsp70, Hsp90*
Bck1, Bud2, Cdc34, Cdc48, Chs3, Chs5, Dia1, Dia2, Elm1, Far1, Far11, [Ssa1, Ssa2, Ssa3, Ssa4, Ssb1, Ssb2, Sse1, Fes1, Ydj1, Hsp42, Hsc82, Hsp82, Hsp104, Sti1, Ydj1]

Ssc1: *Hsp70 ATPase of translocase of inner mitochondrial membrane (paralog Ecm10)*

Bem2, Bud2, Cdc24, Cdc28, Cdc34, Cdc55, Elm1, Far7, Far8, Fpk1, Hbt1, Lsg1, Myo4, Pph21, Ste7, Ste11, Scp160, Spo14, Spt7, Ssf1, Ypk1, Ypk2, [Ssa1, Ssa2, Ssa4, Ssb1, Ssb2, Ssc1, Ssq1, Ssz1, Ecm10, Hsp42, Hsp82, Hsp104, Fes1]

Ecm10: *Hsp70 ATPase chaperone in mitochondrial nucleoids*
Bud6, Cln2, Pph21, Pph22, Ste2, Ste24, [Ssa4, Ssc1]

Ssq1: *Hsp70 chaperone in mitochondria*
Mss11, Pph21, Rsv167, [Ssa1, Ssa2, Ssb1, Ssc1, Ssz1]

Kar2: *Hsp70 chaperone in ER and for protein translocation across ER*
Arr4/Get3, Cdc28, Chs5, Far3, Far8, Ste24, Kel1, Mdy2, Prm2, Sac7, Scp160, Skp1, Ste24, [Ssa1, Ssa2, Ssa3, Ssa4, Ssb1, Ssb2, Sse1, Kar2, Hsp82]

Ydj1: *Hsp40/DnaJ family chaperone chaperone - stimulates Hsp70 ATPase activity, promotes re-folding and/or proteosomal degradation of client proteins*
Act1, Ash1, Bck1/Sik1, Bem2, Bni1, Bos1/Sec32, Bud1/Rsr1, Bud2, Bud6, Cdc4, Cdc28, Cdc39, Cdc48, Chs3, Cla4, Cln1, Cln3, Cmd1, Cna1, Dia2, Dnf2, Dnf3, Far3, Gpa1, Kel1, Kel3, Msn5/Ste21, Mdy2, Mpt5, Myo2, Osh2, Pkc1, Pph22, Prr1, Rck1, Rga1, Rga2, Rri2, Scp160, Spa2, Spc110. Spo14, Sst2, Ste4, Ste6, Ste11, Ste12, Swi6, Tos3, Ubi4, Ypk1, [Ssa1, Ssa2, Ssa3, Ssa4, Ssb1, Sse1, Sse2, Hsp26, Hsp42, Hsc82, Hsp82, Sis1, Ydj1]

Sis1: *Hsp40/DnaJ family co-chaperone*
Bck1/Sik1, Bos1, Bud3, Cdc4, Cdc34, Cdc37, Cdc39, Chs5, Cmd1, Csn9, Dia2, Dig1, Far1, Far3, Far11, Far7, Fig2, Flo8, Glc7, Mss11, Myo2, Myo5, Pkc1, Ptc2, Rod1, Rri1, Sak1, Scp160, Sey1, Spt7, Ste2, Ste12, Ste20, Ste21/Msn5, Tec1, Whi2, [Ssa1, Ssa2, Ssb1, Ssb2, Sse1, Hsp42, Ydj1, Sti1]

Fes1: *Hsp70-J protein nucleotide exchange factor, promotes release of client from Hsp70 to ubiquitin-proteasome, also functions with Hsp90s*
Fbp1, Lsg1, Mpk1/Slt2 [Ssa1, Ssa2, Ssa4, Ssb1, Ssb2, Sse1, Hsp42, Sis1]

Snl1[2]: *Hsp70 co-chaperone*
Mpt5, [Ssa1, Ssb1, Ssb2]

Sti1[2]: *Hsp90 co-chaperone that links to Hsp70*
Cdc37, Scp160, Ubi4 [Ssa1, Ssa2, Ssa3, Ssa4, Ssb1, Ssb2, Sse1, Sse2, Ssz1, Hsc82, Hsp82, Hsp104, Sis1, Sti1]

Hsp26: *Small Hsp chaperone*
Bni1, Far1, Fus3, Cdc28, Cmr1, Spt7, Ste2, Ste50, [Ssa1, Ssa2, Ssb1, Ssb2, Ydj1]

Hsp42: *Small Hsp chaperone*
Bem2, Cdc3, Chs3, Cik1, Dia2, Dnf3, Fus3, Fes1, Gic1, Myo2, Myo5, Rvs161, Sfg1, Ypk1, Ypk2,Ysp1, [Ssa1, Ssa2, Ssb1, Ssc1, Sse1, Sse2, Hsc82, Hsp82, Hsp104, Sis1, Fes1, Ydj1]

Hsp104: *Hsp100 chaperone*

Matalpha1, Bem2, Bni1, Cdc4, Cdc34, Cdc48, Cdc53, Chs3, Cln2, Dig2, Dnf3, Far1, Far3, Far11, Fks1, Fus1, Gpa1, Ho, Mss11, Pkc1, Rga1, Rog3, Scp160, Skp1, Spa2, Ste24, Ysp1, [Ssa1, Ssb1, Ssb2, Ssc1, Sse1, Sse2, Hsp42, Hsc82, Hsp104, Ydj1]

---

[1]The databases of Gong (S30), Truman (S31), Biogrid (SGD)(S32), IntAct (Uniprot)(S33), Babu (S32), Willmund (S34) and literature (S35-39) were screened for mating and invasive growth pathway proteins that interact with HSP70 components (Table S5 has a list of mating and invasive growth pathway proteins). The descriptions are based on the literature and SGD.

[2]Not in the Gong database which has the majority of known HSP70 interactions (S30).
