## Supplementary material for "Effects of HSP70 chaperones Ssa1 and Ssa2 on Ste5 scaffold and the mating mitogen-activated protein kinase (MAPK) Pathway in *Saccharomyces cerevisiae*": TableS7

Table S7. Cell morphology of Hsp70 family mutants before and after α factor treatment.

| Strain[1] | αF (μM) | N[2] | %RUB | %B | %EI | %S | %LS | %MS | %CS | %DS | %BB | %TS | %UCE | %WT TS |
|---|---|---|---|---|---|---|---|---|---|---|---|---|---|---|
| Wild type1 | 0 | N>300 | 41.4 | | 0 | 0 | 0 | | | | | | | 0 |
| Wild type1[3] | 5 | N>300 | | | 5.2 | 94.9 | 0 | | | | | | 100 | 100 |
| *ssa1 ssa2*[3] | 0 | N>300 | 56.2 | | 0 | 0 | 0 | | | | | | | 0 |
| *ssa1 ssa2*[3] | 5 | N>300 | | | 67.4 | 32.6[2] | 0 | | | | | | 100 | 34.4 |
| Wild type2 | 0 | 392 | 43.6 | 56.4 | 0 | 0 | 0 | 0 | 0 | 0 | 0 | 0 | 0 | 0 |
| Wild type2 | 2.5 | 483 | 21.3 | 1.0 | 25.9 | 36.0 | 15.7 | 0 | 0 | 0 | 0 | 51.7 | 77.6 | 100 |
| Wild type2 | 5.0 | 944 | 14.9 | 0.02 | 25.0 | 54.2 | 0 | 1.7 | 0 | 0 | 0 | 55.9 | 80.9 | 100 |
| *ydj1* | 0 | 381 | 53.9 | 47.0 | 0 | 0 | 0 | 0 | 0 | 0 | 0 | 0 | 0 | 0 |
| *ydj1* | 2.5 | 549 | 15.5 | 1.8 | 36.2 | 24.2 | 0 | 0 | 8.4 | 0 | 13.8 | 32.6 | 68.8 | 88.9 |
| *ydj1* | 5.0 | 482 | 18.9 | 0.6 | 35.7 | 22.8 | 0 | 0.2 | 12.7 | 0 | 9.1 | 35.9 | 71.4 | 64.2 |
| *fes1*[4] | 0 | 2199 | 60.7 | 34.9 | 3.3 | 1.1 | 0 | 0 | 0 | 0 | 0 | 1.1 | 4.4 | 0 |
| *fes1*[4] | 2.5 | 2146 | 34.2 | 1.4 | 33.0 | 30.7 | 0 | 0.6 | 0 | 0 | 0 | 31.3 | 64.3 | 56.0 |
| *snl1* | 0 | 1094 | 39.9 | 60.0 | 0 | 0 | 0 | 0 | 0 | 0 | 0 | 0 | 0 | 0 |
| *snl1* | 5.0 | 780 | 28.9 | 1.1 | 23.9 | 30.5 | 13.5 | 0 | 0 | 0 | 0 | 44.0 | 68.0 | 78. 7 |
| *sti1* | 0 | 563 | 48.8 | 51.1 | 0 | 0 | 0 | 0 | 0 | 0 | 0 | 0 | 0 | 0 |
| *sti1* | 5.0 | 462 | 6.7 | 0 | 7.4 | 81.8 | 1.7 | 2.4 | 0 | 0 | 0 | 85.9 | 93.3 | 153.6 |
| *ssb1* | 0 | 494 | 46.4 | 53.6 | 0 | 0 | 0 | 0 | 0 | 0 | 0 | 0 | 0 | 0 |
| *ssb1* | 5.0 | 486 | 15.2 | 0.02 | 33.7 | 49.6 | 0 | 0 | 0 | 0 | 0 | 49.6 | 83.3 | 88.7 |
| *sse1* | 0 | 588 | 53.4 | 46.6 | 0 | 0 | 0 | 0 | 0 | 0 | 0 | 0 | 0 | 0 |
| *sse1* | 5.0 | 630 | 21.4 | 3.2 | 33.8 | 41.6 | 0 | 0 | 0 | 0 | 0 | 41.6 | 75.4 | 74.4 |
| *sse2* | 0 | 483 | 38.9 | 61.1 | 0 | 0 | 0 | 0 | 0 | 0 | 0 | 0 | 0 | 0 |
| *sse2* | 5.0 | 202 | 38.6 | 5.9 | 42.6 | 0 | 4 | 0 | 5.4 | 3.5 | 0 | 9.4 | 55.5 | 16.9 |
| *ssz1*[5] | 0 | 528 | 59.8 | 40.1 | 0 | 0 | 0 | 0 | 0 | 0 | 0 | 0 | 0 | 0 |
| *ssz1*[5] | 5.0 | 668 | 24.2 | 3.3 | 34.4 | 6.02 | 0 | 0 | 0 | 0 | 0 | 6.02 | 40.4 | 10.8 |

[1]Wild type 1 is EY3136 *ssa1 ssa2* is EY3141, wild type 2 is S288c BY4741, and *ydj1*, *fes1*, *snl1*, *sti1*, *ssb1*, *sse1*, *sse2*, *ssz1* are isogenic null mutant derivatives of BY4741 (Table S1). As photos were done live, separate experiments were done for different α factor concentrations and only 2 strains were done at a time with staggered α factor addition times to permit timely photographing. Wild type1 and *ssa1 ssa2* strains were grown streaked on YPD plates at 30°C then grown overnight in liquid culture at 30°C, diluted in fresh prewarmed liquid culture and grown to an $A_{600}$ of ~0.5 then shaken with α factor or peptide buffer before photographing and/or tallying live. Wild type 2 and the other mutants strains were freshly grown overnight in thin streakouts on YPD plates at 30°C, then inoculated into liquid YPD to an $A_{600}$ of 0.2 then shaken at 30°C for 2 hours, centrifuged and resuspended at $A_{600}$ of 0.4 in prewarmed YPD medium with or without α factor or peptide buffer then incubated at 30°C with shaking for two hours. Cells were sonicated prior to being photographed.
[2]N is the number of cells counted, %RUB is round unbudded cells, %B is budded cells, %EI is enlarged irregular or football

ellipse shaped unbudded cells, %B is budded cells, %S is shmoos, % LS is shmoos with long projection, %MS is shmoos with two projections (multi-shmoo tip), %CS is shmoo with crooked projection, %DS is two enlarged shmoos joined at middle, %BB enlarged budded cells. %TS is % total shmoos, unbudded and budded cells with projections (%S+%LS+%MS+%CS+%DS). UCE is total unbudded cell enlargement (%EI+%S+%LS+%MS+%CS). % WT S is the % total unbudded shmoos divided by the % total unbudded shmoos for the wild type strain, a normalized value.

[3]It was difficult to identify many true shmoos with typical elongated projections for *ssa1 ssa2* mutants compared to wild type 1. The cells that were classified as shmoos include triangular cells with small projections, most cells were round and enlarged. After longterm treatment with 5 mM $\alpha$ factor, 7.9 +/ 1.8 S.E. percent of wild type 1 cells and 3.2 +/- 0 S.E. of *ssa1 ssa2* cells had a crumpled morphology typical of cells that have died by lysis or yeast “apoptosis”.

[4]*fes1* cells treated with a factor have novel cell morphology phenotypes of cigar shaped unbudded cells and shmoos with a bud on the side of the cell rather than at the projection tip as shown in Table S8.

[5]*ssz1* cells had a flocculent phenotype and were very clumpy even after sonication.
