## Supplementary material for "Effects of HSP70 chaperones Ssa1 and Ssa2 on Ste5 scaffold and the mating mitogen-activated protein kinase (MAPK) Pathway in *Saccharomyces cerevisiae*": TableS8

Table S8. Morphology of wild type and *fes1* mutant strains before and after exposure to α factor

| | Mean Percentage of Total Cells +/- S.E.[3] | | | |
|---|---|---|---|---|
| Morphology[1] | WT –αF | WT + αF | fes1 –αF | fes1 +αF |
| <u>1.Tally 1</u> | | | | |
| Unbudded | 43.2 ± 1.5 | 25.1 ± 0.9 | 48.4 ± 2.7 | 26.5 ± 0.7 |
| Budded | 55.5 ± 2.3 | 0.6 ± 0.2 | 51.1 ± 1.1 | 11.9 ± 3.0 |
| Cigar unbudded | 0 | 7.1 ± 4.6 | 0.5 ± 0.4 | 1.2 ± 0.7 |
| Shmoo/long shmoo | 0 | 41.7 ± 2.7 | 0 | 30.0 ± 4.4 |
| Bent shmoo | 0 | 9.6 ± 2.7 | 0 | 1.3 ± 0.8 |
| Irregular unbudded (Almost shmoo) | 0.8 ± 0.4 | 13.6 ± 9.5 | 0 | 25.6 ± 7.6 |
| Multi-shmoo | 0.5 ± 0.3 | 1.8 ± 0.3 | 0 | 0.4 ± 0.7 |
| Budding shmoo | 0 | 0.5 ± 0.5 | 0 | 0.8 ± 0.4 |
| Shmoo with side bud[2] | 0 | 0 | 0 | 15.0 +/- 2.0 |
| <u>2.Tally 2</u> | | | | |
| Short shmoo | | 27.5 ± 3.0 | | 18.2 ± 4.0 |
| Medium shmoo | | 45.7 ± 5.6 | | 19.4 ± 5.4 |
| Long shmoo | | 14.1 ± 0.6 | | 1.3 ± 0.7 |
| Total shmoo | | 87.4 ± 9.1 | | 38.9 ± 9.7 |

[1]Strains EYL1740 (S288c BY4741) and a *fes1* derivative of BY4741 were grown in YEPD to early logarithmic phase at room temperature (A600 of ~0.2), samples were adjusted to equal density and grown to an A600 of 0.6 then equal A600 units of cells were pelleted and resuspended in fresh YEPD with or without 2.5 μM α factor and incubated at room temperature with shaking. Samples were then fixed with 1/10 volume of 37% formaldehyde and photographed on the microscope. Tallies for cell morphology were done from the images. Approximately 350-600 cells were counted per sample.

[2]A shmoo with a round bud on its side is a novel phenotype only detected in the fes1D strain..

[3]S. D. is standard deviattion and S. E. is the standard error $\frac{\text{S.D.}}{\sqrt{N}}$ where $\text{N} = 3$ is the sample size.
