## SupportingInformationText for "Effects of HSP70 chaperones Ssa1 and Ssa2 on Ste5 scaffold and the mating mitogen-activated protein kinase (MAPK) Pathway in *Saccharomyces cerevisiae*"

### Supplemental Methods

#### Whole cell extract and pull-down methods in Fig 2

The large-scale co-immunoprecipitations of Ste5-Myc9 for mass spectrometry were done in 125 mM potassium acetate and no NaCl, whereas the co-immunoprecipitations of Ste5-Myc9 and HA3-Ste5 with Ssa1 and signaling partners were done with 250 mM NaCl and no potassium acetate. In Fig 1B, cells expressing Ste5-Myc9 were grown in SC-selective medium containing 2% dextrose to logarithmic phase to an A600 of 1.0 with or without treatment with 50 nM $\alpha$ factor for 2 hrs. Cells were pelleted, washed and immediately broken using liquid nitrogen and grinding with a mortar and pestle to break the cells, keeping samples in an ice bath in a cold room. Protein concentration was determined by the Biorad method and a portion of the samples were run on SDS-PAGE gels and the other portion was frozen in dry ice-ethanol and stored at $-80^{o}C$. The extraction and immunoprecipitation (IP) buffers contained 20 mM Tris-HCl pH 7.2, 125 mM potassium acetate, 0.5 mM EGTA, 0.5 mM EDTA, 1 mM DTT, 0.1% Tween 20, 12.5% Glycerol (vol/vol) 1 mM 4-(2-aminoethyl)-benzenesulfonylfluoride, 1 mM sodium ortho-vanadate, 25 mM β-glycerophosphate, 10 μg/ml pepstatin A.

In Fig 2, cells expressing Ssa1-GFP and either Ste5-Myc9 or HA3-Ste5 were grown in SC-selective medium containing 2% dextrose to mid-logarithmic phase of A600 of 0.8-1.0 and induced with $\alpha$ factor for 15 minutes (where indicated, 50 nM $\alpha$ factor for *bar1D/sst1D* cells, 5 $\mu$M for *BAR1/SST1* cells), pelleted, washed, frozen in an dry ice/ethanol bath, then thawed on ice and broken with six 30-s pulses with glass beads using a vortex after resuspending cells in modified-H buffer containing 25 mM Tris-HCl pH 7.4; 10% glycerol (vol/vol); 250 mM NaCl; 15 mM EGTA; 0.1% Triton-X-100; 1 mM sodium azide; 0.25 mM each meta- and orthovanadate; 1 µg/ml PMSF; 2 mM benzamidine; 4 mM 1,10-phenanthroline; 50 mM NaF; 5 µg/ml each of pepstatin A, leupeptin, peptin and chymostatin. Lysates were clarified with a 10 min centrifugation at 3,000 RPM to remove cellular debris followed by centrifugation for 10 min at 10,000 RPM at 4$^{o}$C. Protein concentration was determined by the BioRad method and samples were frozen in dry ice ethanol bath and stored at -80$^{o}$C. IPs were done with similar buffer with 200 mM NaCl and 7% glycerol.

**Fig S6. Representative fields of wild type and Hsp70 mutants treated with 2.5 $\mu$M $\alpha$ factor.** Cells were grown at room temperature to logarithmic phase and treated with 2.5 $\mu$M $\alpha$F for 90 minutes. A. WT EYL1740 (S288c-BY4741), B. *fes1*, C. *ydj1*. See Table S7 and Table S8 for tallies of morphology.

**Fig S7. Representative fields of wild type and hsp70 mutants treated with 5 $\mu$M $\alpha$F.** Cells were grown at room temperature to logarithmic phase and treated with 5 $\mu$M $\alpha$F for 90 minutes. A. WT EYL1740 (S288c-BY4741), B. *ydj1*, C. *ssb1*, D. *sse1*, E. *sse2*, F. *ssz1*, G. *sti1*. See Table S8 for tallies of morphology.

S10. Certus Technology methods to use N.I.H. ImageJ for densitometry measurements. ImageJ|Sybil https://www.sybil-fp7.eu/node/95.

S11. Lancaster AK, Nutter-Upham A, Lindquist S and King OD. PLAAC: a web and command-line application to identify proteins with Prion-Like Amino Acid Composition Bioinformatics 2014; doi:10.1093/bioinformatics/btu310.

S12. Alberti S, Halfmann R, King O, Kapila A and Lindquist S. A systematic survey identifies prions and illuminates sequence features of prionogenic proteins Cell 2009; 137:146–158.

S13. Prilusky J, Felder CE, Zeev-Ben_mordehai T, Rydberg EH, Mann O, Beckmann JS, Silman I and Sussman JL. FoldIndex: a simple tool to predict whether a given protein sequence is intrinsically unfolded. Bioinformatics 2005; 21: 3435–3438.

S14. Berman HM, Westbrook J, Feng Z, Gilliland G, Bhat TN, Weissig H, Shindyalov IN, Bourne PE. The Protein Data Bank. Nucleic Acids Research 2000; 28: 235-242. doi:10.1093/nar/28.1.235 URL rcsb.org

S15. Ward JJ, Sodhi JS, McGuffin LJ, Buxton BF and Jones DT. Prediction and functional analysis of native disorder in proteins from the three kingdoms of life. J. Mol. Biol. 2004; 337: 635-645. DISOPRED2, DISOPRED3 server at http://bioinfDOTcsDOTuclDOTacDOTuk/psipred/

S16. Dosztányi Z, CsizmokV, Tompa P and Simon I. IUPred: web server for the prediction of intrinsically unstructured regions of proteins based on estimated energy content Bioinformatics Applications Note 2005. 21: 3433–3434.

S17. Dosztanyi Z. Prediction of protein disorder based on IUPred. Protein Science 2017; 27: 331-340.

S18. Good M, Tang G, Singleton J, Remenyi A and Lim, WA. The Ste5 scaffold directs mating signaling by catalytically unlocking the Fus3 MAP kinase for activation. Cell 2009; 136: 1085–1097.

S19. Zalatan JG, Coyle SM, Rajan S, Sidhu SS, and Lim WA. Conformational control of the Ste5 scaffold protein insulates against MAP kinase misactivation. Science 2012; 337: 1218–1222.

S20. Wang J, Youkharibache P, Marchler-Bauer A, Lanczyckj C, Zhang D *et al*. iCn3D: From Web-Based 3D Viewer to Structural Analysis Tool in Batch Mode. Front. Mol. Biosci. 2022 9: 831740.

S21. Slabinski, L, Jaroszewski, L., Rychlewski, L, Wilson, IA, Lesley, SA and Godzik, A. XtalPred: a web server for prediction of protein crystallization. Bioninformatics 2007; 23: 3403-3405. XTalPred-RF server at.http://ffasDOTburnhamDOTorg/XtalPred.

S22. Schneider M, Rosam M, Glaser M, Patronov A, Shah H, Back C, Daake A, Buchner J and Antes I. BiPPred: Combined sequence- and structure-based prediction of peptide binding to the Hsp70 chaperone BiP. Proteins Structure Function Bioinformatics. 2016; 84:1390-1407.

S23. Rose GD, Geselowitz AR, Lesser GJ, Lee RH, and Zehfus MH. Hydrophobicity of amino acid residues in globular proteins. Science. 1985;229:834-838.

S24. Zimmerman JM, Eliezer N and Simha R. The characterization of amino acid sequences in proteins by statistical methods. Journal of Theoretical Biology 1968; 21:170-201.

S25. Jahandideh S, Jaroszewski L and Godzik A. Improving the chances of successful protein str5cture determination with a random forest classifier 2014; Acta Crystallog.r D. Biol. Crystallogr. 70(Pt 3),627-663.

S26. Dosztányi Z, CsizmokV, Tompa P and Simon I. IUPred: web server for the prediction of intrinsically unstructured regions of proteins based on estimated energy content Bioinformatics Applications Note 2005. 21: 3433–3434.

S27. Dosztanyi Z. Prediction of protein disorder based on IUPred. Protein Science 2017; 27: 331-340.

S28. Meszaros B, Erdos G and Dosztanyi Z. IUPred2A: context-dependent prediction of protein disorder as a function of redox state and protein binding. Nucl. Acids Res 2018; 46: W329-W337.

S29. Gasteiger E, Hoogland C, Gattiker A, Duvaud S, Wilkins MR, Appel RD and Bairoch A. Protein Identification and Analysis Tools on the ExPASy Server (In) John M. Walker (ed): The Proteomics Protocols Handbook, Humana Press 2005; pp. 571-607 web.expasy.org

S30. Gong Y, Kakihara Y, Krogan Y, Greenblatt J, Emili E, Zhang Z= and Houry WA. An atlas of chaperone–protein interactions in *Saccharomyces cerevisiae*: implications to protein folding pathways in the cell Molecular Systems Biology 2009; 5: 275-289.

S31. Truman AW, Kristjansdottir K, Wolfgeher D, Hasin N, Polier S and Zhang H. CDK-Dependent Hsp70 Phosphorylation Controls G1 Cyclin Abundance and Cell-Cycle Progression. Cell 2012; 151: 1308–1318.

S32. Cherry J, Hong EL, Amundsen C, Balakrishnan R, Binkley G, Chan ET, Christie KR, Constanzo MC *et al.* Saccharomyces Genome Database: the genomics resource of budding yeast. Nucleic Acids Res.2012; 40: D700–D705.

S33. Orchard, S. Ammari, M., Aranda, B., Breuza, L., Briganti, L., Broackes-Carter, F. et al. The MI project—IntAct as a common curation platform for 11 molecular interaction databases. Nucleic Acids Res. 2014; 42(D1): D358-D363.

S34. Babu, M, Vlasblom, J, Pu, S., Guo, X, Graham, C, Bean, BDM, Burston, H E, Vizeacoumar, FJ, Snider, J, Phanse, S, Fong, V, Yi, Y, Tam, C, Davey, M, Hnatshak, O, Bajaj, N, Chandran, S, Punna, T., Christopolous, C, Wong, V, Yu, A, Zhong, G, Li, J, Stagljar, I, Conibear, E, Wodak, SJ, Emili, A., Greenblatt, JF. Interaction landscape of membrane-protein complexes in *Saccharomyces cerevisiae* Nature 2009; 489: 585-590.

S35. Willmund, F, del Alamo, M, Pechmann, S, Chen,T, Albanese,V, Dammer, EB, Peng, J, and Frydman, J. The Cotranslational Function of Ribosome-Associated Hsp70 in Eukaryotic Protein Homeostasis. Cell 2013; 152: 196–209.

S36. Jamuczak AF, Evers CE, Schwartz J-M, Grant CM and Hubbard SJ. Quantitative proteomics and network analysis of SSA1 and SSB1 deletion mutants reveals robustness of chaperone HSP70 network in Saccharomyces cerevisiae. Proteomics. 2015; 15: 3126-3139.

S38. Orchard S, Ammari M, Aranda B, Breuza L, Briganti L, Broackes-Carter F, *et al.* The MI project—IntAct as a common curation platform for 11 molecular interaction databases. Nucleic Acids Res. 2014; 42(D1): D358-D363.

S39. Wiederhold E, Veenhoff LM, Poolman B, and Slotboom J. Proteomics of *Saccharomyces cerevisiae* organelles. Mol. And Cell. Proteomics. 2010; 9.3: 431-445.
